# RNA-mediated double-strand break repair in human cells

**DOI:** 10.1101/2022.11.01.514688

**Authors:** Youngkyu Jeon, Margherita Maria Ferrari, Tejasvi Channagiri, Penghao Xu, Sathya Balachander, Vivian S. Park, Stefania Marsili, Zachary F. Pursell, Nataša Jonoska, Francesca Storici

**Author notes:** these authors contributed equally (Margherita Maria Ferrari, Tejasvi Channagiri, Penghao Xu).

## Abstract

Double-strand breaks (DSBs) in DNA are challenging lesions to repair. Human cells employ at least three DSB repair mechanisms, with a preference for non-homologous end joining (NHEJ) over homologous recombination (HR) and microhomology-mediated end joining (MMEJ)^1,2^. In contrast to HR, NHEJ and MMEJ do not utilize a DNA template molecule to recover damaged and/or lost nucleotides^2^. NHEJ directly ligates broken DNA ends, while MMEJ exploits the alignment of short microhomologies on the DSB sides and is associated with deletions of the sequence between the microhomologies^3,4^. It is unknown whether and to what extent a transcript RNA has a direct role in DSB-repair mechanisms in mammalian cells. Here, we show that both coding and non-coding transcript RNA facilitates DSB repair in a sequence-specific manner in human cells. Depending on its sequence complementarity with the broken DNA ends, the transcript RNA could promote the repair of a DSB or gap in its DNA gene via NHEJ or MMEJ, or mediate RNA-templated repair. The transcript RNA influences DSB repair by NHEJ and MMEJ even when the transcription level is low. The results demonstrate an unexpected role of transcript RNA in directing the way DSBs are repaired in human cells and maintaining genome stability.

---

RNA, transcribed from DNA as a complementary single strand copy, is a multifunctional nucleic acid that has a primary role of messenger RNA (mRNA) to convert the information stored in DNA into proteins via translation. More than 90% of the human genome is transcribed into RNA^5^. Of this, the fraction transcribed into mRNA represents ∼1%; all the rest is transcribed into non-coding RNA^6–8^. Yet, beyond its significant role in the regulation of gene expression, little is known about non-coding RNA functions^9^, and it is unknown whether both coding and non-coding RNA may have a more prevalent role in DNA repair and genome integrity than anticipated.

In budding yeast, an endogenous RNA transcript can be used as a direct homologous template for accurate DSB repair of its DNA gene in *cis* via RNA-templated DSB repair (R-TDR)^10,11^. R-TDR is blocked by ribonucleases (RNases) H1 and H2, which cleave RNA in a hybrid with DNA^10,11^. Nevertheless, such RNase H inhibition of R-TDR highlights the marked capacity of RNA to form RNA/DNA hybrids at the site of a DSB. Several studies have identified different, non-templating roles of transcript RNA in promoting DSB repair of transcribed DNA by HR in mammalian cells^12–14^. Beyond HR, NHEJ-related proteins have been found to form a multiprotein complex with RNA polymerase II and to be associated with transcribed genes after inducing a DSB in these DNA loci, suggesting that RNA may help error-free NHEJ in human cells^15^. The chromatin context was shown to have an impact on DNA DSB repair induced by the clustered regularly interspaced short palindromic repeats (CRISPR) associated protein 9 (Cas9). In fact, it was recently found that NHEJ is broadly biased toward euchromatin in human cells, while MMEJ has a higher contribution in the heterochromatin context^16^. Furthermore, there is mounting evidence that DNA:RNA hybrids form at DNA DSBs from pre-transcribed RNA in transcriptionally active loci^17^. However, it remains unknown whether RNA has the capacity to promote DSB repair directly as a template, participate in the DSB repair processes, and affect the frequency of the DSB repair products in a sequence-dependent manner in mammalian cells.

To unravel the role of a transcript RNA in DSB repair in human cells, we developed an assay employing various RNA transcripts that differ by sequence and transcription level and are generated from a constitutively transcribed gene, in which we induced one or two DSBs by the Cas9 endonuclease. We engineered a plasmid to contain the *DsRed* gene with two exons (Exon1 and Exon2) and an artificial intron (Intron) in between. The *DsRed* gene was transcribed from the strong constitutive CMV promoter, and carried the SV40 origin for DNA replication in the human embryonic kidney cells expressing the T antigen (HEK-293T). This construct is called Sense (**Figure 1a** and **Extended Data Figure 1**). We made another construct, called BranchΔ, in which we deleted the branch region of the intron to prevent splicing of the intron. We made a third construct, called pCMVΔ, in which we removed the CMV promoter to minimize transcription of the *DsRed* gene while still allowing intron splicing (**Figure 1a** and **Extended Data Figures 1 and 2a, b**). In all three constructs, we induced a DSB in the *DsRed* gene on either side of the intron, or simultaneously on both sides using two synthetic guide RNAs (sgRNAs) with Cas9 endonuclease. The sgRNA A binds near the junction of the 5′ exon, Exon1, and the intron. The sgRNA B binds near the junction of the intron and the 3′ exon, Exon2 (**Figure 1a**). We verified that Cas9 with sgRNA A, sgRNA B, or both sgRNAs A and B generates DSBs at predesigned sites in the constructs carrying the intact intron (Sense and pCMVΔ) as well as in the construct with branch deletion (BranchΔ) with similar efficiency. This was evaluated by performing in vitro cleavage of two PCR products of the *DsRed* gene, one containing the branch region of the intron like in the Sense and pCMVΔ and one lacking it like in the BranchΔ, by Cas9 and sgRNA A and/or sgRNA B. The results showed that Cas9 with sgRNA A and/or sgRNA B cleaved the two different DNA substrates equally well (**Extended Data Figure 3a**). Individually, the plasmids with the Sense, BranchΔ, and pCMVΔ constructs were transfected four independent times each into the HEK-293T wild-type cells as well as HEK-293T knock-out cells having mutations in the catalytic subunit of RNase H2 (RNase H2A KO) (**Extended Data Figure 4**). In each *DsRed* construct, we transfected a plasmid expressing Cas9 and a plasmid producing sgRNA A or sgRNA B to generate 1 DSB, or both plasmids for sgRNA A and B to generate a double-strand gap (2 DSBs) (**Figure 1a, b**). As a no-DSB control, each plasmid was also co-transfected in the wild-type or RNase H2A KO cells with sgRNA A and B, but without Cas9. After a few days, the plasmid DNAs were extracted from the cells and prepared for next generation sequencing (NGS) to study the sequence around the DSB site or the gap (see Methods, and **Supplementary Tables 1 and 2**). The sequences of the reads from each sample library were then analyzed for specific signatures to categorize them as either DSB repair by NHEJ (small in/dels), MMEJ (deletions between microhomologies), or R-TDR (identical sequence to that of the spliced RNA transcript) (**Figure 1b**). Then the frequency of each of these three repair mechanisms was calculated (see Methods).

**Figure 1.**
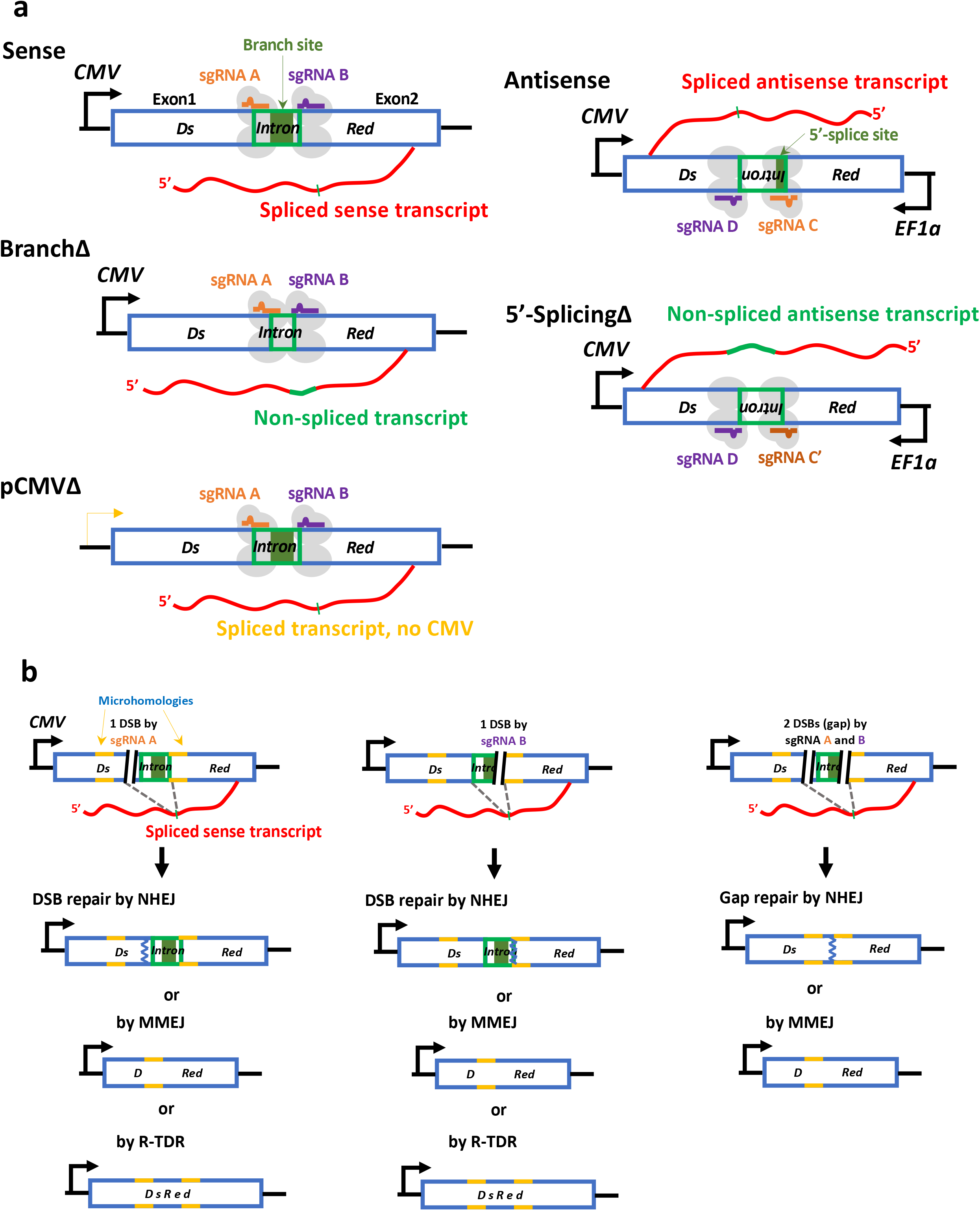
Assay to study RNA-mediated DSB repair in human cells. **a**, Schemes of sense and antisense-genetic constructs expressing RNA transcripts that differ by sequence and transcription level. (left) Sense constructs: Sense, BranchΔ, and pCMVΔ. (right) Antisense constructs: Antisense and 5′-SplicingΔ. All these constructs contain the *DsRed* gene (blue-framed box) with an intron (green-framed box) in the sense or antisense orientation, respectively. The sense and antisense transcript RNAs (in red) are depicted after intron splicing (thin green mark) or carrying the intron (thick green line) for the splicing-mutant constructs. The single-guide RNA (sgRNA) A (orange line) and sgRNA B (purple line) with Cas9 endonuclease (light gray ovals) bind to the complementary DNA of the sense constructs to generate a DSB. The sgRNA C (orange line) or C’ (dark orange line) and D (purple line) with Cas9 endonuclease bind to the complementary DNA of the antisense constructs to generate a DSB. Black arrows: *CMV*, cytomegalovirus promoter, or *EF1α*, human eukaryotic translation elongation factor 1 alpha promoter. Transcription activity by a cryptic promoter is indicated with a yellow arrow. Dark green box: branch site of the intron or 5′-splice site for the sense and antisense constructs, respectively. **b**, Scheme of DNA products for the Sense construct obtained following DSB repair by different DSB repair mechanisms. NHEJ, non-homologous end joining; MMEJ, microhomology-mediated end joining; R-TDR, RNA-templated DSB repair; DSB (black parallel lines); an example of a microhomology pair (yellow lines); DNA repaired by NHEJ (blue zigzag line).

## The sequence of a transcript RNA guides DSB repair in its DNA by NHEJ

We first examined the sequencing data obtained from the induction of one DSB by sgRNA A or B in the Sense, BranchΔ, and pCMVΔ constructs transfected in the RNase H2 wild-type and RNase H2A KO HEK-293T cells. As expected, the most prevalent mechanism of DSB repair in all constructs was NHEJ. While it was not possible to distinguish the sequence of error-free repair by NHEJ from that of the uncut constructs, we searched for in/dels near the DSB site as the signature for NHEJ (see Methods). The in/del signature was practically absent in the no-DSB controls, as well as in the control sequences 30 nucleotides distant from the DSB site (**Extended Data Figure 5a, b**). The analysis of the sequencing data revealed that the construct without splicing, BranchΔ, had higher frequency of NHEJ in/dels than the constructs with splicing (Sense and pCMVΔ) in wild-type and more evidently in the RNase H2A KO cells (**Figure 2a** and **Supplementary Table 3**). The frequencies of error-free NHEJ/uncut constructs did not compensate for such difference in the NHEJ in/del frequency between the non-splicing and the splicing constructs (**Extended Data Figure 6a, b**). We then performed an analysis of the NHEJ in/del variation observed in the samples. Using the NGS data, we aligned the sequence reads to an error-free end-joining reference sequence, extracted the 20 nucleotides of the alignment around the DSB site (called the *DSB-sequence window*), and determined the type and number of variations of each read sequence compared to the reference sequence to generate *variation-distance graphs* for each of the constructs (see Methods). The results showed that the three constructs displayed a very similar pattern of in/dels (**Extended Data Figures 7** and **8a**, and **Figure 2b**), suggesting a similar mechanism of DSB repair. Notably, when we compared Sense vs. BranchΔ, the comparison graphs revealed that the most prevalent in/dels were insertions in sgRNA A and deletions in sgRNA B, but these in/dels usually had higher frequency for the BranchΔ construct (**Figure 2c**). Moreover, in RNase H2A KO cells, the majority of in/dels had higher frequency for the BranchΔ compared to the Sense construct (**Figure 2c** and **Extended Data Figures 8b, d**). Interestingly, when comparing the NHEJ data of the Sense vs. the pCMVΔ construct, we identified a higher frequency of in/dels for Sense than pCMVΔ following a DSB by sgRNA B. Overall, these results provide evidence that the sequence of a transcript RNA, its complementarity with the DSB ends, and the level of transcription directly influence the efficiency of DSB repair by NHEJ and the frequency of the repaired products by NHEJ.

**Figure 2.**
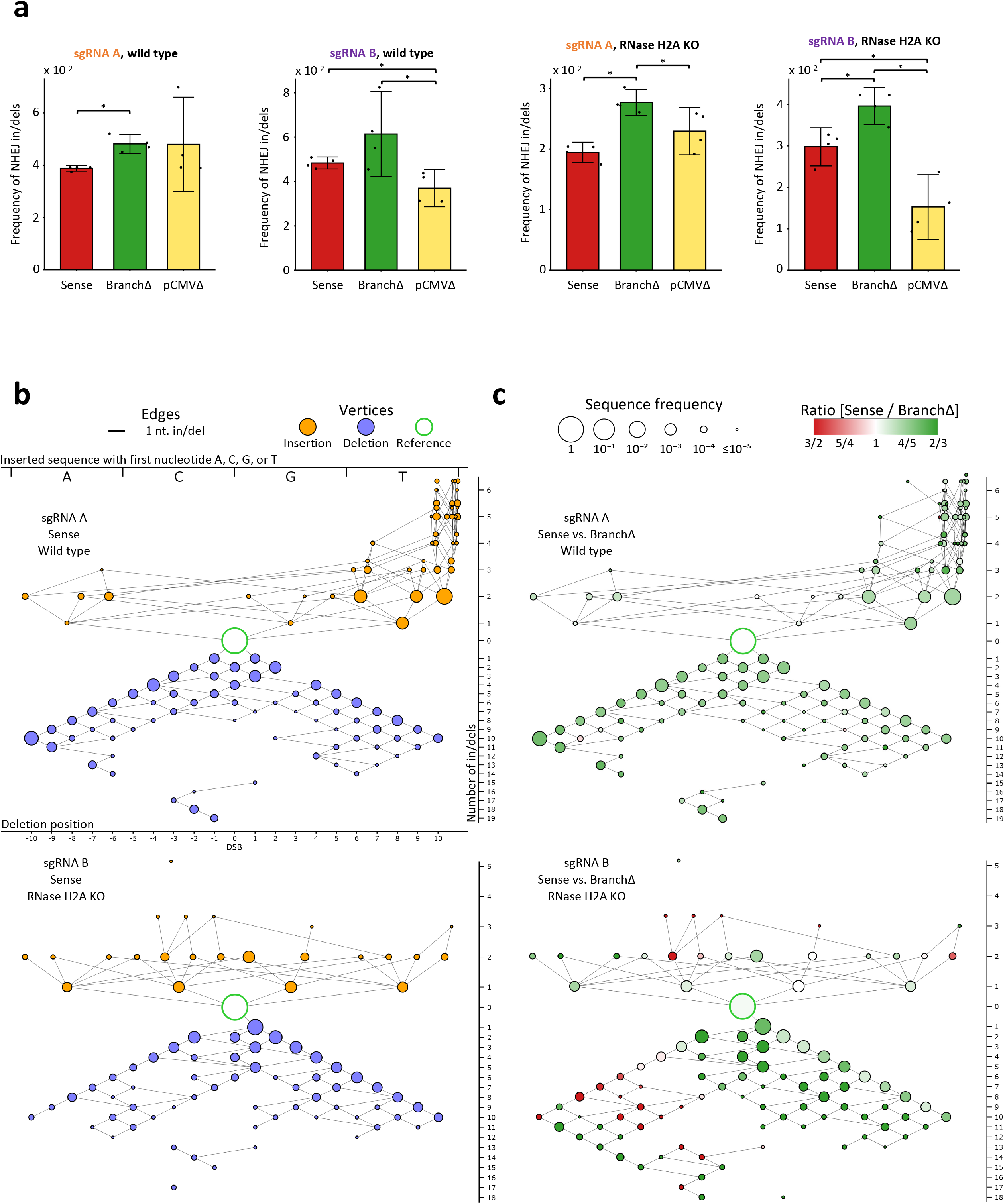
Transcript RNA promotes DSB repair by NHEJ in a sequence-dependent manner. **a**, Frequencies of NHEJ in/dels observed following a DSB by the sgRNA A or B in the Sense (red), BranchΔ (green), and pCMVΔ (yellow) constructs of wild-type (left) and RNase H2A KO (right) cells. Plotted data are mean ± s.d. with 4 biological replicates indicated with dots; N=4. \**P*-value = 0.029 (two-tailed Mann-Whitney *U* test). **b**, Individual variation-distance graphs illustrating sequence variations within DSB-sequence windows obtained after DSB induction by sgRNA A (top) or sgRNA B (bottom) in the Sense construct of wild-type cells. Each vertex represents a single DSB-sequence window. An edge between two vertices indicates that the two corresponding DSB-sequence windows differ by a single nucleotide in/del. Insertion vertices (in orange) are placed above the reference vertex (in white with a green border), while deletion vertices (in blue) are placed below it. The vertex size shows the log of the mean frequency of the corresponding DSB-sequence window in the four repeats of the experiment. For insertions, the alphabetical order of the inserted sequences, from A on the left to T on the right, are indicated by the *x*-coordinate. Insertions of size 3 or more have vertices on multiple lines, staggered vertically to reduce overlap. The *x*-coordinate of deletions indicates the position of the first deleted nucleotide, from the most upstream (left-most) to the most downstream (right-most). The *y*-coordinate indicates the number of variations in the DSB-sequence windows, with higher variations placed further from the reference. See **Extended Data Fig. 6** for the variation-distance graph key. **c**, Comparison variation-distance graphs of the DSB-sequence windows obtained after DSB induction by sgRNA A (top) or sgRNA B (bottom) for the Sense vs. the BranchΔ constructs of wild-type cells. The vertices represent the same DSB-sequence windows as for the individual graphs while the vertex colors specify the relative frequency in the Sense (red) vs. the BranchΔ (green) construct; the vertex sizes show the log of the higher of the two mean frequencies of the corresponding DSB-sequence windows in the two compared constructs.

## The sequence of a transcript RNA guides DSB repair in its DNA by MMEJ

Seeking MMEJ-signature sequences and their frequencies, we analyzed the sequencing data obtained from DSB induction by Cas9 and sgRNA A or B in the Sense, BranchΔ, and pCMVΔ constructs transfected in the wild-type as well as in the RNase H2A KO HEK-293T cells. We identified all possible microhomology pairs of three base pairs or longer in the *DsRed* exon and intron sequences in common between the splicing and no-splicing constructs (**Extended Data Figures 9a-c**). We then determined the deletion signature for each microhomology pair and identified the signatures that were detected in all the NGS libraries of the Sense, BranchΔ, and pCMVΔ constructs for the repair of the DSB generated by sgRNA A or B in the wild-type or RNase H2A KO HEK-293T cells. The results of DSB repair by MMEJ for the exon-exon microhomology pairs (those between the two exons: Exon1-Exon2 for cleavage by sgRNA A and Exon2-Exon1 for cleavage by sgRNA B) were opposite to those of exon-intron microhomology pairs (those between an exon and intron: Exon1-Intron for cleavage by sgRNA A, and Exon2-Intron for cleavage by sgRNA B). The exon-exon MMEJ had higher frequency of DSB repair for the constructs with splicing (Sense and pCMVΔ) than for the construct without (BranchΔ) (**Figure 3a** and **Extended Data Figure 9d**), while the exon-intron MMEJ had higher frequency of DSB repair in the construct without splicing than for those with splicing (**Figure 3b** and **Extended Data Figure 9d**). Next, for each microhomology pair, we calculated the ratio of the mean frequency in the Sense construct to the mean frequency in the BranchΔ construct. As shown in **Figure 3c**, the ratio Sense/BranchΔ for the exon-exon microhomologies was mostly higher than 1, while the ratio Sense/BranchΔ for the exon-intron microhomology pairs was mostly lower than 1. Also, the ratio of total exon-exon frequencies to total exon-intron frequencies for MMEJ within the Sense and pCMVΔ libraries was significantly higher than that obtained for the BranchΔ libraries suggesting more efficient MMEJ between exon sequences for the constructs with splicing compared to the construct without splicing (**Extended Data Figure 9e**). These results support the role of spliced RNA in promoting MMEJ between exon-exon microhomologies, and the role of the non-spliced RNA in promoting MMEJ between exon-intron microhomologies. Notably, the results for DSB repair by NHEJ (**Figure 2a**) parallel those of DSB repair by exon-intron microhomologies (higher frequency in constructs with spliced RNA than those with non-spliced RNA) but are opposite to those of DSB repair by exon-exon microhomologies (**Figure 3a, b**). Also, the results obtained from the RNase H2A KO cells were stronger than those obtained from the wild-type HEK-293T cells, especially for repair of the DSB caused by sgRNA B (**Figure 3** and **Supplementary Table 3**), which suggests that the interactions of the transcript RNA with the broken DNA ends have greater stability when RNase H2 is not functional. Altogether, these findings highlight the capacity of RNA to promote DSB repair via MMEJ in a sequence-specific manner.

**Figure 3.**
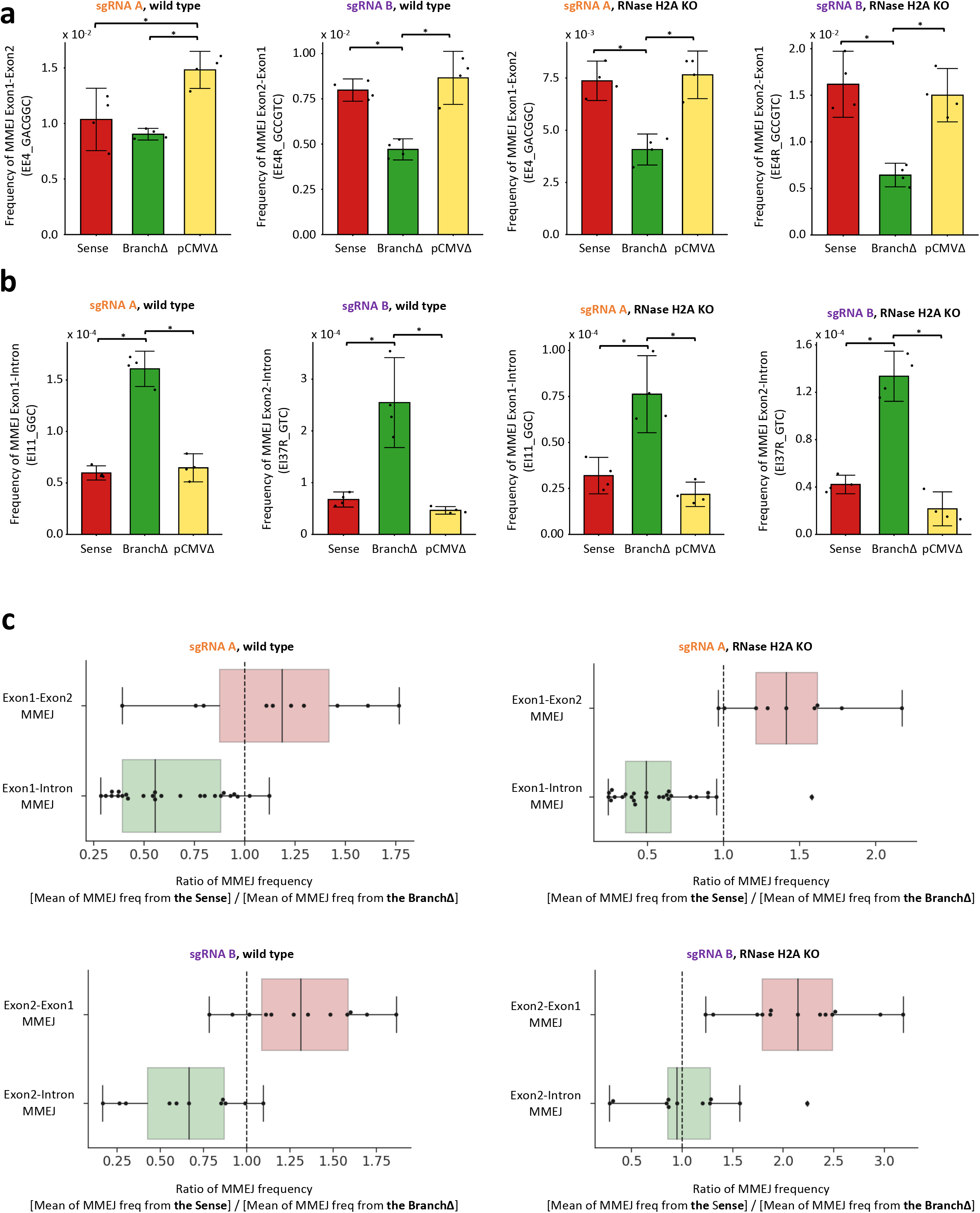
Transcript RNA promotes DSB repair by MMEJ in a sequence-dependent manner. **a**, MMEJ frequencies from a 6-bp microhomology pair as an example of an exon-exon MMEJ following a DSB by the sgRNA A (EE4_GACGGC, see **Extended Data Figure 9b, d**) or B (EE4_GACGGC, see **Extended Data Figure 9c, d**) in the Sense (red), BranchΔ (green), and pCMVΔ (yellow) constructs of wild-type (left) and RNase H2A KO (right) cells. Data are mean ± s.d. of 4 biological replicates indicated with dots; N=4. **b**, MMEJ frequencies from a 3-bp microhomology pair as an example of exon-intron MMEJ following a DSB by the sgRNA A (EI11_GGC, see **Extended Data Figure 9b, d**) or B (EI37R_GTC, see **Extended Data Figure 9c, d**) in the Sense (red), BranchΔ (green), and pCMVΔ (yellow) constructs of wild-type (left) and RNase H2A KO (right) cells. \**P*-value = 0.029 (Two-tailed Mann-Whitney *U* test). **c**, Boxplot showing the ratios of MMEJ frequencies from exon-exon (red box) and exon-intron (green box) microhomologies between the Sense and the BranchΔ constructs of wild-type (left) and RNase H2A KO (right) cells after DSB by sgRNA A (top) or B (bottom). Each ratio was calculated by using an average of 4 repeats of MMEJ frequencies from each construct. Ten Exon1-Exon2 and 25 Exon1-Intron microhomologies (black dots) are shown for MMEJ following a DSB by the sgRNA A (top). 13 Exon2-Exon1 and eleven Exon2-Intron microhomologies are shown for MMEJ following a DSB by the sgRNA B (bottom). The median of the points is shown as the middle line of the box. The first and third quartiles are indicated by the box frames and the whiskers represent the largest point not more than 1.5 interquartile range (IQR) beyond the box frame. All data points outside the whiskers are classified as outliers and shown as diamond points.

## The sequence of a transcript RNA guides gap repair in its DNA by NHEJ and MMEJ

To investigate whether RNA mediates the repair of a double-strand gap in DNA, we generated a DSB on each side of the intron in the *DsRed* gene by simultaneously using sgRNA A and B to cut the Sense, BranchΔ, or pCMVΔ constructs (**Figure 1b**) in both wild-type and RNase H2A KO HEK-293T cells. The NGS data from the Sense, BranchΔ, and pCMVΔ constructs were analyzed to determine the frequency of NHEJ signatures with intron pop-out and MMEJ signatures with intron pop-out by recombination between exon-exon microhomologies. We then compared these NHEJ and MMEJ frequencies of intron pop-out with those obtained for the signatures of intron retention in the Sense, BranchΔ or pCMVΔ constructs. We found that the constructs with splicing (Sense and pCMVΔ) had a higher frequency of intron pop-out by NHEJ than the BranchΔ construct. The results were stronger in the RNase H2A KO cells, suggesting direct RNA-DNA interaction (**Figure 4a** and **Supplementary Table 3**). We then performed an analysis of the in/del variation observed in these samples that had gap repair by NHEJ. Using the NGS data, we aligned the sequence reads to an error-free end-joining reference sequence, extracted the 20 nucleotides of the alignment around the gap site, and determined the type and number of variations of each read sequence compared to the reference sequence to generate variation-distance graphs (see Methods). The results showed that the Sense, BranchΔ, and pCMVΔ constructs displayed a similar pattern of NHEJ in/dels (**Extended Data Figure 10a, c** and **Figure 4b**), suggesting a similar mechanism of DSB repair. However, the comparison graphs, showing the relative frequencies of in/dels in the Sense vs. the BranchΔ construct, revealed a higher frequency of both insertions and deletions by NHEJ in the Sense construct (**Figure 4b** and **Extended Data Figure 10b**). The comparison graphs of the Sense vs. the pCMVΔ construct showed higher frequencies of insertions and deletions for the pCMVΔ construct (**Extended Data Figure 10b**), as also revealed by results shown in **Figure 4a**. Results in RNase H2A KO cells corroborated those obtained in the RNase H2 wild-type cells (**Extended Data Figure 10c, d**). The pop-out of the intron from the *DsRed* gene following induction of the two DSBs near the intron-exon junctions was more frequent in the splicing constructs than in the non-splicing construct also for gap repair via MMEJ for exon-exon microhomologies (**Figure 4a, c, d** and **Extended Data Figure 11a, b, d**). The results were somewhat stronger in the RNase H2A KO compared to the wild-type cells (**Figure 4a, c, d, Extended Data Figure 11d**, and **Supplementary Table 3**).

**Figure 4.**
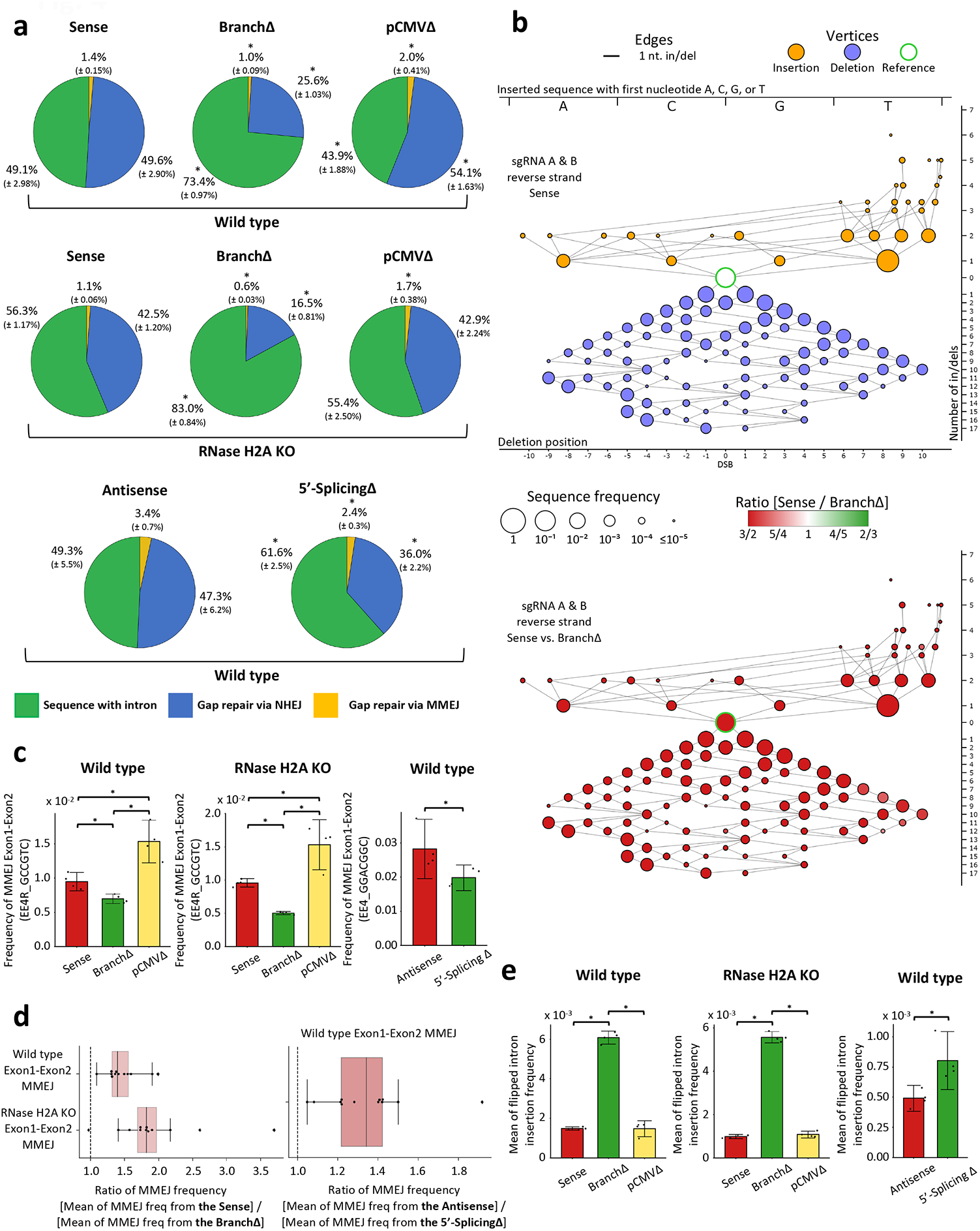
Transcript RNA promotes gap repair in a sequence-dependent manner via NHEJ and MMEJ. **a**, Pie charts showing frequencies of sequencing reads displaying intron retention or pop-out following a gap by the sgRNA A and B in the Sense, BranchΔ, and pCMVΔ constructs of wild-type (top) and RNase H2A KO (middle) cells, or gap by the sgRNA C/C’ and D in the Antisense and 5′-SplicingΔ constructs of wild-type cells (bottom). The fraction of sequences containing the intron is shown in green, the one derived from gap repair via NHEJ in blue, and the one derived from gap repair via MMEJ in yellow. Percentages represent an average of 4 repeats with standard deviation in parenthesis; N=4. *, *P* = 0.029 comparing frequencies of the BranchΔ or the pCMVΔ with those of the Sense construct, or comparing frequencies of the Antisense with those of the 5′-SplicingΔ construct via the two-tailed Mann-Whitney *U* test. **b**, (Top) individual variation-distance graph illustrating sequence variations within DSB-sequence windows obtained after gap induction by the sgRNA A and B in the Sense construct of wild-type cells. (Bottom) comparison variation-distance graph of the DSB-sequence windows obtained after gap induction by the sgRNA A and B in the Sense vs. the BranchΔ constructs of wild-type cells. Refer to **Figure 2** and **Extended Data Figure 7** for details. The sequences are reverse-complemented prior to computing *x-y* coordinates so that they correspond to the forward strand sequence coordinates. **c**, MMEJ frequencies from a 6-bp (EE4R_GCCGTC, sense constructs, see **Extended Data Figure 11b, d**) or 7-bp (EE4_GGACGGC, antisense constructs, see **Extended Data Figure 11c, e**) microhomology pairs as examples of exon-exon MMEJ, following a gap by (left) the sgRNA A and B in the Sense (red), BranchΔ (green), and pCMVΔ (yellow) constructs of wild-type and RNase H2A KO cells, and (right) the sgRNA C/C’ and D in the Antisense (red) and the 5′-SplicingΔ (green) constructs of wild-type cells. Data are mean ± s.d. with 4 biological replicates indicated with dots; N=4. *, *P* = 0.029 (two-tailed Mann-Whitney *U* test). **d**, Boxplot showing the ratios of MMEJ frequencies between the Sense and the BranchΔ constructs of wild-type and RNase H2A KO cells (left) and between the Antisense and the 5′-SplicingΔ constructs of wild-type cells (right) from Exon1-Exon2 microhomologies located upstream and downstream of the DNA gap. Each ratio was calculated by using an average of 4 repeats of MMEJ frequencies from each construct. Twelve Exon1-Exon2 microhomologies (black dots) are shown for MMEJ following the DNA gap in the sense constructs generated by the sgRNA A and B. 10 Exon1-Exon2 microhomologies (black dots) are shown for MMEJ following the DNA gap in the antisense constructs generated by the sgRNA C/C’ and D. The median of the points is shown as the middle line of the box. The first and third quartiles are indicated by the box frames and the whiskers represent the largest point not more than 1.5 interquartile range (IQR) beyond the box frame. All data points outside the whiskers are classified as outliers and shown as diamond points. **e**, Frequency of intron flipping caused by re-capture of the intron via NHEJ following a gap by (left) the sgRNA A and B in the Sense (red), BranchΔ (green), and pCMVΔ (yellow) constructs of wild-type and RNase H2A KO cells, and (right) the sgRNA C/C’ and D in the Antisense (red) and the 5′-SplicingΔ (green) constructs of wild-type cells. Data are mean ± s.d. with 4 biological replicates indicated with dots; N=4. *, *P* = 0.029 (two-tailed Mann-Whitney *U* test).

Opposite results were obtained for the insertion of the intron in the flipped orientation via NHEJ. The frequency of the flipped-intron insertion was a factor of three higher for the non-splicing than for the splicing constructs both in wild-type and RNase H2A KO cells (**Figure 4e**). These results, for the repair of the gap generated by two DSBs near the intron-exon junctions, point towards a role of the spliced RNA in promoting intron deletion, and, vice versa, a role of the non-spliced RNA in maintaining the intron in its original locus or in facilitating its flipping. Interestingly, the frequency of gap repair with intron flipping or deletion both by NHEJ and MMEJ was similar or higher in the pCMVΔ than the Sense construct, in both wild-type and RNase H2A KO cells (**Figure 4e, a, c**), suggesting that the level of transcription is not a limiting factor for transcript RNA to mediate DNA gap repair via NHEJ or MMEJ.

## A non-coding transcript RNA guides gap repair in its DNA by NHEJ and MMEJ

To understand whether the capacity of RNA to mediate DSB repair in human cells could occur also with non-coding RNA, we engineered a plasmid construct, called Antisense, in which the *DsRed* gene was also transcribed in the antisense orientation from the constitutive EF1a promoter, and the intron sequence was reversed to allow splicing from the antisense transcript of the *DsRed* gene (**Figure 1a**). We then engineered another plasmid, 5′-SplicingΔ, in which the 5′-splice site of the intron in the Antisense construct was deleted to block splicing (**Figure 1a** and **Extended Data Figure 1**). To investigate the capacity of the non-coding antisense RNA to mediate the repair of a gap in DNA, we generated a DSB on each side of the intron by simultaneously using sgRNA C and D to cut the Antisense construct, and C’ and D to cut the 5′-SplicingΔ construct (**Figure 1a** and **Extended Data Figure 1**). The respective sgRNAs cut the Antisense and 5′-SplicingΔ construct similarly in vitro using Cas9 (**Extended Data Figure 3b**). The Antisense and the 5′-SplicingΔ constructs were transfected into the wild-type HEK-293T cells together with the plasmid expressing Cas9 and plasmids for the respective sgRNAs. After a few days, the plasmid DNAs were extracted from the cells and prepared for NGS to study the sequence of the *DsRed* gene around the DNA gap. The NGS data from the Antisense and the 5′-SplicingΔ constructs were analyzed to determine the frequency of signatures of NHEJ with intron deletion, and MMEJ with intron deletion by recombination between exon-exon microhomologies. We then compared these NHEJ and MMEJ frequencies of intron deletion with those obtained for the signatures of intron retention in the Antisense and the 5′-SplicingΔ constructs. Like the results for the sense constructs, we found that the Antisense construct with splicing had a higher frequency of intron loss both by NHEJ and MMEJ than the 5′-SplicingΔ (**Figure 4a, c, d** and **Extended Data Figure 11a, c, e**). We then performed an analysis of the in/del variation observed in the samples that had gap repair by NHEJ. Using the NGS data, we aligned the sequence reads to an error-free end-joining reference sequence, extracted the 20 nucleotides of the alignment around the gap site, and determined the type and number of variations of each read sequence compared to the reference sequence to generate variation-distance graphs (see Methods). The results showed that the Antisense and the 5′-SplicingΔ constructs displayed a similar pattern of NHEJ in/dels (**Extended Data Figure 10e**), suggesting a similar mechanism of DSB repair. However, the comparison graphs, showing the relative frequencies of in/dels in the Antisense vs. the 5′-SplicingΔ construct, revealed a higher frequency of both insertions and deletions by NHEJ in the Antisense construct (**Extended Data Figure 10f**). The results were not as dominant for the Antisense construct as they were for the Sense construct (**Extended Data Figure 10d**, left graphs), likely due to the impact of the sense RNA transcribed from the CMV promoter of the Antisense and the 5′-SplicingΔ constructs.

Moreover, we examined the frequency of the flipped-intron insertion after 2-DSB induction in the Antisense and the 5′-SplicingΔ constructs in the wild-type HEK-293T cells. In line with results obtained for the sense constructs, we found that the flipped-intron insertion was more frequent in the 5′-SplicingΔ construct than in the Antisense construct (**Figure 4e**). This suggests that not just coding but also non-coding RNA has the capacity to promote gap repair by NHEJ or MMEJ in a sequence-dependent manner.

## RNA-templated DSB repair

To determine whether RNA could work as a template for DSB repair in human cells, we searched for the R-TDR signature within the NGS libraries of the Sense, BranchΔ, and pCMVΔ constructs after DSB induction by sgRNA A or B. In the 2-DSB system, the R-TDR signature can be confused with the sequences of repair by NHEJ. However, this is not the case for the 1-DSB systems. We found that in the 1-DSB systems the frequency of R-TDR was significantly higher for the Sense construct compared to the BranchΔ construct both in the wild-type and in the RNase H2A KO cells (**Figure 5**). We considered the possibility of reverse transcription of the RNA into a complementary DNA (cDNA) by LINE1 reverse transcriptase (RT) and successive DSB repair by cDNA. However, due to the capacity of LINE1 to use broken chromosomal DNA ends as a priming sequence for reverse transcription on the annealed RNA at the DSB site via the target-primed reverse transcription (TPRT) mechanism^18^, it is possible that the LINE1 RT utilizes the broken 3′-DNA DSB end as a primer for DNA synthesis on the annealed complementary RNA. Thus, this LINE1-mediated RT process would still represent a mechanism of DSB repair via R-TDR. We found the frequency of R-TDR repair to be higher in the Sense construct than the pCMVΔ construct, which contrasted with the results for NHEJ and MMEJ in which both constructs had similar frequencies of repair. This was particularly evident for the DSB system with sgRNA A both in the wild-type and in the RNase H2A KO cells (**Figure 5**). Moreover, the R-TDR of the DSB by sgRNA A was more efficient in RNase H2A KO cells than in the wild-type cells (**Figure 5** and **Supplementary Table 3**, *P* = 0.008).

**Figure 5.**
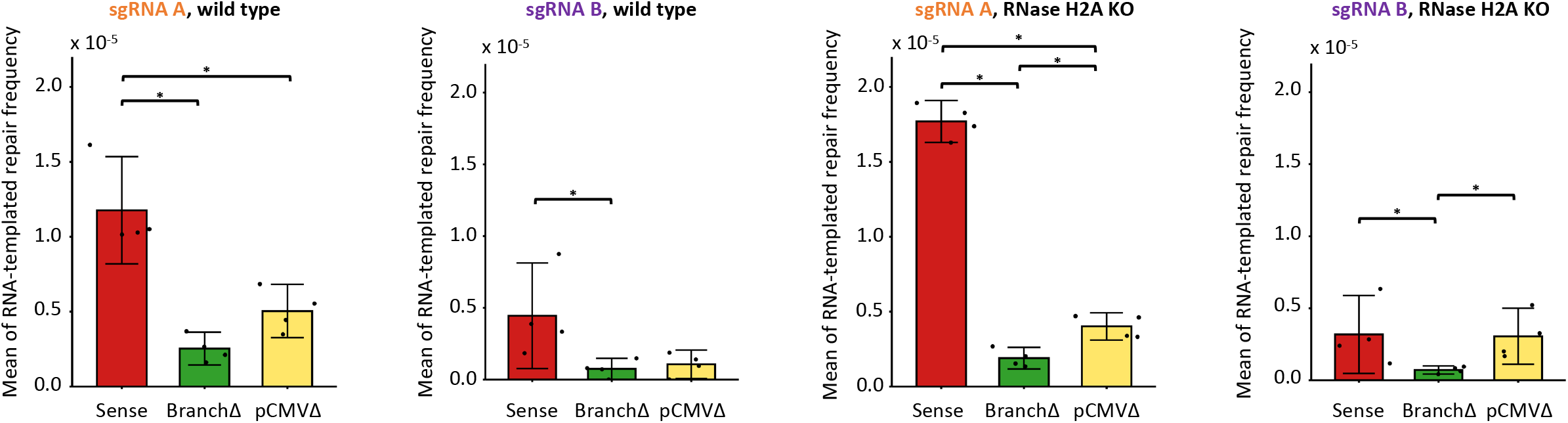
Transcript RNA is a template for DNA DSB repair in human cells. Frequency of R-TDR using the spliced transcript as a template following a DSB by the sgRNA A or B in the Sense (red), BranchΔ (green), and pCMVΔ (yellow) constructs of wild-type (left) and RNase H2A KO (right) cells. Data are mean ± s.d. with 4 biological replicates indicated with dots; N=4. *, *P* = 0.029 (two-tailed Mann-Whitney *U* test).

## Discussion

Here we found that a transcript RNA mediates DNA DSB repair in a sequence-specific manner in human cells. We developed genetic constructs expressing RNA transcripts that differ by the presence or absence of an intron, or the level of transcription due to the presence or absence of the CMV promoter. When the transcript RNA is complementary to the DNA sequences at the DSB ends on both sides of the DSB (non-splicing constructs: BranchΔ and 5′-SplicingΔ), it more often supports the conservation of the original uncut sequence via NHEJ, compared to the RNA that does not have complementarity to both the DSB ends (splicing constructs: Sense, pCMVΔ, and Antisense). On the contrary, the transcript RNA that has complementarity distant from the DSB ends more often promotes deletion of the DNA region near the DSB which it does not share complementarity with, both via MMEJ and via NHEJ in gap repair. We propose two new mechanisms of RNA-mediated DSB repair: RNA-mediated NHEJ (R-NHEJ) and RNA-mediated MMEJ (R-MMEJ), in which the RNA, due to its complementarity to the DNA sequence from which it is transcribed, bridges the DSB ends in a way that facilitates NHEJ or MMEJ, respectively (**Figure 6**).

**Figure 6.**
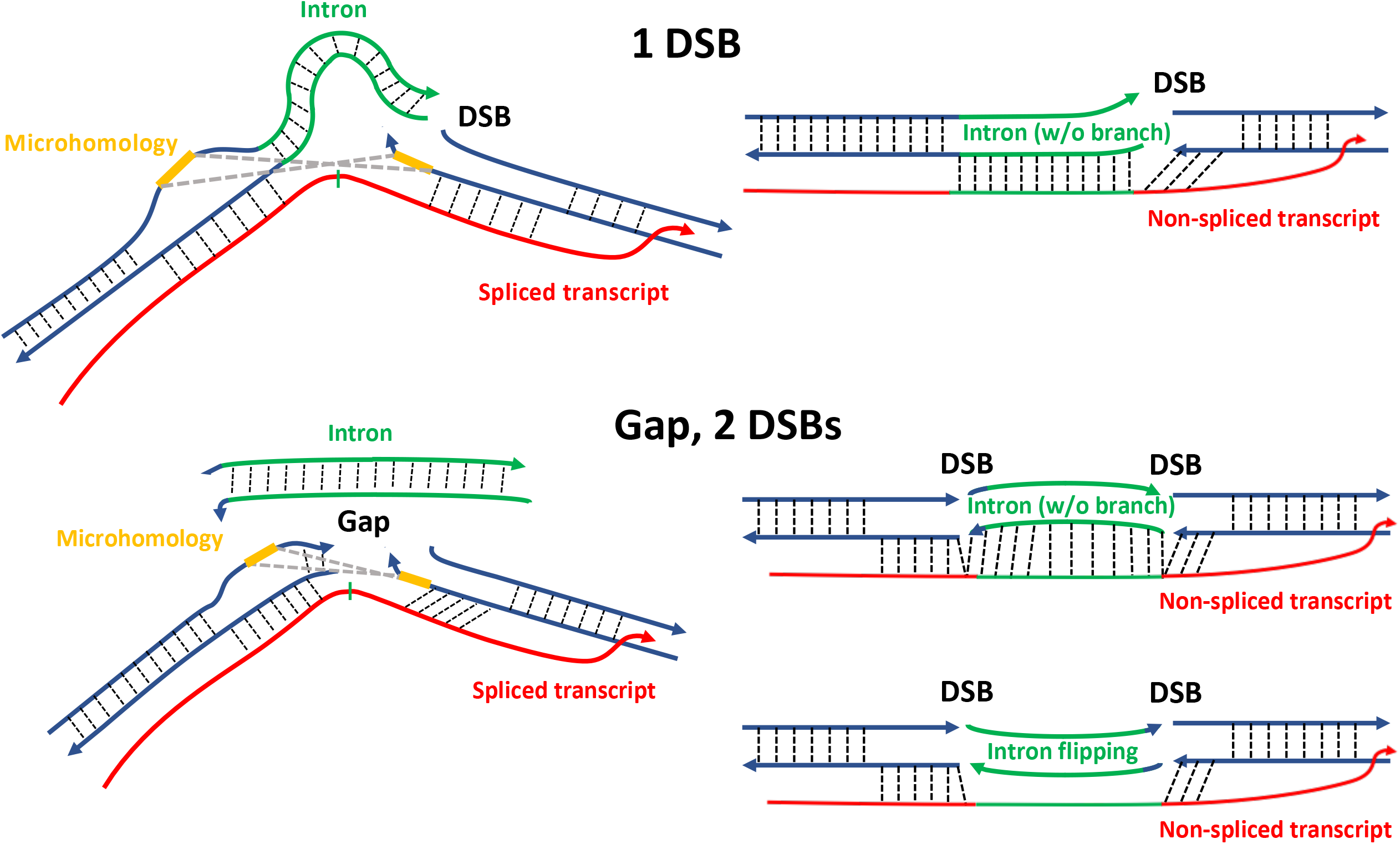
Models of transcript RNA-mediated DSB repair. When a DSB is generated near one or both exon-intron junctions of a gene, the transcript RNA of that gene supports DSB/gap repair with intron pop-out if spliced (schemes on the left), or intron retention if non-spliced (schemes on the right). On the other hand, the non-spliced transcript RNA by interacting with the DSB ends of a gap on both sides of the intron, maintains the gap ends distant from each other allowing intron flipping (scheme on the bottom right). Dotted lines, hydrogen bonds; blue lines, exon sequences; green lines, intron sequences in DNA or RNA; red line, the gene RNA transcript; thin green mark, splice site of the intron in RNA; yellow lines, microhomology; gray, dashed lines, the interaction between the microhomologies.

In addition to R-NHEJ and R-MMEJ, we detected the R-TDR signature in the human cells. The R-TDR signature, which could be due to RNA or to a DNA copy (cDNA) being the template for DSB repair, was found in both wild-type and RNase H2A KO cells. It was recently reported that the knock-out of *RNASEH2A* abolishes LINE1 activity in HEK-293T and HeLa cells^19^. Thus, cDNA of the *DsRed* transcript is unlikely to be the template for R-TDR, particularly in the RNase H2A KO cells. Moreover, LINE1 RT often uses DSB ends, like those generated by CRISPR/Cas9 cleavage, as primers for reverse transcription of RNA complementary to the DSB ends^20^. This is different from the yeast retrotransposon that generates cDNA in retroviral-like particles within the cytoplasm of cells^21^. Therefore, the LINE1 RT could be the polymerase directly using the RNA template in R-TDR. The R-TDR signature we found in the human cells is in the order of 10^−5^. This frequency is much higher than the R-TDR frequency in yeast cells, in which it was not possible to detect R-TDR in RNase H1, H2, nor H1 and H2 wild-type cells, out of 10^8^ viable cells^10,11^. The human recombination protein RAD52 was shown to catalyze RNA-DNA annealing and RNA-inverse strand exchange, which are key steps of R-TDR, more efficiently than yeast Rad52^10,22^. In addition, human DNA Polϑ, which is not present in yeast, was recently shown to exhibit significant RNA-dependent DNA synthesis activity in vitro and was proposed to play a role in RNA-templated DNA repair^23^. Alternatively, as shown in yeast^11^, R-TDR could exploit the RNA-templated synthesis by DNA Polζ in human cells. The results for R-TDR in the pCMVΔ are consistent with previous findings in yeast RNase H1 and H2 knock-out cells that R-TDR is significantly reduced by a low level of transcription of the template RNA^10^.

We also showed that, contrary to R-TDR, a low level of transcription of the RNA still has a major impact on R-NHEJ and/or R-MMEJ in the assays we used. Nonetheless, the transcription level does impact R-NHEJ and R-MMEJ. The frequency of NHEJ repair was similar in both the Sense and pCMVΔ constructs for a DSB by sgRNA A, but significantly lower in the pCMVΔ construct for a DSB by sgRNA B (**Figure 2a** and bottom right panels in **Extended Data Figure 8b, d**), suggesting that the amount of RNA from the pCMVΔ construct still carrying the intron over the DSB site of sgRNA B is likely less than that from the Sense construct. Thus, in this case, the low level of transcription of the spliced RNA in the pCMVΔ construct decreases the efficiency of NHEJ repair of a DSB at the 3′-intron-exon junction compared to not only the non-spliced RNA of the BranchΔ construct, but also the highly-expressed spliced RNA of the Sense construct. Moreover, we established that both coding and non-coding RNA guides DSB repair via NHEJ and MMEJ. Because non-coding RNAs are usually expressed at a low level compared to mRNAs of coding genes^24^, the results suggest that transcript RNA, regardless of its type and level of transcription, plays a significant role in genome stability. It is interesting to note that while introns in mRNAs tend to be removed co-transcriptionally, introns in long non-coding RNAs (lncRNAs) tend to be spliced post-transcriptionally^25–27^. This suggests that non-coding RNA of lncRNAs even if transcribed at low levels, could serve a more general function than anticipated in maintaining the integrity of both exonic and intronic regions in the genome.

The variation-distance graphs applied to the analysis of the sequencing reads derived from R-NHEJ gave a detailed snapshot of the DSB-repair sequence variation types (insertions and deletions), nucleotide compositions, and their frequencies. In the 1-DSB system, we found that graphs arising from different guide RNAs had marked differences, while those arising from the same guide RNA were similar to each other. In graphs corresponding to sgRNA A, inserted nucleotides had a preference to start with T, while in those of sgRNA B, inserted nucleotides had no preference among the four bases (**Figure 2b, Extended Data Figure 8a, c**). Such difference in the in/del type of NHEJ repair at the two-DSB sites supports marked dependence on the sequence context, as previously reported^28^. Interestingly, in all constructs, the generation of a gap in the 2-DSB system produced greater variety of in/dels (i.e., different vertices in the graph) after repair by NHEJ than in the 1-DSB systems, particularly for deletions (compare graphs in **Figure 4c** and **Extended Data Figure 10a, c** with those in **Figure 2b** and **Extended Data Figure 8a, c**). Moreover, although the variation of the in/dels among the different constructs is similar within the 1-DSB and the 2-DSB systems, the relative frequencies of the corresponding in/dels were consistently higher in one construct than in the other as the comparison graphs indicate.

Future work, directed at investigating not just genetic players but also how the position of the DSB relative to exon and intron sequences of genes affects RNA-mediated DSB repair, will help us understand the dynamics and the impact of RNA-mediated DSB repair in human cells. The most efficient mechanism of DSB repair in human cells is NHEJ^1,2^. This study shows that the transcript RNA of a broken gene helps NHEJ when the RNA retains complementarity with the DSB ends. Therefore, the results suggest that DSB repair mediated by a spliced mRNA may more efficiently stabilize gene regions that experience a DSB in exonic sequences rather than near exon-intron junctions or in intronic sequences, for which the spliced mRNA does not have complementarity to one or both broken DNA ends of a DSB, respectively.

RNA could also play a role in destabilizing the genome. The experiments, which involved a cut on both sides of the intron in the *DsRed* gene of the sense and antisense constructs, revealed that the non-spliced constructs have a higher frequency of intron flipping than the spliced constructs. These results highlight the capacity of RNA to hold DNA ends not just close to each other, but also distant from each other (see bottom right scheme in **Figure 6**), depending on the sequence complementarity between the RNA and the DNA. Thus, RNA can guide genome alterations. In summary, we identified a novel function of RNA in modulating the repair of DSBs and DNA gaps. The findings provide new avenues for understanding the mechanisms of genome integrity, genome modification, and evolution.

## Acknowledgements

We are grateful to D. Garfinkel for plasmid pSM50, and to M. Jasin for the enhanced empty gRNA expression vector. We thank N. Djeddar and A. Bryksin for their technical assistance with sequencing. We thank K. Lobachev, J. Dahlam, and Y. Lu for critical reading of the manuscript, and all the members of the Storici and Jonoska groups for assistance and feedback on this research. We acknowledge funding from the National Institute of General Medical Sciences (NIGMS) of the NIH (grant GM115927 to F.S.), the National Institute of Environmental Health Sciences (NIEHS) of the NIH (grant ES028271 to Z.F.P.), the National Science Foundation fund (grant MCB-1615335 to F.S.), the Howard Hughes Medical Institute Faculty Scholar (grant 55108574 to F.S.), and grants from the Southeast Center for Mathematics and Biology (NSF [DMS-1764406] and the Simons Foundation [594594] to F.S. and N.J.) for supporting this work.

## Author Contributions

F.S. together with Y.J., M.M.F. and N.J. conceived the project and designed experiments. F.S. wrote the manuscript with help from Y.J., M.M.F., T.C., P.X., N.J., and help from all authors. Y.J. performed all the experiments with help from S.B. and S.M. Y.J., M.M.F., T.C., and P.X. worked on the sequencing data analyses. M.M.F. and T.C. worked on the variation-distance graphs. V.S.P. and Z.F.P. constructed the RNase H2A KO cell lines. All authors commented on and approved the manuscript.

## References

1 Mao, Z., Bozzella, M., Seluanov, A. & Gorbunova, V. Comparison of nonhomologous end joining and homologous recombination in human cells. DNA Repair (Amst) 7, 1765–1771 (2008).

2 Scully, R., Panday, A., Elango, R. & Willis, N. A. DNA double-strand break repair-pathway choice in somatic mammalian cells. Nat Rev Mol Cell Biol 20, 698–714 (2019).

3 Ramsden, D. A., Carvajal-Garcia, J. & Gupta, G. P. Mechanism, cellular functions and cancer roles of polymerase-theta-mediated DNA end joining. Nat Rev Mol Cell Biol 23, 125–140 (2022).

4 McVey, M. & Lee, S. E. MMEJ repair of double-strand breaks (director’s cut): deleted sequences and alternative endings. Trends Genet 24, 529–538 (2008).

5 Pertea, M. The human transcriptome: an unfinished story. Genes (Basel) 3, 344–360 (2012).

6 Lee, H., Zhang, Z. & Krause, H. M. Long Noncoding RNAs and Repetitive Elements: Junk or Intimate Evolutionary Partners? Trends Genet 35, 892–902 (2019).

7 Consortium, E. P. An integrated encyclopedia of DNA elements in the human genome. Nature 489, 57–74 (2012).

8 Djebali, S. et al. Landscape of transcription in human cells. Nature 489, 101–108 (2012).

9 Kazimierczyk, M., Kasprowicz, M. K., Kasprzyk, M. E. & Wrzesinski, J. Human Long Noncoding RNA Interactome: Detection, Characterization and Function. Int J Mol Sci 21 (2020).

10 Keskin, H. et al. Transcript-RNA-templated DNA recombination and repair. Nature 515, 436–439 (2014).

11 Meers, C. et al. Genetic Characterization of Three Distinct Mechanisms Supporting RNA-Driven DNA Repair and Modification Reveals Major Role of DNA Polymerase zeta. Mol Cell 79, 1037–1050 e1035 (2020).

12 Petermann, E., Lan, L. & Zou, L. Sources, resolution and physiological relevance of R-loops and RNA-DNA hybrids. Nat Rev Mol Cell Biol 23, 521–540 (2022).

13 Ali, A., Xiao, W., Babar, M. E. & Bi, Y. Double-Stranded Break Repair in Mammalian Cells and Precise Genome Editing. Genes (Basel) 13 (2022).

14 Meers, C., Keskin, H. & Storici, F. DNA repair by RNA: Templated, or not templated, that is the question. DNA Repair (Amst) 44, 17–21 (2016).

15 Chakraborty, A. et al. Classical non-homologous end-joining pathway utilizes nascent RNA for error-free double-strand break repair of transcribed genes. Nat Commun 7, 13049 (2016).

16 Schep, R. et al. Impact of chromatin context on Cas9-induced DNA double-strand break repair pathway balance. Mol Cell 81, 2216–2230 e2210 (2021).

17 Bader, A. S. & Bushell, M. DNA:RNA hybrids form at DNA double-strand breaks in transcriptionally active loci. Cell Death Dis 11, 280 (2020).

18 Gasior, S. L., Wakeman, T. P., Xu, B. & Deininger, P. L. The human LINE-1 retrotransposon creates DNA double-strand breaks. J Mol Biol 357, 1383–1393 (2006).

19 Bartsch, K. et al. Absence of RNase H2 triggers generation of immunogenic micronuclei removed by autophagy. Hum Mol Genet 26, 3960–3972 (2017).

20 Tao, J., Wang, Q., Mendez-Dorantes, C., Burns, K. H. & Chiarle, R. Frequency and mechanisms of LINE-1 retrotransposon insertions at CRISPR/Cas9 sites. Nat Commun 13, 3685 (2022).

21 Curcio, M. J., Lutz, S. & Lesage, P. The Ty1 LTR-Retrotransposon of Budding Yeast, Saccharomyces cerevisiae. Microbiol Spectr 3, MDNA3-0053-2014 (2015).

22 Mazina, O. M., Keskin, H., Hanamshet, K., Storici, F. & Mazin, A. V. Rad52 Inverse Strand Exchange Drives RNA-Templated DNA Double-Strand Break Repair. Mol Cell 67, 19–29 e13 (2017).

23 Chandramouly, G. et al. Poltheta reverse transcribes RNA and promotes RNA-templated DNA repair. Sci Adv 7 (2021).

24 Uszczynska-Ratajczak, B., Lagarde, J., Frankish, A., Guigo, R. & Johnson, R. Towards a complete map of the human long non-coding RNA transcriptome. Nat Rev Genet 19, 535–548 (2018).

25 Derrien, T. et al. The GENCODE v7 catalog of human long noncoding RNAs: analysis of their gene structure, evolution, and expression. Genome Res 22, 1775–1789 (2012).

26 Tilgner, H. et al. Deep sequencing of subcellular RNA fractions shows splicing to be predominantly co-transcriptional in the human genome but inefficient for lncRNAs. Genome Res 22, 1616–1625 (2012).

27 Palazzo, A. F. & Lee, E. S. Non-coding RNA: what is functional and what is junk? Front Genet 6, 2 (2015).

28 Lemos, B. R. et al. CRISPR/Cas9 cleavages in budding yeast reveal templated insertions and strand-specific insertion/deletion profiles. Proc Natl Acad Sci U S A 115, E2040–E2047 (2018).

