## Supplementary materials for "RNA-mediated double-strand break repair in human cells"

#### Extended Data Fig. 1

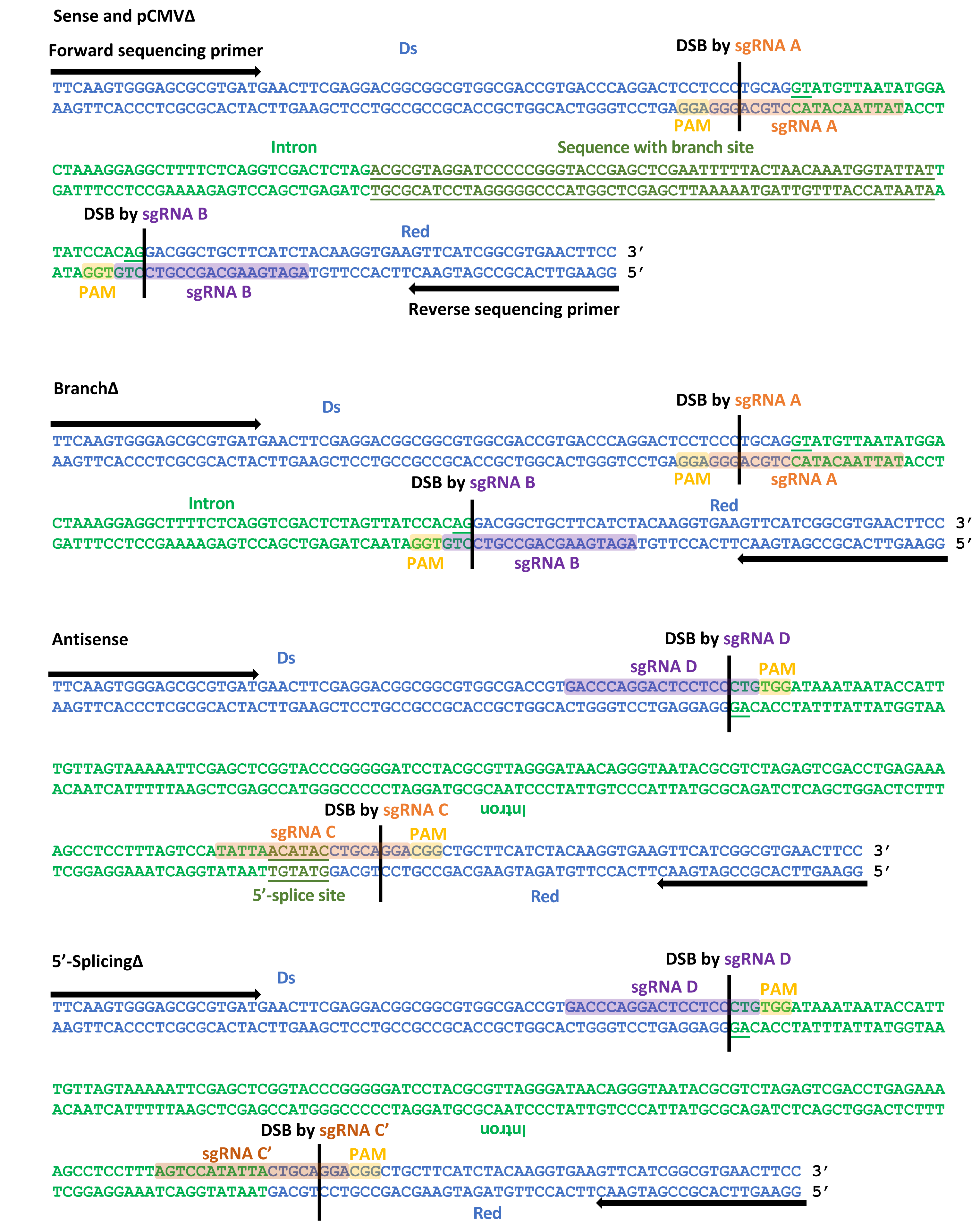

**Extended Data Figure 1 | DNA sequence of the *DsRed* loci of the sense and antisense constructs.**
Blue sequence, exons of the *DsRed* gene; green sequence, intron; underlined green sequence, canonical GT- and -AG splice sites of the intron; underlined dark green sequence, intron sequence containing the branch site (Sense and pCMVΔ) or the 5′-splice site (Antisense). Black arrows, sequencing primers; yellow highlighted sequence, PAM site; orange highlighted sequence, sequence of sgRNA A, C or C'; purple highlighted sequence, sequence of sgRNA B or D. Vertical black bar, site of DSB by sgRNA.

#### Extended Data Fig. 2

**a**

| Plasmid | spliced transcript frequency | non-spliced transcript frequency | alt-spliced transcript frequency | non-canonical alt-spliced frequency | unaligned frequency |
|---|---|---|---|---|---|
| Sense | **0.96** | 0.0048 | 0.00043 | 0.0077 | 0.026 |
| BranchΔ | 0.0034 | **0.95** | 0.000015 | 0.010 | 0.035 |
| pCMVΔ | **0.98** | 0.0043 | 0.000076 | 0.0032 | 0.014 |
| Antisense | **0.16** | 0.70 | 0.092 | 0.0068 | 0.043 |
| 5'-SplicingΔ | 0.0042 | **0.92** | 0.033 | 0.0075 | 0.034 |

**b**

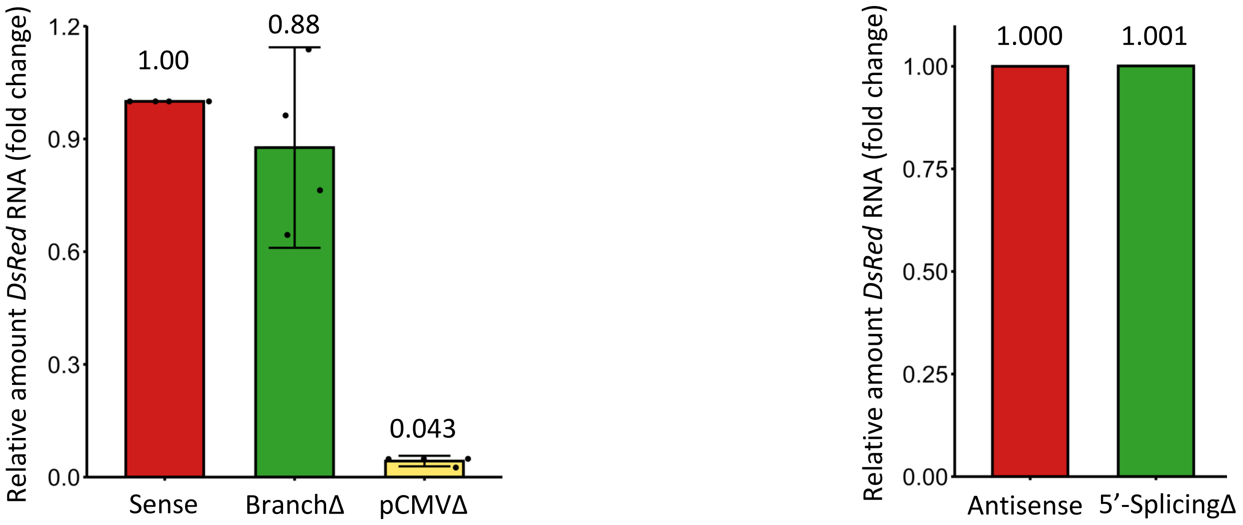

**Extended Data Figure 2 | The RNA transcribed from the BranchΔ and the 5′-SplicingΔ constructs has no intron splicing, and the one transcribed from the pCMVΔ is in low amount.**

**a,** Table of RNA sequencing results of *DsRed* transcripts from the Sense, BranchΔ, and pCMVΔ, as well as the Antisense and the 5′-SplicingΔ constructs. RNA sequencing reads for each transcript of the different constructs were categorized after alignment to the corresponding DNA sequence (details in Methods). Spliced transcript, a transcript in which the intron was spliced out from the canonical GT---AG-splice sites (see **Extended Data Figure 1**); alt-spliced transcript, a transcript in which splicing happened at alternative GT---AG sites; non-canonical alternative splicing, a transcript sequence that had splicing at sites different from the GT---AG sites. Splicing frequencies were calculated by dividing the number of the spliced-transcript reads by the total number of reads within each RNA sequencing library. **b,** Results of RT-qPCR of the *DsRed* transcripts generated from the sense and antisense constructs. Shown on the left is the relative amount of the *DsRed* transcripts from the BranchΔ (green bar) and the pCMVΔ (yellow bar) constructs compared to the amount of the Sense (red bar) transcripts. Shown on the right is the relative amount of the *DsRed* transcripts from the 5′-SplicingΔ construct (green bar) compared to the amount of the Antisense construct (red bar). Plotted data are the mean fold change ± s.d. with 4 biological replicates indicated with dots; N=4. Individual values are plotted. The mean value is shown above each bar.

**Extended Data Fig. 3**

**a**

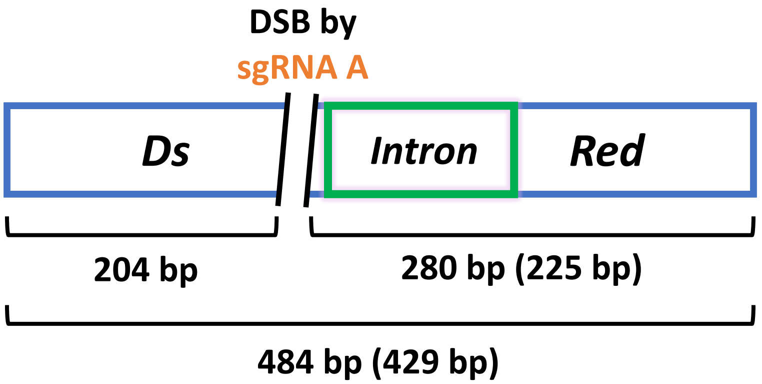


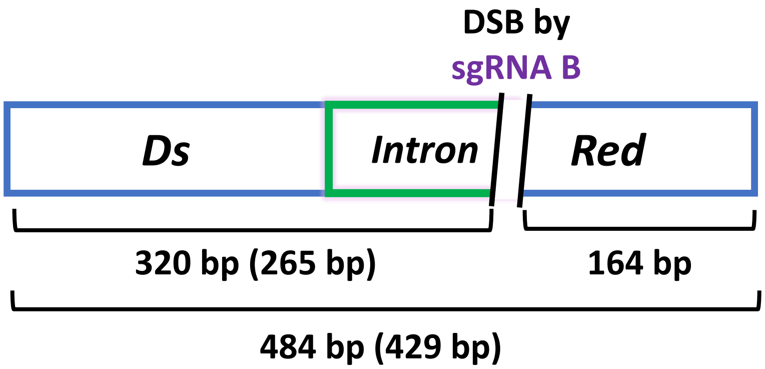


$$DSB\ efficiency = \frac{\frac{204bp + 280bp}{2}}{\frac{204bp + 280bp}{2} + 484bp}$$

(Calculate by molarity)

$$DSB\ efficiency = \frac{\frac{320bp + 164bp}{2}}{\frac{320bp + 164bp}{2} + 484bp}$$

(Calculate by molarity)

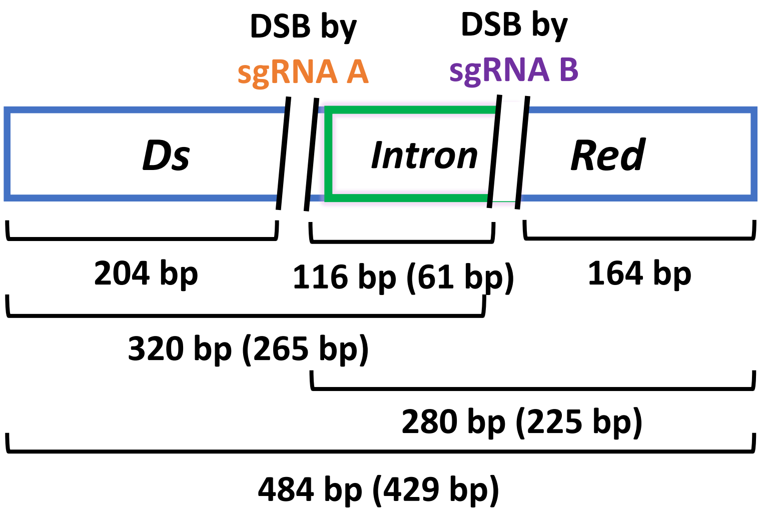


$$2\text{-}DSB\ efficiency = \frac{116bp}{116bp + 280bp + 320bp + 484bp}$$

(Calculate by molarity)

| **sgRNA A** | 1st | 2nd | 3rd | 4th | Average |
|---|---|---|---|---|---|
| Sense | **0.936** | **0.939** | **0.952** | **0.938** | **0.941** |
| BranchΔ | **0.939** | **0.943** | **0.945** | **0.941** | **0.942** |

| **sgRNA B** | 1st | 2nd | 3rd | 4th | Average |
|---|---|---|---|---|---|
| Sense | **0.943** | **0.940** | **0.937** | **0.944** | **0.941** |
| BranchΔ | **0.952** | **0.955** | **0.948** | **0.943** | **0.949** |

| **2-DSB** | 1st | 2nd | 3rd | 4th | 5th | 6th | 7th | 8th | 9th | 10th | 11th | Avg. |
|---|---|---|---|---|---|---|---|---|---|---|---|---|
| Sense | **0.890** | **0.891** | **0.883** | **0.879** | **0.891** | **0.882** | **0.884** | **0.885** | **0.876** | **0.882** | **0.889** | **0.885** |
| BranchΔ | **0.884** | **0.878** | **0.879** | **0.876** | **0.870** | **0.851** | **0.880** | **0.879** | **0.873** | **0.873** | | **0.874** |

**Extended Data Fig. 3**

**b**

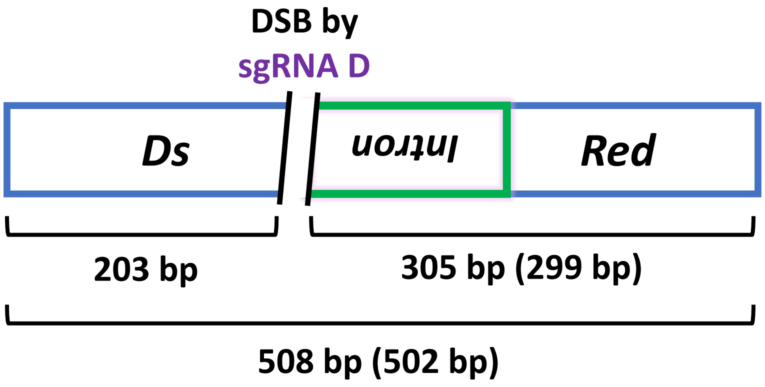


$$\text{DSB efficiency} = \frac{\frac{203bp+305bp}{2}}{\frac{203bp+305bp}{2}+508bp}$$

(Calculate by molarity)

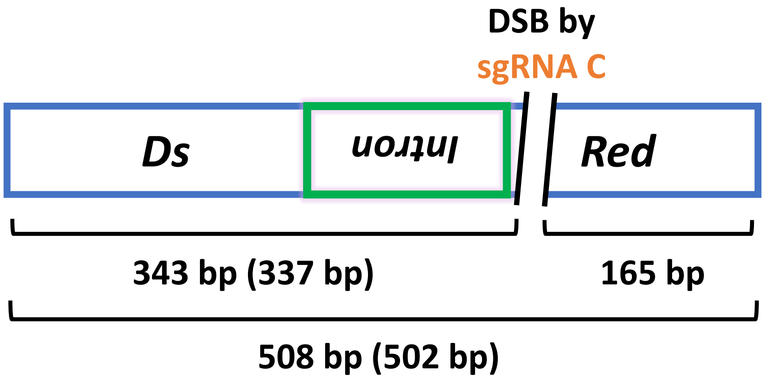


$$\text{DSB efficiency} = \frac{\frac{343bp+165bp}{2}}{\frac{343bp+165bp}{2}+508bp}$$

(Calculate by molarity)

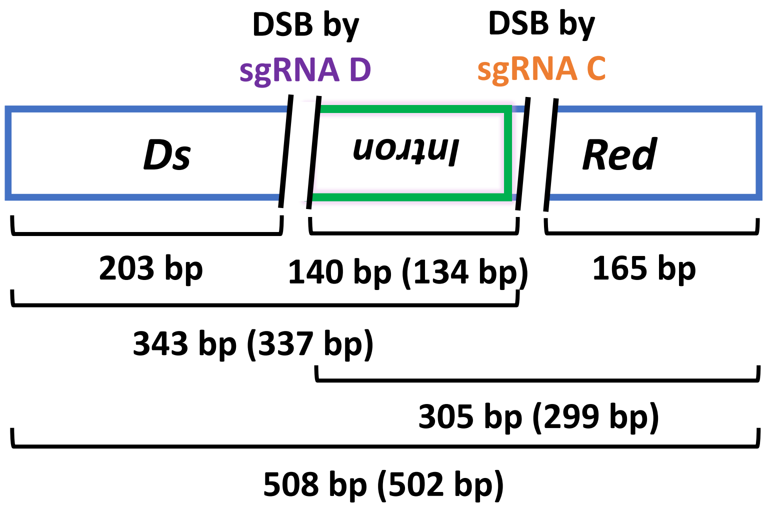


$$\text{2-DSB efficiency} = \frac{140bp}{140bp+305bp+343bp+508bp}$$

(Calculate by molarity)

| **sgRNA C/C'** | 1st | 2nd | 3rd | 4th | Average |
|---|---|---|---|---|---|
| Antisense (sgRNA C) | **0.949** | **0.947** | **0.955** | **0.952** | **0.951** |
| 5'-SplicingΔ (sgRNA C') | **0.961** | **0.952** | **0.957** | **0.953** | **0.956** |

| **sgRNA D** | 1st | 2nd | 3rd | 4th | Average |
|---|---|---|---|---|---|
| Antisense | **0.836** | **0.824** | **0.846** | **0.849** | **0.839** |
| 5'-SplicingΔ | **0.864** | **0.861** | **0.855** | **0.861** | **0.860** |

| **2-DSB** | 1st | 2nd | 3rd | 4th | 5th | 6th | Avg. |
|---|---|---|---|---|---|---|---|
| Antisense | **0.835** | **0.822** | **0.791** | **0.844** | **0.824** | **0.843** | **0.826** |
| 5'-SplicingΔ | **0.835** | **0.853** | **0.846** | **0.860** | **0.854** | | **0.849** |

**Extended Data Figure 3 | Constructs with splicing have similar cleavage efficiency by Cas9 to those without splicing.** Scheme of PCR fragments for the (**a**) Sense or pCMVΔ construct, and the (**b**) Antisense construct used for the in-vitro cleavage assay. Cas9 cleavage was done using sgRNA A, B or A and B for the sense (Sense/pCMVΔ and BranchΔ) constructs, and by using sgRNA C/C', D or C/C' and D for the antisense (Antisense and 5′-SplicingΔ) constructs. The formulas used to calculate the DSB efficiencies from the molarity of each DNA fragment detected by the Bioanalyzer following cleavage by Cas9 with sgRNA A, B, or A and B for the sense constructs (**a**), or sgRNA C/C', D, or C/C' and D for the antisense constructs (**b**) are shown. The bp sizes in the formulas are those for the Sense/pCMVΔ and BranchΔ (in parenthesis when different) in (**a**), and those for the Antisense and 5′-SplicingΔ (in parenthesis when different) in (**b**). Results of the in-vitro cleavage assay for the sense constructs (**a**), and the antisense constructs (**b**) are shown in the tables underneath the construct schemes.

### Extended Data Fig. 4

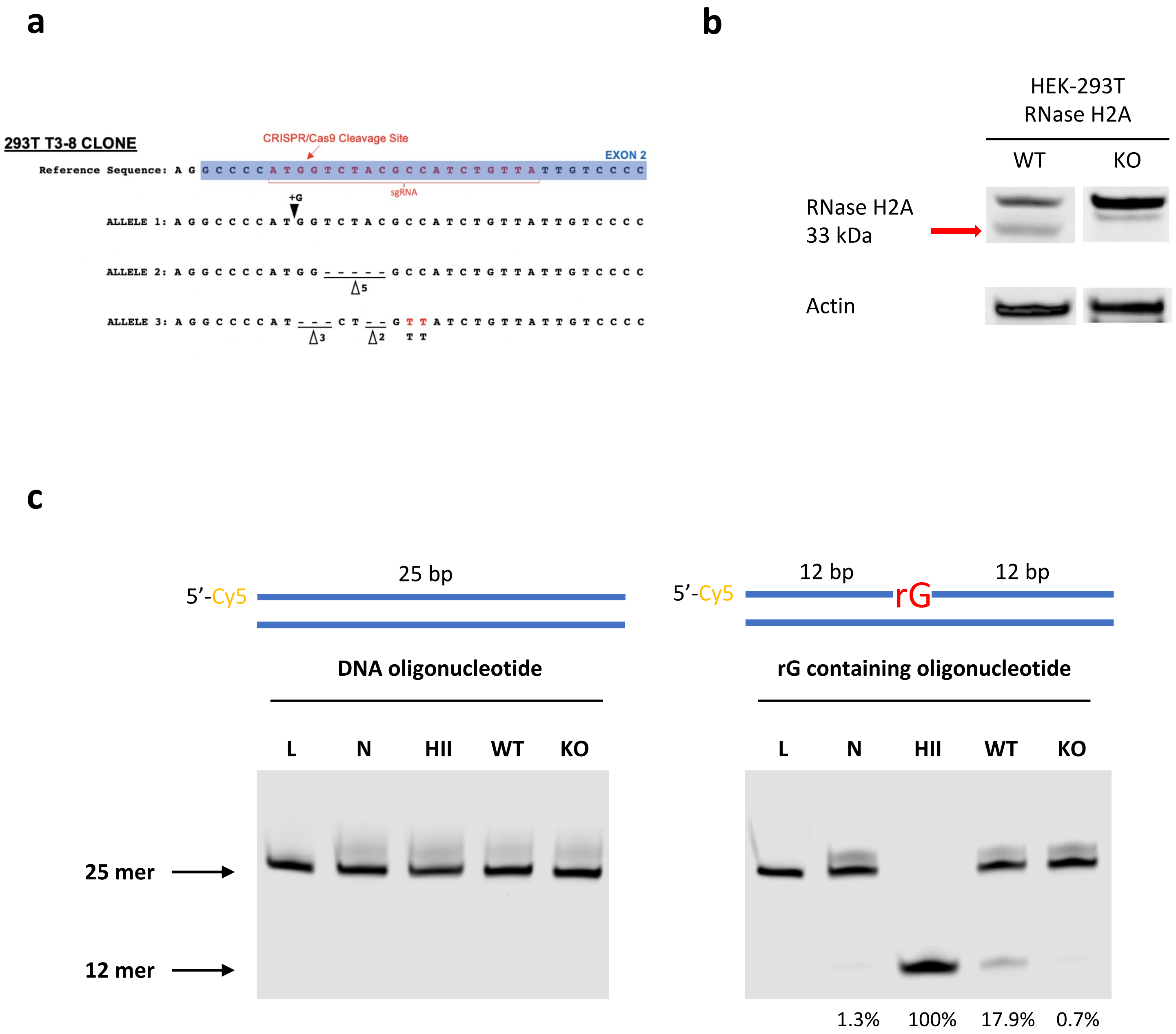

**Extended Data Figure 4 | Characterization of RNase H2A KO in HEK-293T cells.**

**a,** Mutated sequences of the *RNASEH2A* alleles from the RNase H2A KO HEK-293T cells. **b**, Result of Western blot for RNase H2A in HEK-293T RNase H2A wild-type and KO cells. **c,** Double-stranded Cy5- labeled 25-mers oligos were used in the *in vitro* cleavage assay to evaluate RNase H2 activity in protein extracts from HEK-293T RNase H2A wild-type and KO cells, schemes shown on the top of each gel image. L, ladder for 25-mers oligonucleotide; N, negative control with the double-stranded oligonucleotide treated by water; HII, *Escherichia coli* RNase HII was used as a positive control cleaving 5′ of the rGMP embedded in the double-stranded DNA oligonucleotides; WT, protein extract from HEK-293T RNase H2A wild-type cells; KO, protein extract from HEK-293T RNase H2A KO cells. The cleavage % is shown underneath the image.

#### Extended Data Fig. 5

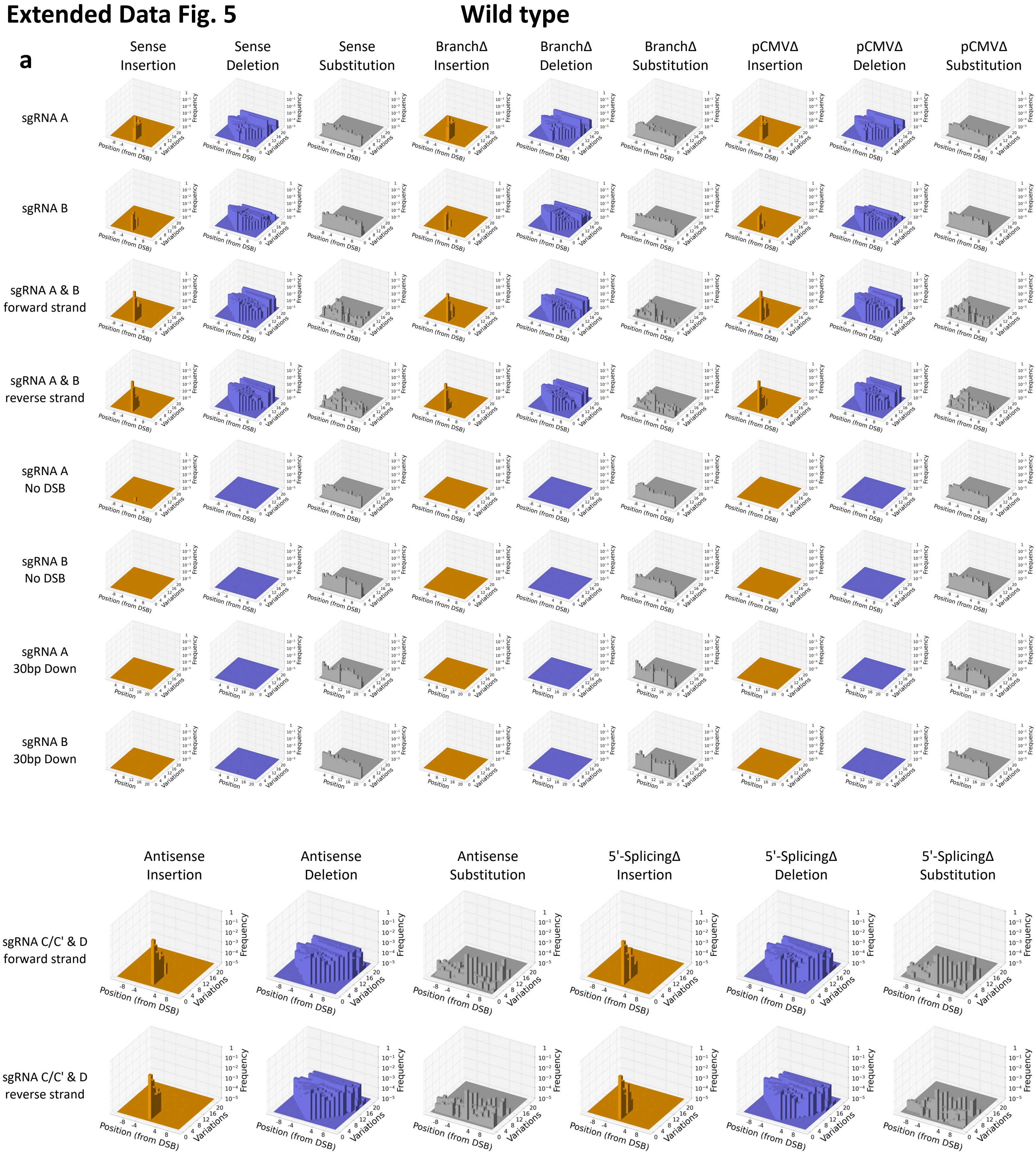

### Extended Data Fig. 5

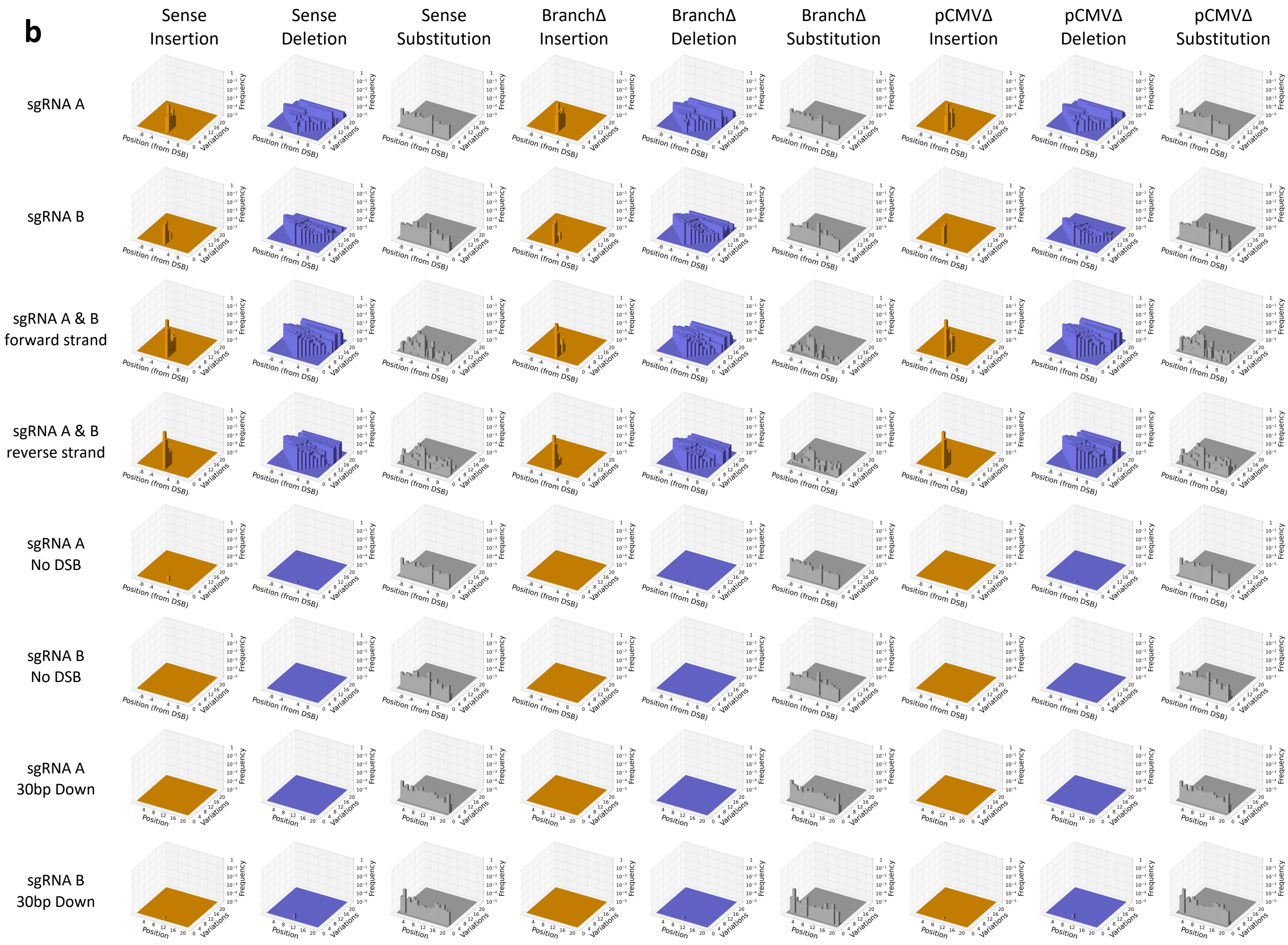

**Extended Data Figure 5 | In/dels are sequence variations specific to DSB repair by NHEJ.**
Variation-position histograms showing the distribution of sequence variations in the 20-bp DSB-sequence windows categorized by the position of the variations and the number of variations. The histograms are arranged in a grid: rows specify the sgRNA(s) used to induce the DSB(s), including the controls (no-DSB and sequence windows 30 bp downstream from the DSB); columns specify the construct and the type of variation (insertion in orange, deletion in blue, or substitution in gray). The *x*-axis indicates the position of the variations relative to the DSB site on the reference sequence. Figures with reverse DSB-sequence windows are reversed such that *x*-axis variation coordinates correspond to forward strand variation coordinates. The *y*-axis indicates the total number of variations in the DSB-sequence windows. The *z*-axis indicates the total sum of frequencies (log-scale) of DSB-sequence windows with *y* variations including position *x*. If a DSB-sequence window has more than one variation (say *k*) at the same position (which can only happen if the DSB-sequence window contains insertions), this window contributes *k*-fold to the corresponding *z*-value. **a,** Variation-position histograms for experiments conducted in the wild-type cells, and **b,** in the RNase H2A KO cells.

### Extended Data Fig. 6

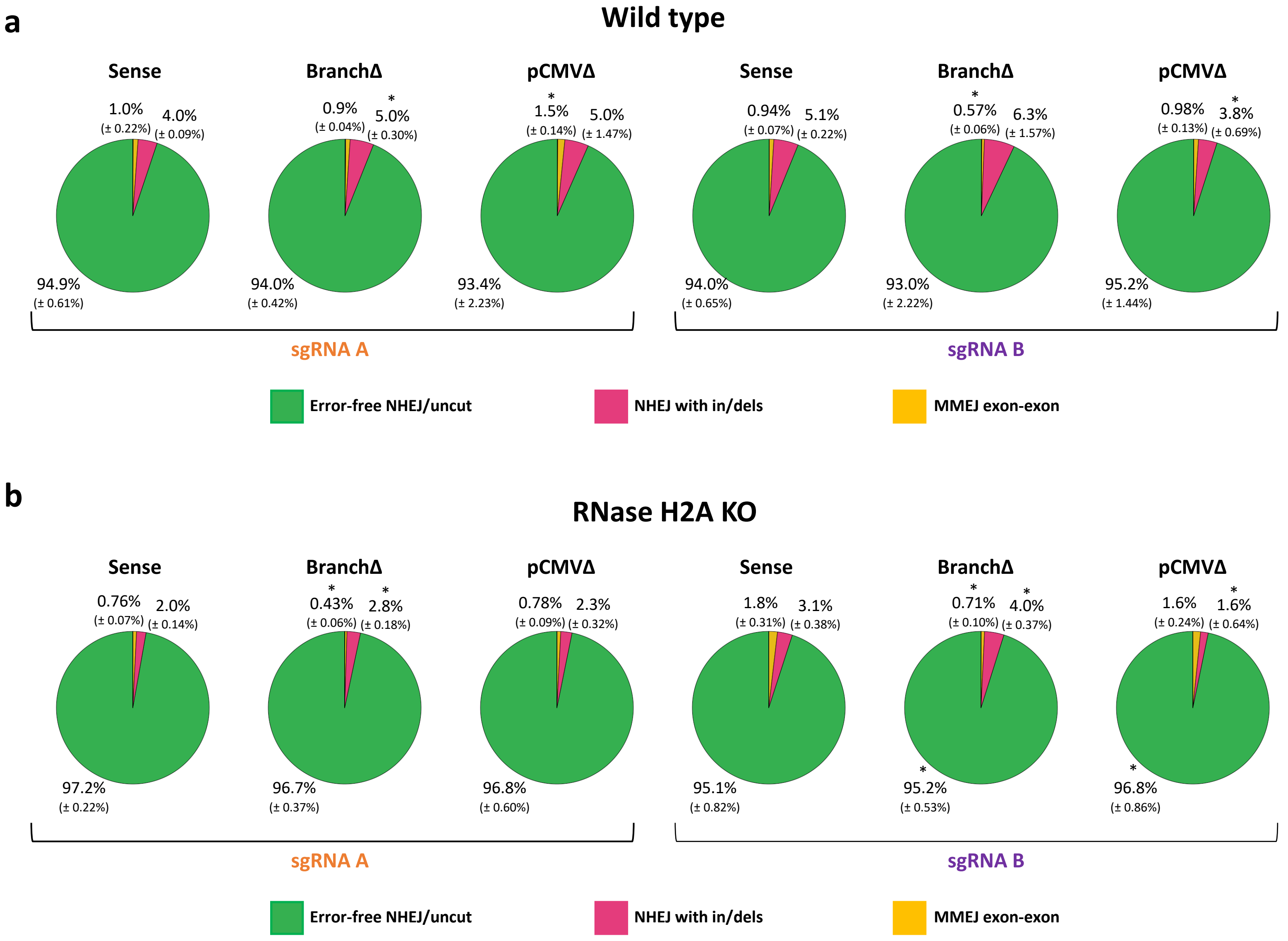

**Extended Data Figure 6 | Transcript RNA affects the repair of a DSB in a sequence-dependent manner via NHEJ and MMEJ**

Pie charts showing frequencies of sequencing reads displaying no change from the uncut sequence (green) or showing signatures of NHEJ (red) or exon-exon MMEJ (yellow) following a DSB by the sgRNA A (left) or B (right) in the Sense, BranchΔ, and pCMVΔ constructs of wild-type (**a**) and RNase H2A KO (**b**) cells. Percentages of the exon-intron MMEJ were 0.07% (wild type, sgRNA A, Sense, ± 0.003%), 0.12% (wild type, sgRNA A, BranchΔ, ± 0.006), 0.07% (wild type, sgRNA A, pCMVΔ, ± 0.013%), 0.03% (wild type, sgRNA B, Sense, ± 0.003%), 0.07% (wild type, sgRNA B, BranchΔ, ± 0.020%), 0.02% (wild type, sgRNA B, pCMVΔ, ± 0.004%), 0.03% (RNase H2A KO, sgRNA A, Sense, ± 0.002%), 0.06% (RNase H2A KO, sgRNA A, BranchΔ, ± 0.008%), 0.03% (RNase H2A KO, sgRNA A, pCMVΔ, ± 0.004%), 0.02% (RNase H2A KO, sgRNA B, Sense, ± 0.002%), 0.03% (RNase H2A KO, sgRNA B, BranchΔ, ± 0.002%), 0.0009% (RNase H2A KO, sgRNA B, pCMVΔ, ± 0.004%). Percentages represent the average of 4 repeats with standard deviations in parenthesis; N=4. *, $P$ = 0.029 comparing frequencies of the BranchΔ or the pCMVΔ with those of the Sense construct via the two-tailed Mann-Whitney $U$ test.

**Extended Data Fig. 7** Variation-distance graph key

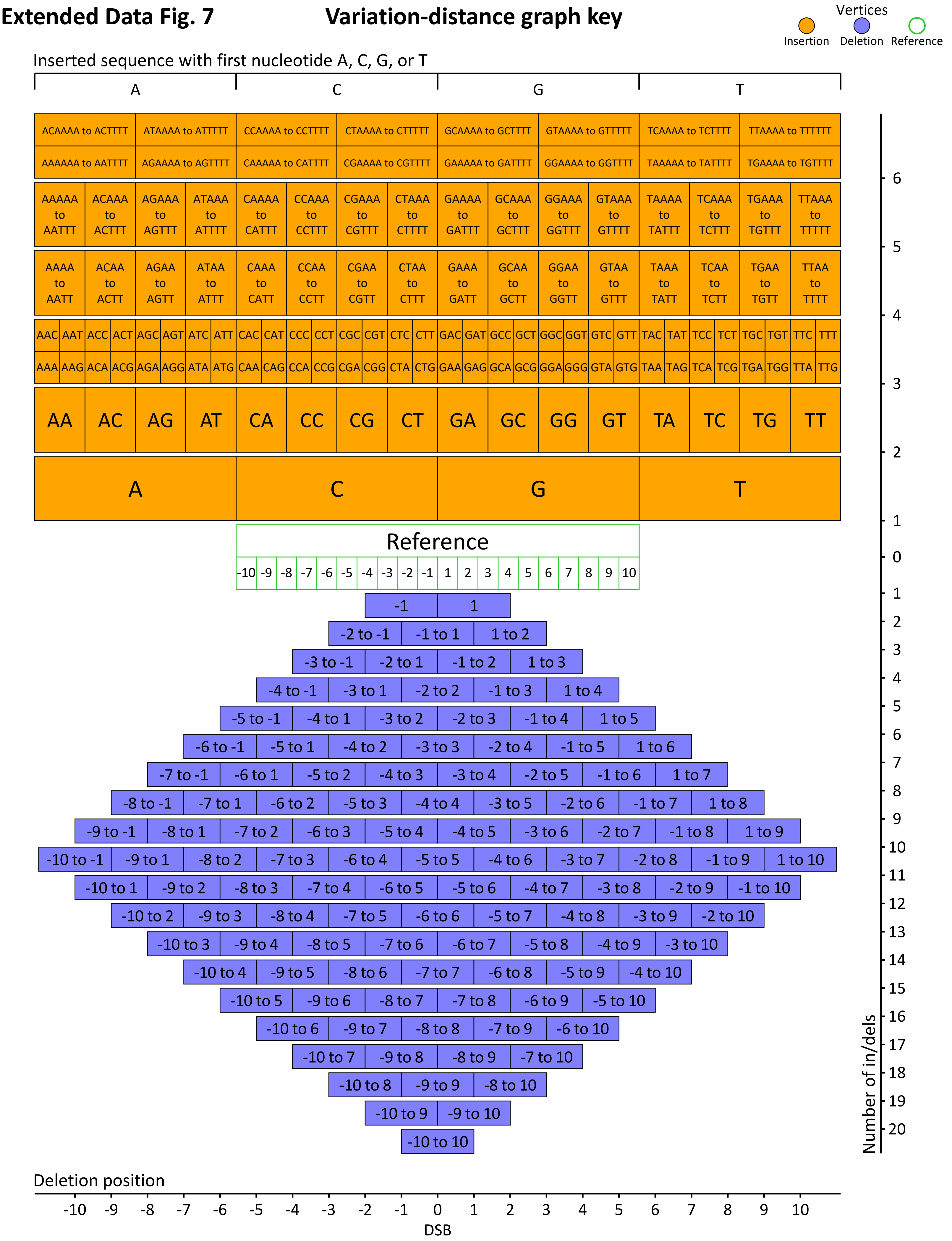

**Extended Data Figure 7 | Variation-distance graph key.**

Variation-distance graph key showing the placement of the vertices representing the analyzed DSB-sequence windows. Vertex placement depends on both the type of variation (insertion or deletion) and the number of variations of the corresponding DSB-sequence window compared to the reference sequence. Insertion vertices are placed above the reference vertex (center), while deletion vertices are placed below it. For insertions, the alphabetical order of the inserted sequence, from A on the left to T on the right, are indicated by the *x*-coordinate. Each insertion box is labeled with the inserted nucleotides of the DSB-sequence window placed in that location. Insertions of size 3 or more have vertices on multiple lines, staggered vertically to reduce overlap. The *x*-coordinate of deletions indicates the position of the first deleted nucleotide in the given DSB-sequence window, from the most upstream (left-most) to the most downstream (right-most). Each deletion box is labeled with the range of the deleted nucleotide sequence, relative to the DSB site, of the DSB-sequence window placed in that location. The *y*-coordinate indicates the number of variations in the DSB-sequence windows, with higher variations placed further from the reference.

**Extended Data Fig. 8**

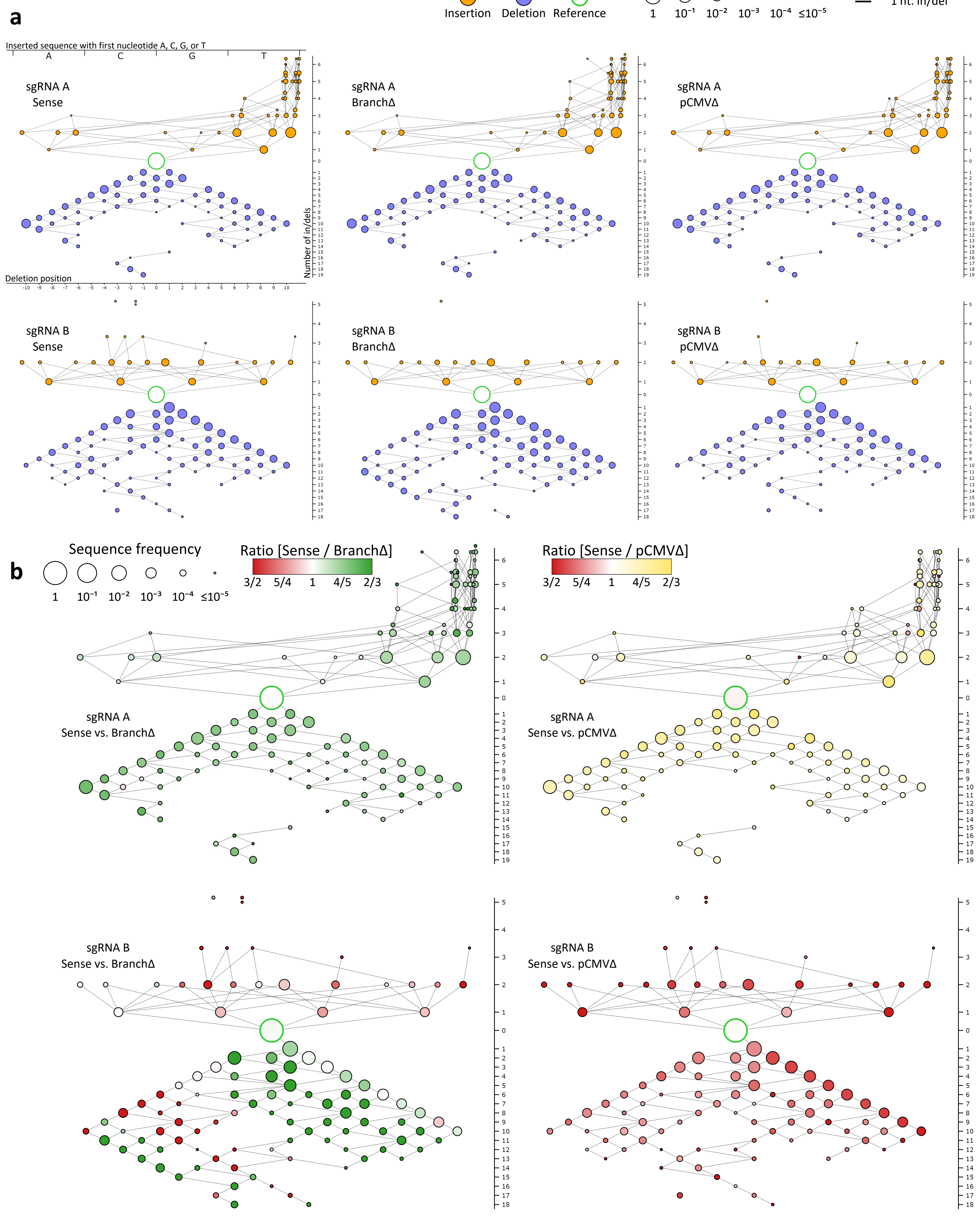

Extended Data Fig. 8

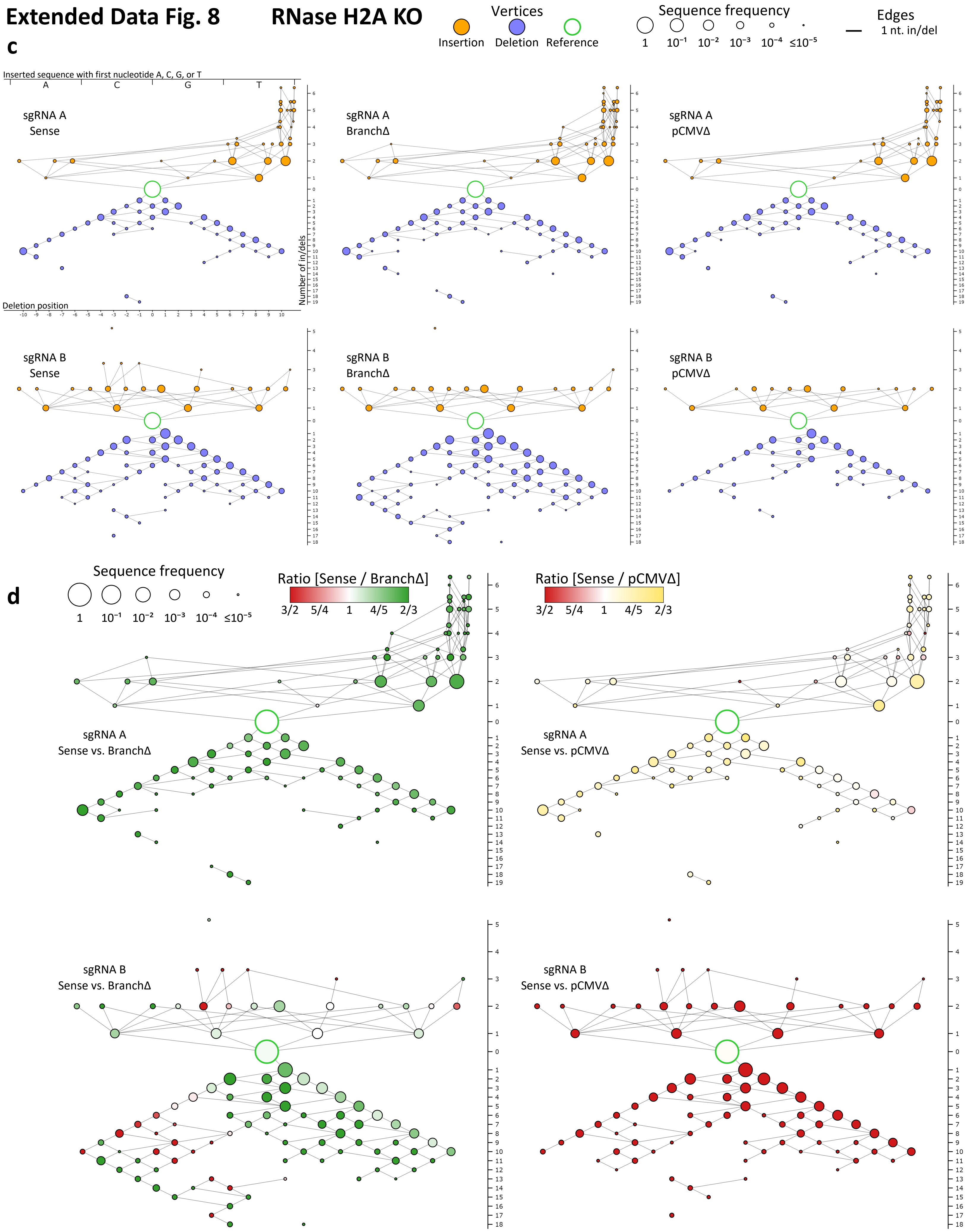

**Extended Data Figure 8 | Transcript RNA enhances DSB repair by NHEJ in a sequence dependent manner – all data for the sense constructs.**

**a,** Individual variation-distance graphs illustrating sequence variations within DSB-sequence windows observed after DSB induction by sgRNA A (top) or sgRNA B (bottom) in the Sense, BranchΔ, and pCMVΔ constructs of wild-type cells. An edge between two vertices indicates that the two corresponding DSB-sequence windows differ by a single nucleotide insertion or deletion (in/del). Insertion vertices (orange circles) are placed above the reference vertex (white circle with a green border), while deletion vertices (blue circles) are placed below it. The vertex size shows the log of the mean frequency of the corresponding DSB-sequence window in the four repeats of the considered experiment. For insertions, the alphabetical order of the inserted sequences, from A on the left to T on the right, are indicated by the *x*-coordinate. Insertions of size 3 or more have vertices on multiple lines, staggered vertically to reduce overlap. The *x*-coordinate of deletions indicates the position of the first deleted nucleotide, from the most upstream (left-most) to the most downstream (right-most). The *y*-coordinate indicates the number of variations in the DSB-sequence windows, with higher variations placed further from the reference. See **Extended Data Fig. 7** for the variation-distance graph key. **b,** Comparison variation-distance graphs of the DSB-sequence windows obtained after DSB induction by sgRNA A (top) or sgRNA B (bottom) for the Sense vs. the BranchΔ constructs (left) or for the Sense vs. the pCMVΔ constructs (right) of wild-type cells. The vertices represent the same DSB-sequence windows as for the individual graphs while the vertex colors specify the relative frequency in the Sense (red) vs. the BranchΔ (green) construct or in the Sense (red) vs. the pCMVΔ (yellow) construct; the vertex sizes show the log of the maximum of the two mean frequencies of the corresponding DSB-sequence windows in the two analyzed constructs. **c**, Individual variation-distance graphs illustrating sequence variations within DSB-sequence windows observed after DSB induction by sgRNA A (top) or sgRNA B (bottom) in the Sense, BranchΔ, and pCMVΔ constructs of RNase H2A KO cells. **d**), Comparison variation-distance graphs of the DSB-sequence windows obtained after DSB induction by sgRNA A (top) or sgRNA B (bottom) for the Sense vs, the BranchΔ constructs (left) or for the Sense vs. the pCMVΔ constructs (right) of RNase H2A KO cells.

Extended Data Fig. 9

a

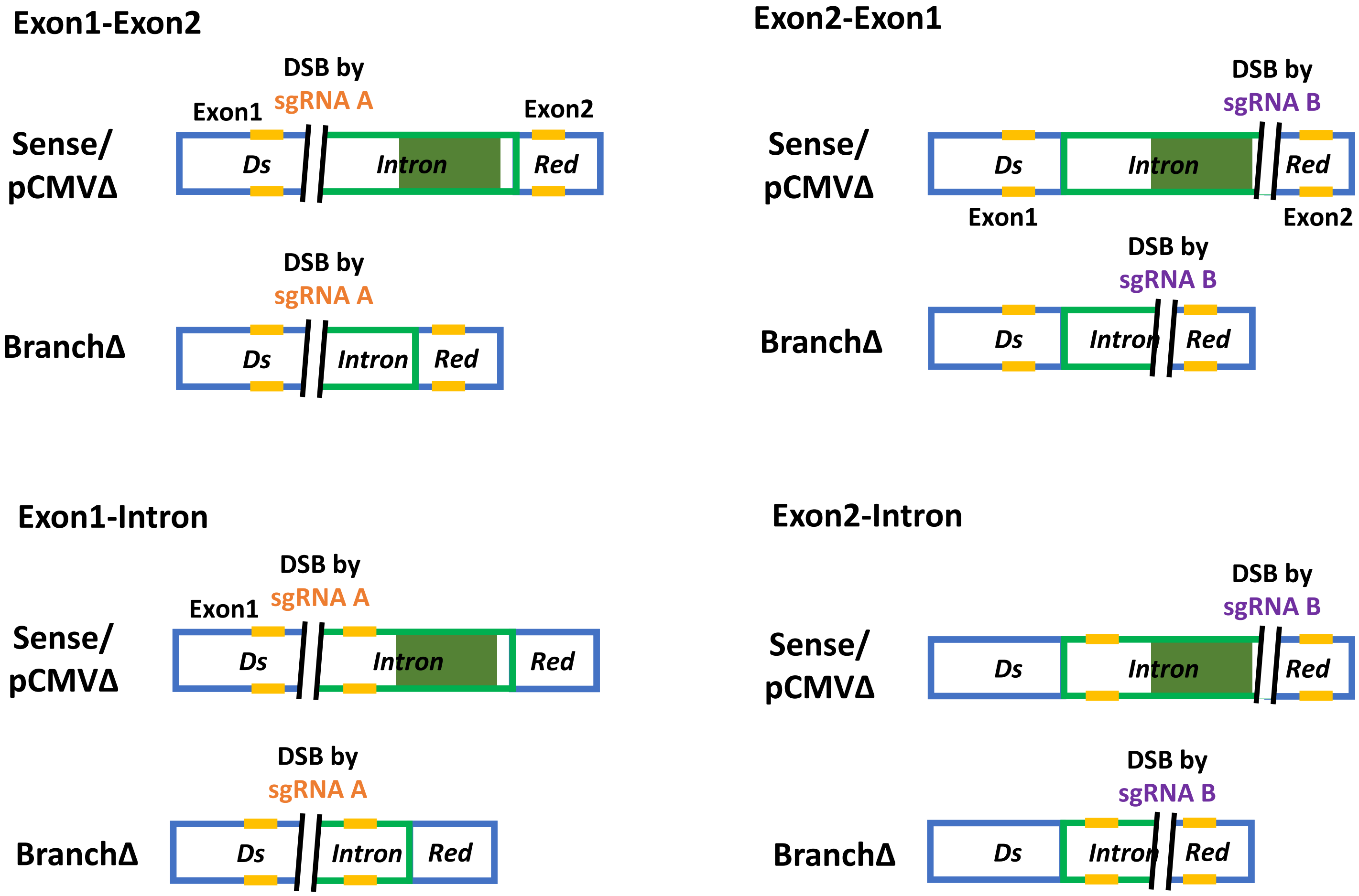

### Extended Data Fig. 9

**b**

Microhomology scheme: Sense/pCMVΔ, sgRNA A, forward strand

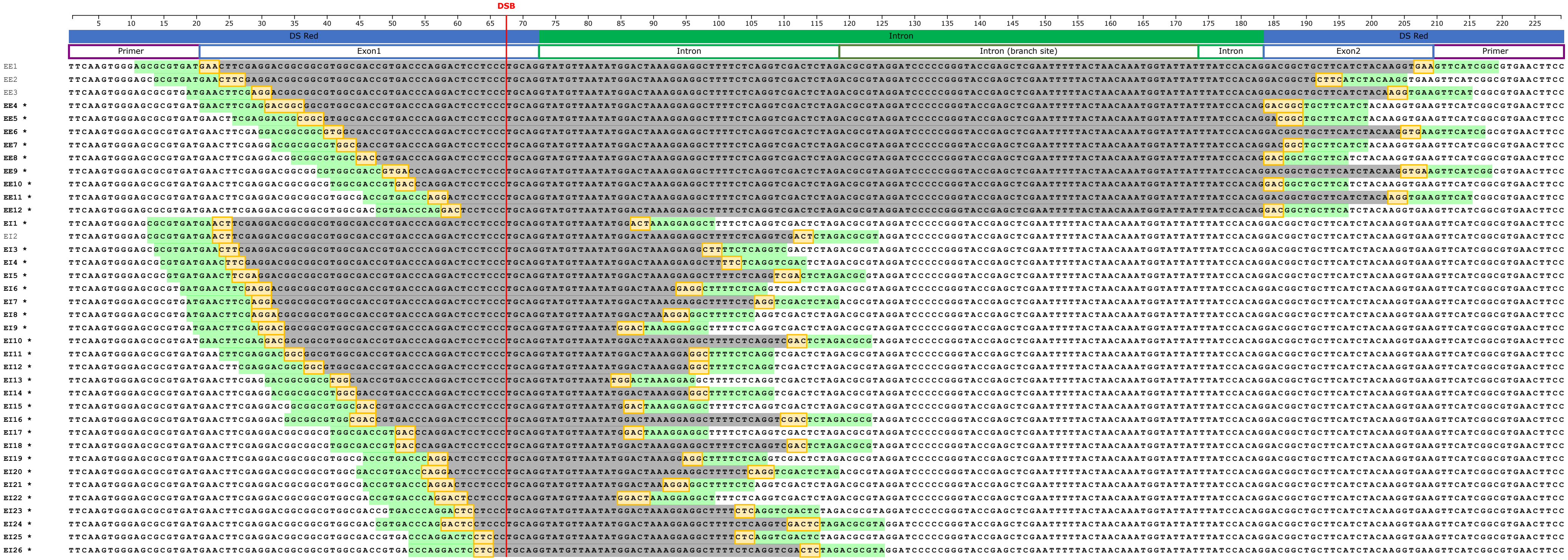


**c**

Microhomology scheme: Sense/pCMVΔ, sgRNA B, reverse strand

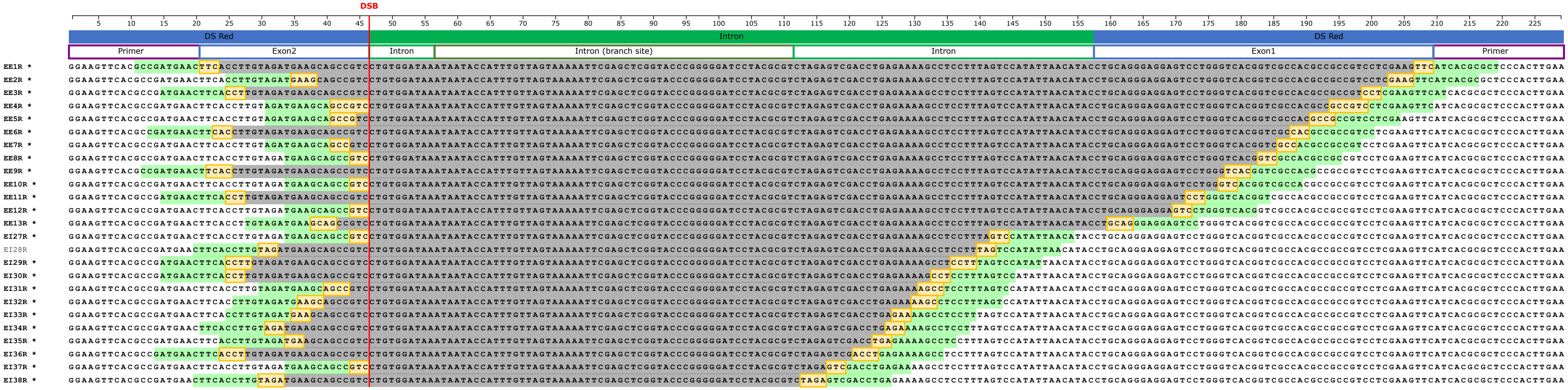

Extended Data Fig. 9

d

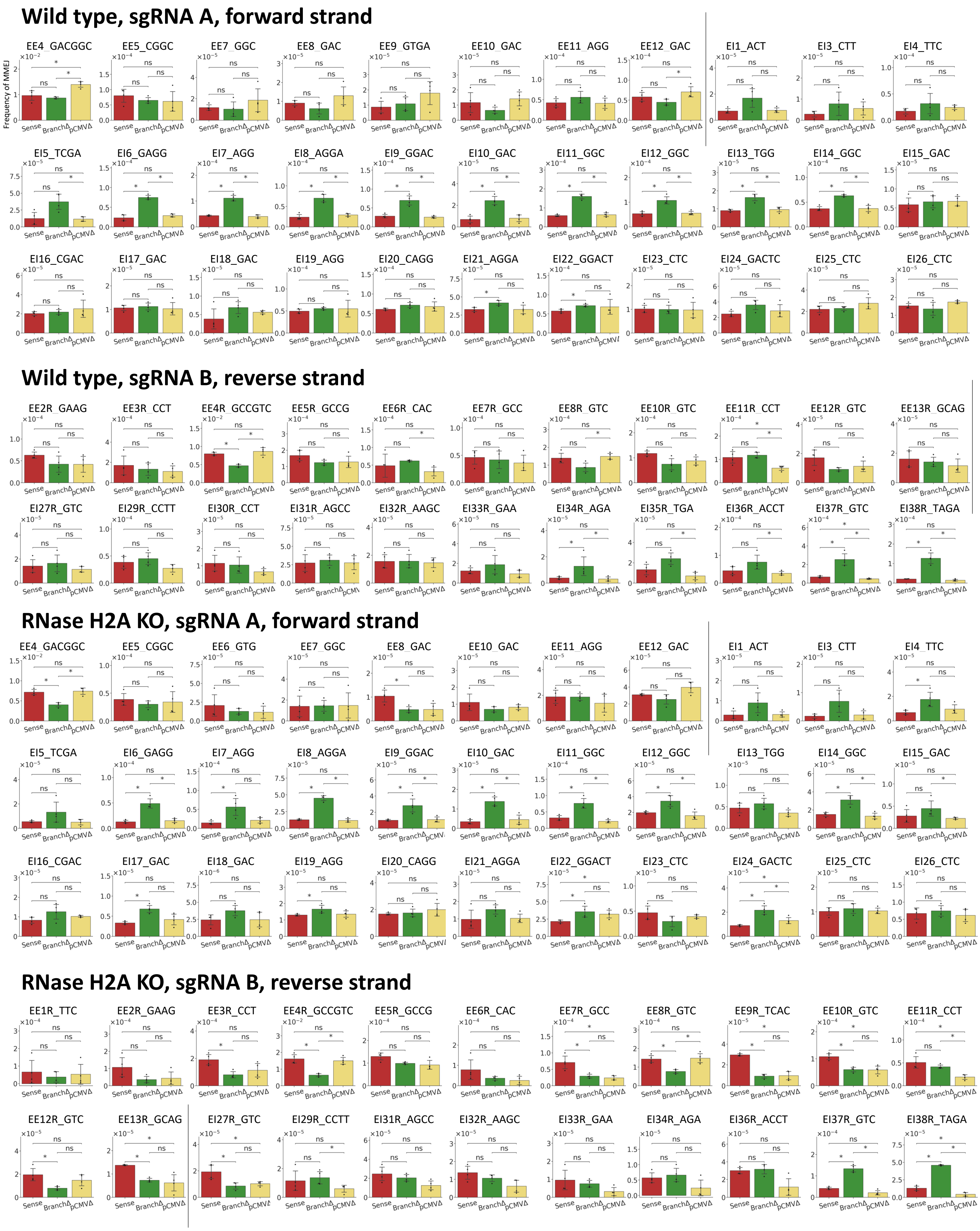

**Extended Data Fig. 9**

**e**

#### HEK-293T wild type

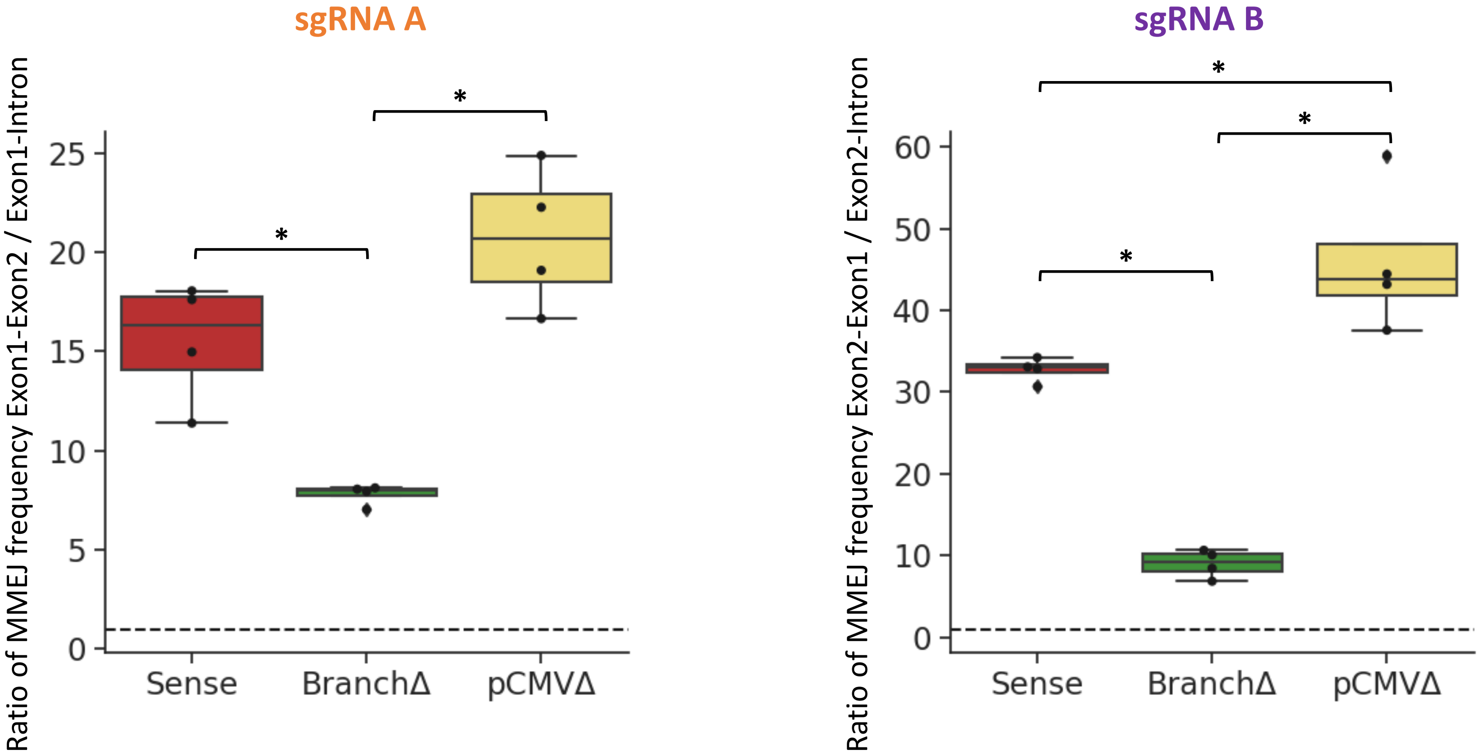


Ratio of MMEJ frequency
[Sum of MMEJ freq using the exon-exon microhomology] / [Sum of MMEJ freq using the exon-intron microhomology]

#### HEK-293T RNase H2A KO

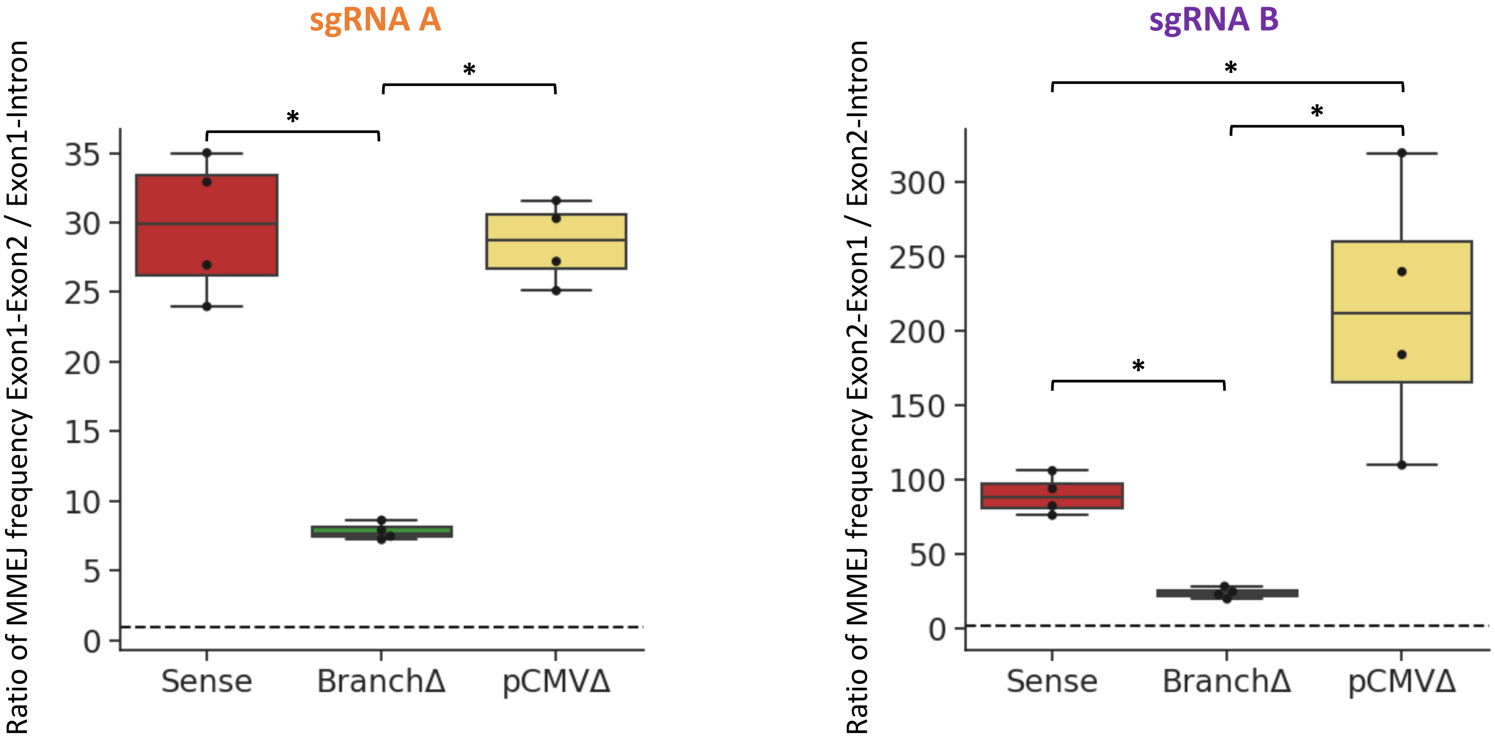


Ratio of MMEJ frequency
[Sum of MMEJ freq using the exon-exon microhomology] / [Sum of MMEJ freq using the exon-intron microhomology]

**Extended Data Figure 9 | The spliced transcript facilitates MMEJ between exon-exon microhomologies, while the non-spliced transcript facilitates MMEJ between exon-intron microhomologies.**

**a,** Schemes of regions in which microhomology pairs were identified within the sequenced area to determine the frequency of MMEJ following a DSB by the sgRNA A (left) or B (right). The microhomology pairs were categorized into two groups: exon-exon (top) and exon-intron (bottom). Yellow lines indicate examples of microhomologies; the black-parallel lines show the DSB site; the dark green box shows the region containing the branch site. **b,** Schemes showing all microhomology pairs 3 bp or longer between the two primer sequences in the Sense and pCMVΔ constructs, sgRNA A, forward strand. The *x*-axis shows nucleotide positions from the 5′-primer sequence. First row: *DsRed* gene exons (solid blue), intron (solid green). Second row: primer sequence (purple outline), Exon1/Exon2 (blue outline), intron (green outline), branch site (dark green outline). DSB position is indicated with a red vertical line. Left margin labels are microhomology IDs (EE, exon-exon; EI, exon-intron); a bold asterisk indicates MMEJ products detected in the sequencing data. Each following row shows the nucleotide sequence of the construct between primer sequences; region deleted by MMEJ repair (gray highlight); microhomology pair (yellow highlight); +/- 10-bp flanking region of microhomology pair used for MMEJ detection in the sequencing data (light green highlight). **c,** As in (b) for Sense and pCMVΔ constructs, sgRNA B, reverse strand. **d,** MMEJ frequencies from each microhomology pair detected in the sequencing libraries following a DSB by the sgRNA A or B in the Sense (red), BranchΔ (green), and pCMVΔ (yellow) constructs of wild-type and RNase H2A KO cells. Individual frequencies are plotted. The ID of the microhomology pair being analyzed is shown on top of each bar graph. Data are mean ± s.d. with 4 biological replicates; N=4. The 'ns' on the bar graphs means a non-significant difference ($P$-value > 0.05, two-tailed Mann-Whitney $U$ test). **e,** Ratio of MMEJ frequency for the exon-exon and exon-intron following a DSB by the sgRNA A or B in the Sense (red), BranchΔ (green), and pCMVΔ (yellow) constructs of wild-type (top) and RNase H2A KO (bottom) cells. The ratio was calculated by dividing the sum of MMEJ frequencies from the exon-exon microhomology pairs by the sum of MMEJ frequencies from the exon-intron microhomology pairs in each sequencing library. The median of the points is shown as the middle line of the box. The first and third quartiles are indicated by the box frames and the whiskers represent the largest point not more than 1.5 interquartile range (IQR) beyond the box frame. All data points outside the whiskers are classified as outliers and shown as diamond points. *, $P$ = 0.029 (two-tailed Mann-Whitney $U$ test).

Extended Data Fig. 10

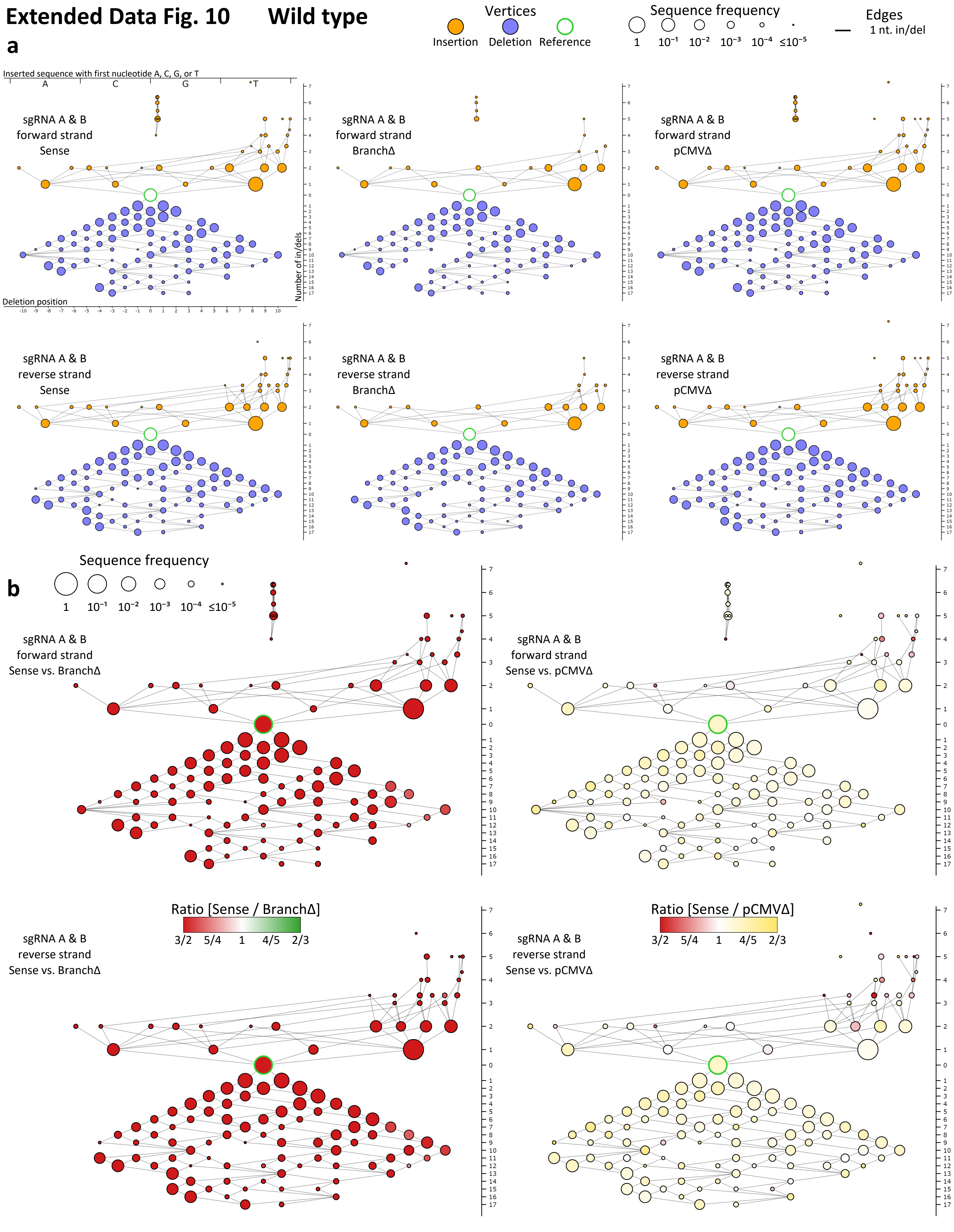

### Extended Data Fig. 10

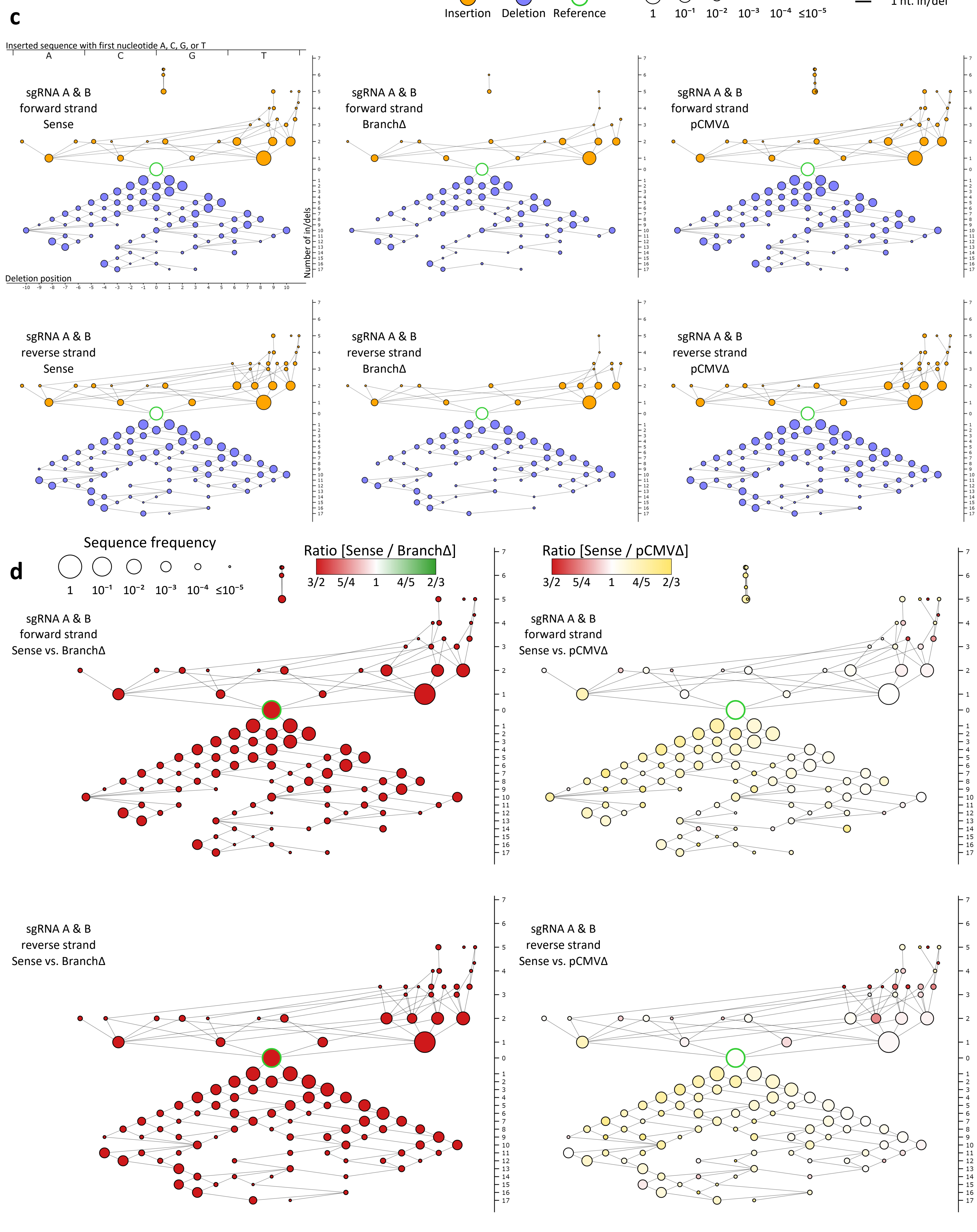

#### Extended Data Fig. 10 Wild type antisense

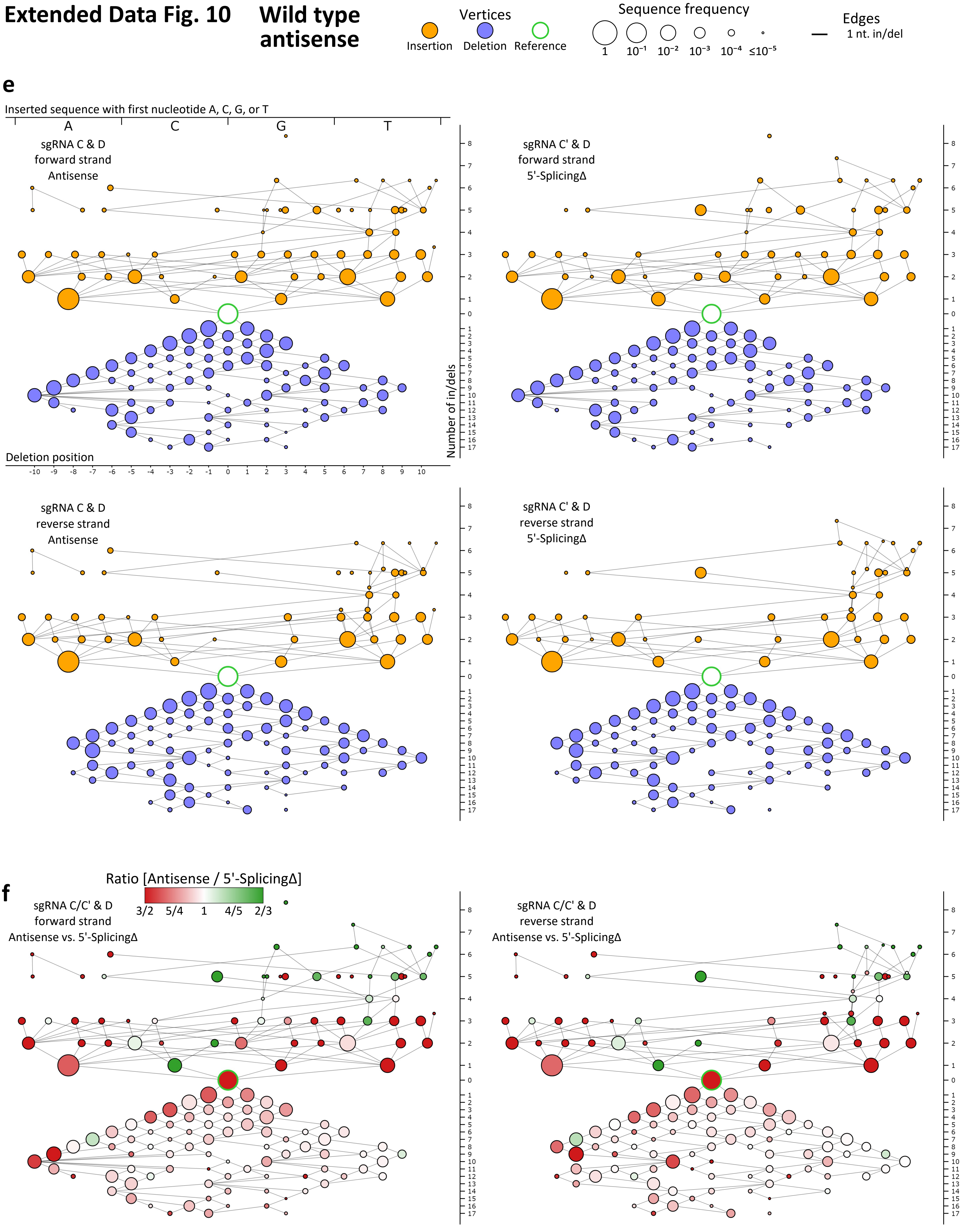

**Extended Data Figure 10 | Transcript RNA promotes gap repair in a sequence-dependent manner via NHEJ – all data for the sense and antisense constructs.**

**a,** Individual variation-distance graphs illustrating sequence variations within DSB-sequence windows observed after gap induction by sgRNA A and sgRNA B in the Sense, BranchΔ, and pCMVΔ constructs of wild-type cells. Data obtained by sequencing the forward (top) and the reverse (bottom) strands. **b,** Comparison variation-distance graphs of the DSB-sequence windows obtained after gap induction by sgRNA A and sgRNA B for the Sense vs. the BranchΔ constructs (left) or for the Sense vs. the pCMVΔ constructs (right) of wild-type cells. The vertices represent the same DSB-sequence windows as for the individual graphs while the vertex colors specify the relative frequency in the Sense (red) vs. the BranchΔ (green) construct, or the Sense (red) vs. the pCMVΔ (yellow) construct; the vertex sizes show the log of the maximum of the two mean frequencies of the corresponding DSB-sequence window in the two compared constructs. Data obtained by sequencing the forward (top) and the reverse (bottom) strands. **c,** Same as in (a) for constructs of RNase H2A KO cells. **d,** Same as in (b) for constructs of RNase H2A KO cells. **e,** Individual variation-distance graphs illustrating sequence variations within DSB-sequence windows observed after gap induction by sgRNA C/C' and sgRNA D in the Antisense and 5′-SplicingΔ constructs of wild-type cells. Data obtained by sequencing the forward (top) and the reverse (bottom) strands. **f,** Comparison variation-distance graphs of the DSB-sequence windows obtained after gap induction by sgRNA C/C' and sgRNA D for the Antisense vs. the 5′-SplicingΔ constructs of wild-type cells. The vertices represent the same DSB-sequence windows as for the individual graphs while the vertex colors specify the relative frequency for the Antisense (red) vs. the 5′-Splicing (green) construct; the vertex sizes show the log of the maximum of the two mean frequencies of the corresponding DSB-sequence window in the two compared constructs. Data obtained by sequencing the forward (left) and the reverse (right) strands. Refer to **Extended Data Figure 7** for the description of variation-distance graphs. For graphs showing reverse-strand data, the sequences are reverse-complemented prior to computing *x*-*y* coordinates so that they correspond to forward strand sequence coordinates.

#### Extended Data Fig. 11

**a**

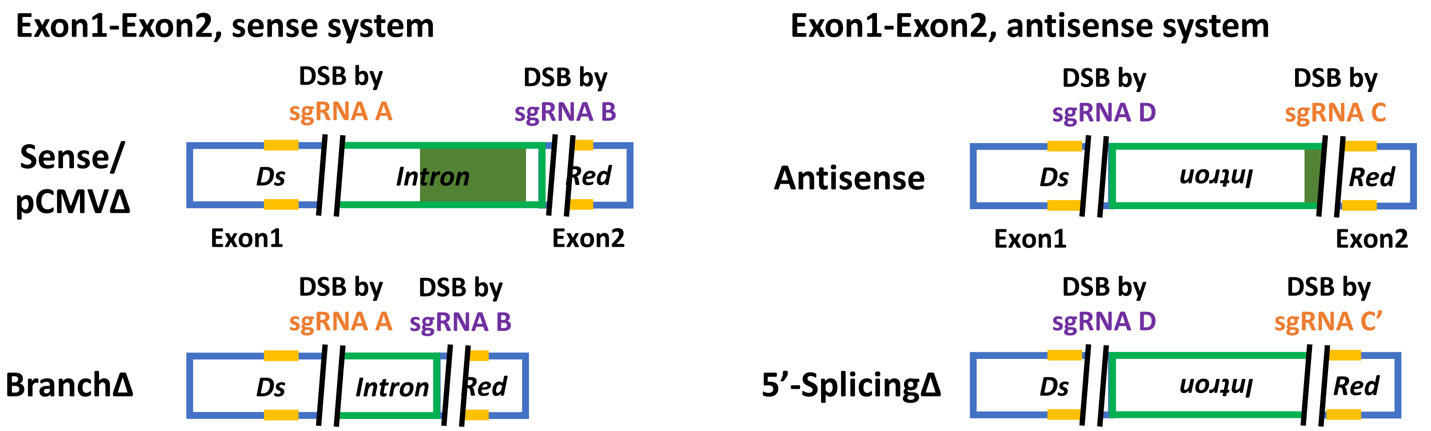

### Extended Data Fig. 11

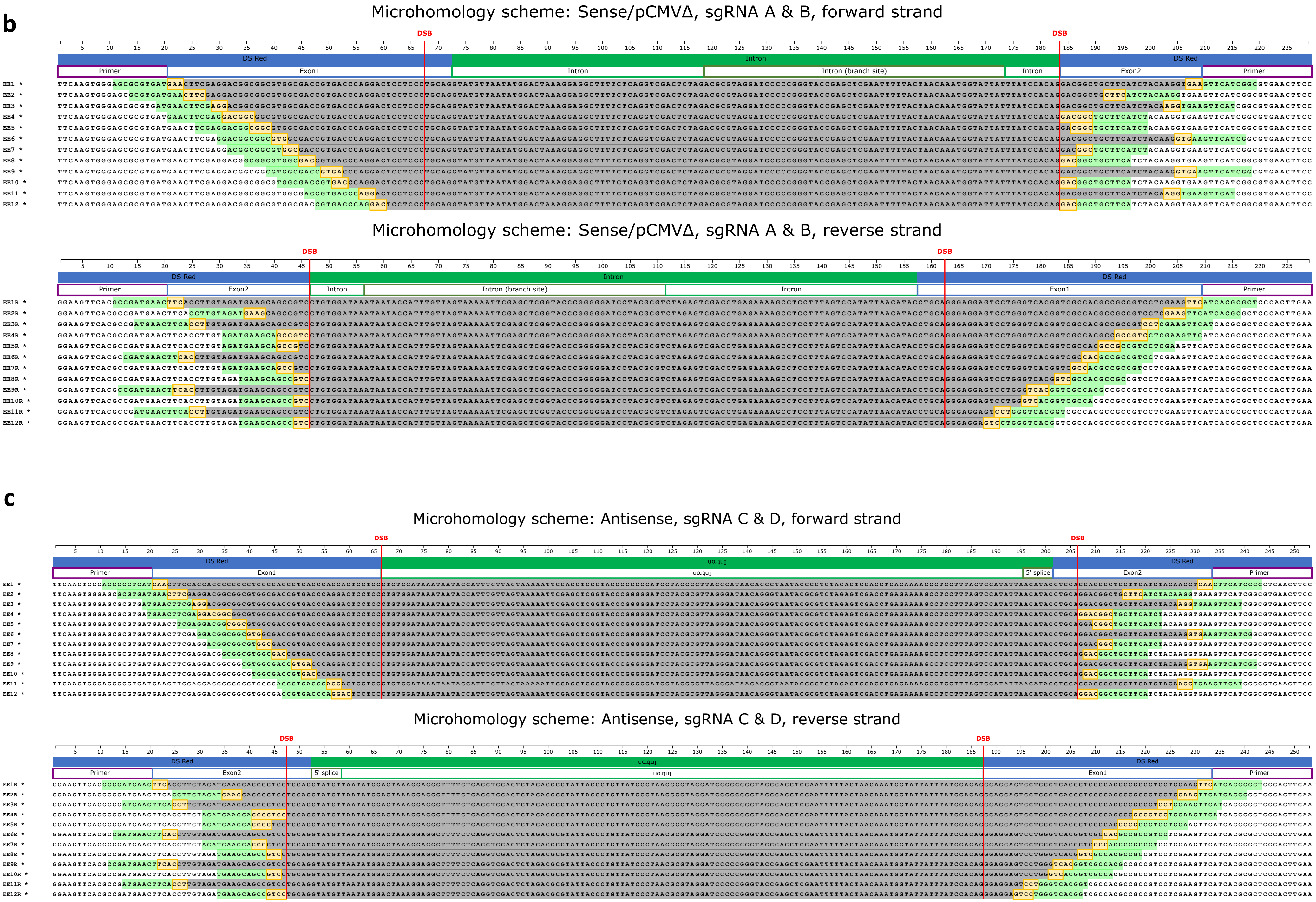

### Extended Data Fig. 11

## d

##### Wild type, sgRNA A and B (2 DSBs), forward strand

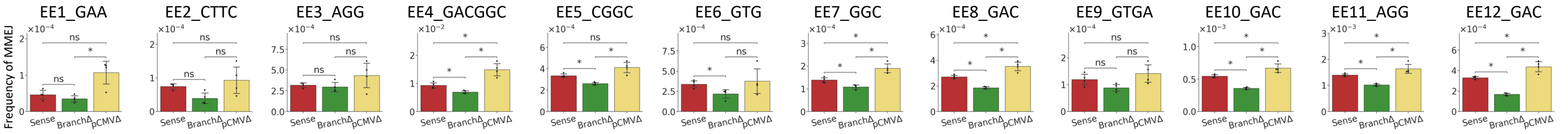


##### Wild type, sgRNA A and B (2 DSBs), reverse strand

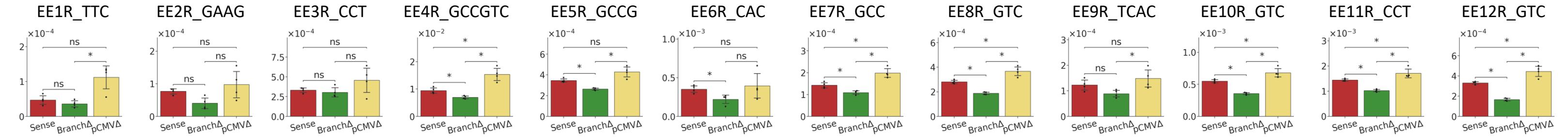


##### RNase H2A KO, sgRNA A and B (2 DSBs), forward strand

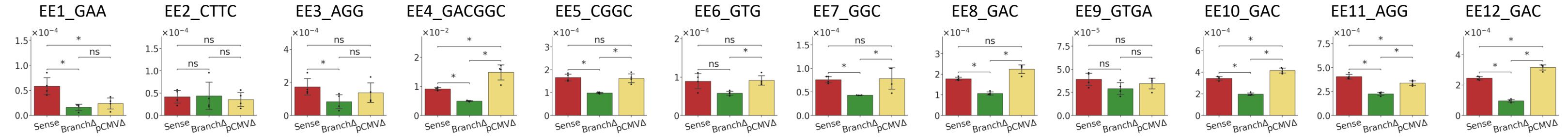


##### RNase H2A KO, sgRNA A and B (2 DSBs), reverse strand

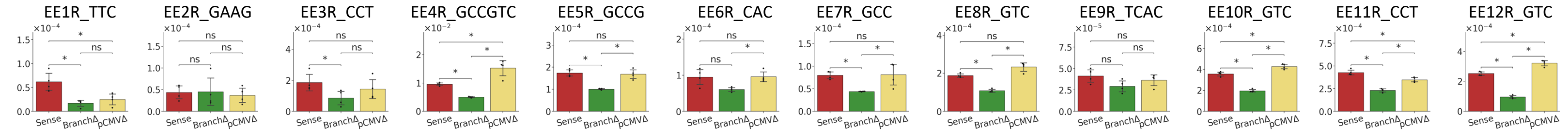


## e

##### Antisense, wild type, sgRNA C (or C') and D (2 DSBs), forward strand

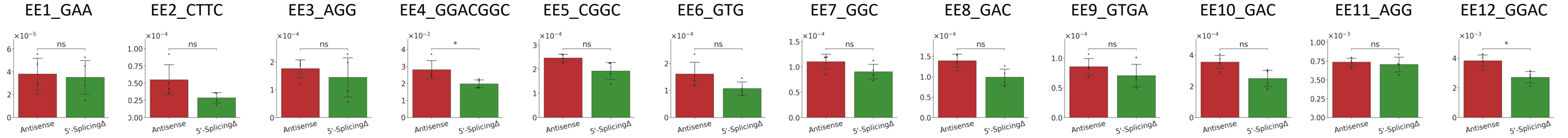


##### Antisense, wild type, sgRNA C (or C') and D (2 DSBs), reverse strand

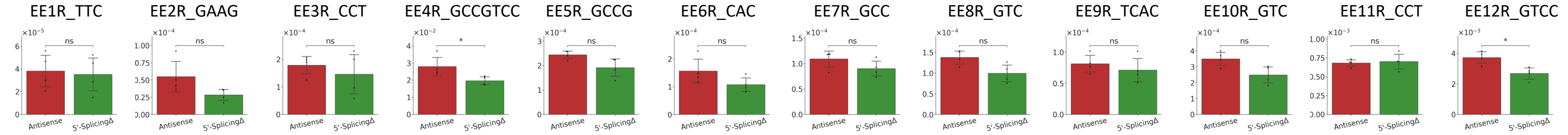

**Extended Data Figure 11 | Transcript RNA promotes gap repair in a sequence-dependent manner via MMEJ – all data for the sense and antisense constructs.**

**a,** Schemes of regions in which Exon1-Exon2 microhomology pairs were identified within the sequenced area to determine the frequency of MMEJ following a gap by the sgRNA A and B in the sense constructs (left), or by the sgRNA C/C' and D in the antisense constructs (right). Yellow lines indicate examples of microhomologies; the black, parallel lines show the DSB sites; the dark, green box shows the region containing the branch site for the Sense and pCMVΔ constructs, and the region with the 5′-splice site for the Antisense construct. **b,** Schemes showing all microhomology pairs 3 bp or longer between the two primer sequences in the Sense and pCMVΔ constructs, sgRNA A and B, forward strand (top), reverse strand (bottom). The x-axis shows nucleotide positions from the 5′-primer sequence. First row: *DsRed* gene exons (solid blue), intron (solid green). Second row: primer sequence (purple outline), Exon1/Exon2 (blue outline), intron (green outline), branch site (dark green outline). DSB positions are indicated with two red vertical lines. Left margin labels are microhomology IDs (EE, exon-exon); bold/asterisk indicates MMEJ products detected in the sequencing data. Each following row shows the nucleotide sequence of the construct between primer sequences; region deleted by MMEJ repair (gray highlight); microhomology pair (yellow highlight); +/- 10-bp flanking region of microhomology pair used for MMEJ detection in the sequencing data (light green highlight). **c,** As in (b) for Antisense construct, sgRNA C and D, forward strand (top), reverse strand (bottom). The reversed intron in the antisense constructs is indicated with an upside-down label; 5′-splice site (dark green outline). **d,** Mean of frequencies of MMEJ repair from each microhomology pair detected in the sequencing libraries following a gap by the sgRNA A and B in the Sense (red), BranchΔ (green), and pCMVΔ (yellow) constructs of wild-type and RNase H2A KO cells, and **e,** following a gap by the sgRNA C/C' and D in the Antisense (red) and 5′-SplicingΔ (green) constructs of wild-type cells. Plotted data are mean ± s.d. with 4 biological replicates indicated with dots. The ID of the microhomology pair being analyzed is on top of each bar graph; N=4. The 'ns' on the bar graphs means a non-significant difference ($P$-value > 0.05, two-tailed Mann-Whitney $U$ test). *, $P$ = 0.029 (two-tailed Mann-Whitney $U$ test).

**Supplementary Table 1. List of oligonucleotides used in this study.**

| Name | Sequence shown in 5′ to 3′ direction | Description |
|---|---|---|
| YJo 11 | ACACTCTTTCCCTACACGACGCTCTTCCGATCTNNAGTCTTCAAGTGGGAGCGCGTGA*T | Forward primers for the 1st PCR of the NGS library prep |
| YJo 12 | ACACTCTTTCCCTACACGACGCTCTTCCGATCTNNCATGTTCAAGTGGGAGCGCGTGA*T | |
| YJo 13 | ACACTCTTTCCCTACACGACGCTCTTCCGATCTNNGCTATTCAAGTGGGAGCGCGTGA*T | |
| YJo 14 | ACACTCTTTCCCTACACGACGCTCTTCCGATCTNNTAGCTTCAAGTGGGAGCGCGTGA*T | |
| YJo 15 | GTGACTGGAGTTCAGACGTGTGCTCTTCCGATCTNNAGGCGGAAGTTCACGCCGATGAA*C | Reverse primers for the 1st PCR of the NGS library prep |
| YJo 16 | GTGACTGGAGTTCAGACGTGTGCTCTTCCGATCTNNTACTGGAAGTTCACGCCGATGAA*C | |
| YJo 17 | GTGACTGGAGTTCAGACGTGTGCTCTTCCGATCTNNAGCGGGAAGTTCACGCCGATGAA*C | |
| YJo 18 | GTGACTGGAGTTCAGACGTGTGCTCTTCCGATCTNNTGAGGGAAGTTCACGCCGATGAA*C | |
| YJo 19 | GTGACTGGAGTTCAGACGTGTGCTCTTCCGATCTNNGCAGGGAAGTTCACGCCGATGAA*C | |
| YJo 20 | GTGACTGGAGTTCAGACGTGTGCTCTTCCGATCTNNGAGCGGAAGTTCACGCCGATGAA*C | |
| YJo 21 | GTGACTGGAGTTCAGACGTGTGCTCTTCCGATCTNNATCTGGAAGTTCACGCCGATGAA*C | |
| YJo 22 | GTGACTGGAGTTCAGACGTGTGCTCTTCCGATCTNNCTCGGGAAGTTCACGCCGATGAA*C | |
| YJo 23 | GTGACTGGAGTTCAGACGTGTGCTCTTCCGATCTNNGTGTGGAAGTTCACGCCGATGAA*C | |
| YJo 24 | GTGACTGGAGTTCAGACGTGTGCTCTTCCGATCTNNAGATGGAAGTTCACGCCGATGAA*C | |
| YJo 25 | GTGACTGGAGTTCAGACGTGTGCTCTTCCGATCTNNCGCAGGAAGTTCACGCCGATGAA*C | |
| YJo 26 | GTGACTGGAGTTCAGACGTGTGCTCTTCCGATCTNNTGTCGGAAGTTCACGCCGATGAA*C | |
| D501 | AATGATACGGCGACCACCGAGATCTACACTATAGCCTACACTCTTTCCCTACACGAC | Forward primers for the 2nd PCR of the NGS library prep |
| D502 | AATGATACGGCGACCACCGAGATCTACACATAGAGGCACACTCTTTCCCTACACGAC | |
| D503 | AATGATACGGCGACCACCGAGATCTACACCCTATCCTACACTCTTTCCCTACACGAC | |
| D504 | AATGATACGGCGACCACCGAGATCTACACGGCTCTGAACACTCTTTCCCTACACGAC | |
| D505 | AATGATACGGCGACCACCGAGATCTACACAGGCGAAGACACTCTTTCCCTACACGAC | |
| D506 | AATGATACGGCGACCACCGAGATCTACACTAATCTTAACACTCTTTCCCTACACGAC | |
| D507 | AATGATACGGCGACCACCGAGATCTACACCAGGACGTACACTCTTTCCCTACACGAC | |
| D508 | AATGATACGGCGACCACCGAGATCTACACGTACTGACACACTCTTTCCCTACACGAC | |

| | | |
|---|---|---|
| D701 | CAAGCAGAAGACGGCATACGAGATCGAGTAATGTGACTGGAGTTCAGACGTGT | Reverse primers for the 2nd PCR of the NGS library prep |
| D702 | CAAGCAGAAGACGGCATACGAGATTCTCCGGAGTGACTGGAGTTCAGACGTGT | |
| D705 | CAAGCAGAAGACGGCATACGAGATTTCTGAATGTGACTGGAGTTCAGACGTGT | |
| D707 | CAAGCAGAAGACGGCATACGAGATAGCTTCAGGTGACTGGAGTTCAGACGTGT | |
| D712 | CAAGCAGAAGACGGCATACGAGATCTATCGCTGTGACTGGAGTTCAGACGTGT | |
| DsRed.pCis.F3 | CTTCGCCTGGGACATCCTGT | PCR primer amplifying part of the *DsRed* gene |
| DsRed.pCis.F8 | AGCTGAAGGTGACCAAGGGC | PCR primer amplifying part of the *DsRed* gene |
| DsRed.pCis.R3 | TCCTTCAGCTTCAGGGCCTT | PCR primer amplifying part of the *DsRed* gene |
| 55bpDel.AI.F | TCAGGTCGACTCTAGTTATCCACAGCTGCAGGACG | Primers for deleting the branch site through in-fusion |
| 55bpDel.AI.R | CTAGAGTCGACCTGAGAAAA | |
| YJo 1 | AATGATACGGCGACCACCGAGATCTACACTATAGCCTACACTCTTTCCCTACACGACGCTCTTCCGATCTTGATGAACTTCGAGGACGG*C | Forward primers for the 2nd PCR of the NGS library prep (antisense system) |
| YJo 9 | AATGATACGGCGACCACCGAGATCTACACAGGCGAAGACACTCTTTCCCTACACGACGCTCTTCCGATCTTGATGAACTTCGAGGACGG*C | |
| YJo 10 | AATGATACGGCGACCACCGAGATCTACACTAATCTTAACACTCTTTCCCTACACGACGCTCTTCCGATCTTGATGAACTTCGAGGACGG*C | |
| YJo 4 | CAAGCAGAAGACGGCATACGAGATCGAGTAATGTGACTGGAGTTCAGACGTGTGCTCTTCCGATCTGGAAGTTCACGCCGATGAA*C | Reverse primers for the 2nd PCR of the NGS library prep (antisense system) |
| YJo 5 | CAAGCAGAAGACGGCATACGAGATTCTCCGGAGTGACTGGAGTTCAGACGTGTGCTCTTCCGATCTGGAAGTTCACGCCGATGAA*C | |
| YJo 6 | CAAGCAGAAGACGGCATACGAGATAATGAGCGGTGACTGGAGTTCAGACGTGTGCTCTTCCGATCTGGAAGTTCACGCCGATGAA*C | |
| YJo 8 | CAAGCAGAAGACGGCATACGAGATGGAATCTCGTGACTGGAGTTCAGACGTGTGCTCTTCCGATCTGGAAGTTCACGCCGATGAA*C | |
| YJo 11 | CAAGCAGAAGACGGCATACGAGATTTCTGAATGTGACTGGAGTTCAGACGTGTGCTCTTCCGATCTGGAAGTTCACGCCGATGAA*C | |

| | | |
|---|---|---|
| RNASEH2A.F | CGAGACCCGCTCCTGCAGTATT | RNASEH2A sequencing primer |
| RNASEH2A.R | CGCTCTCCAATAGGGTCTTTGA | RNASEH2A sequencing primer |
| Cy5.3PS.rG | /Cy5/G*G*A*TCCGGTAGTrGTTAGGCCTG*A*A*C | Used for testing RNase H2 activity |
| Cy5.3PS | /Cy5/G*G*A*TCCGGTAGTGTTAGGCCTG*A*A*C | |
| DNA.comp.3PS | G*T*T*CAGGCCTAACACTACCGGA*T*C*C | |

Names of oligonucleotides used in this study are shown with their sequence in the 5′ to 3′ direction, and corresponding description. *, Phosphorothioate bond; /Cy5/, Cy5 dye; r, ribonucleotide.

**Supplementary Table 2. List of synthesized sgRNAs used in this study.**

| Name | Sequence | Description |
|---|---|---|
| sgRNA A | mU*mA*mU*rUrArArCrArUrArCrCrUrGrCrArGrGrGrGrUrUrUrUrArGrArGrCrUrArGrArArArUrArGrCrArArGrUrUrArArArArUrArArGrGrCrUrArGrUrCrCrGrUrUrArUrCrArArCrUrUrGrArArArArArGrUrGrGrCrArCrCrGrArGrUrCrGrGrUrGrCmU*mU*mU*rU | Synthesized sgRNA targeting the sense plasmids |
| sgRNA B | mA*mG*mA*rUrGrArArGrCrArGrCrCrGrUrCrCrUrGrGrUrUrUrUrArGrArGrCrUrArGrArArArUrArGrCrArArGrUrUrArArArArUrArArGrGrCrUrArGrUrCrCrGrUrUrArUrCrArArCrUrUrGrArArArArArGrUrGrGrCrArCrCrGrArGrUrCrGrGrUrGrCmU*mU*mU*rU | Synthesized sgRNA targeting the sense plasmids |
| sgRNA C | mU*mA*mU*rUrArArCrArUrArCrCrUrGrCrArGrGrArGrUrUrUrUrArGrArGrCrUrArGrArArArUrArGrCrArArGrUrUrArArArArUrArArGrGrCrUrArGrUrCrCrGrUrUrArUrCrArArCrUrUrGrArArArArArGrUrGrGrCrArCrCrGrArGrUrCrGrGrUrGrCmU*mU*mU*rU | Synthesized sgRNA targeting the antisense plasmids |
| sgRNA C’ | mA*mG*mU*rCrCrArUrArUrUrArCrUrGrCrArGrGrArGrUrUrUrUrArGrArGrCrUrArGrArArArUrArGrCrArArGrUrUrArArArArUrArArGrGrCrUrArGrUrCrCrGrUrUrArUrCrArArCrUrUrGrArArArArArGrUrGrGrCrArCrCrGrArGrUrCrGrGrUrGrCmU*mU*mU*rU | Synthesized sgRNA targeting the 5′-SplicingΔ plasmid |
| sgRNA D | mG*mA*mC*rCrCrArGrGrArCrUrCrCrUrCrCrCrUrGrGrUrUrUrUrArGrArGrCrUrArGrArArArUrArGrCrArArGrUrUrArArArArUrArArGrGrCrUrArGrUrCrCrGrUrUrArUrCrArArCrUrUrGrArArArArArGrUrGrGrCrArCrCrGrArGrUrCrGrGrUrGrCmU*mU*mU*rU | Synthesized sgRNA targeting the antisense plasmids |

The sgRNAs used in this study, are shown with their names, sequences, and description. The m is a chemical modification in a nucleotide provided by Integrated DNA Technologies for the Alt-R CRISPR-Cas9 sgRNA products. The r means a ribonucleotide. *, Phosphorothioate bond.

**Supplementary Table 3.** ***P*****-values comparing frequencies of RNA-mediated DNA repair between the HEK-293T RNase H2 wild-type and KO cells by permutation test.**

| DSB | Sequencing Strand | DSB repair | Ratio | *P*-value |
|---|---|---|---|---|
| 1 DSB, sgRNA A | Forward | NHEJ | 1.148 | 0.319 |
| 1 DSB, sgRNA A | Reverse | R-TDR | 0.496 | 0.008* |
| 1 DSB, sgRNA B | Reverse | NHEJ | 1.047 | 0.437 |
| 1 DSB, sgRNA B | Reverse | R-TDR | 1.341 | 0.667 |
| 2 DSBs | Forward | NHEJ | 0.758 | 0.067 |
| 2 DSBs | Forward | Flipped intron | 1.385 | 0.983 |
| 2 DSBs | Reverse | NHEJ | 0.750 | 0.049* |
| 1 DSB, sgRNA A | Forward | MMEJ_EE1 | 2.829 | 0.736 |
| 1 DSB, sgRNA A | Forward | MMEJ_EE2 | 0.472 | 0.284 |
| 1 DSB, sgRNA A | Forward | MMEJ_EE3 | 0.393 | 0.073 |
| 1 DSB, sgRNA A | Forward | MMEJ_EE4 | 0.624 | 0.081 |
| 1 DSB, sgRNA A | Forward | MMEJ_EE5 | 0.954 | 0.464 |
| 1 DSB, sgRNA A | Forward | MMEJ_EE6 | 0.242 | 0.026* |
| 1 DSB, sgRNA A | Forward | MMEJ_EE7 | 1.179 | 0.602 |
| 1 DSB, sgRNA A | Forward | MMEJ_EE8 | 0.673 | 0.143 |
| 1 DSB, sgRNA A | Forward | MMEJ_EE9 | 0.232 | 0.050 |
| 1 DSB, sgRNA A | Forward | MMEJ_EE10 | 1.106 | 0.589 |
| 1 DSB, sgRNA A | Forward | MMEJ_EE11 | 0.753 | 0.317 |
| 1 DSB, sgRNA A | Forward | MMEJ_EE12 | 1.067 | 0.569 |
| 1 DSB, sgRNA A | Forward | MMEJ_EI1 | 1.251 | 0.639 |
| 1 DSB, sgRNA A | Forward | MMEJ_EI2 | 0.442 | 0.249 |
| 1 DSB, sgRNA A | Forward | MMEJ_EI3 | 1.528 | 0.734 |
| 1 DSB, sgRNA A | Forward | MMEJ_EI4 | 1.371 | 0.672 |
| 1 DSB, sgRNA A | Forward | MMEJ_EI5 | 0.825 | 0.409 |
| 1 DSB, sgRNA A | Forward | MMEJ_EI6 | 1.164 | 0.661 |
| 1 DSB, sgRNA A | Forward | MMEJ_EI7 | 1.506 | 0.792 |
| 1 DSB, sgRNA A | Forward | MMEJ_EI8 | 1.194 | 0.671 |
| 1 DSB, sgRNA A | Forward | MMEJ_EI9 | 1.155 | 0.603 |
| 1 DSB, sgRNA A | Forward | MMEJ_EI10 | 1.157 | 0.604 |
| 1 DSB, sgRNA A | Forward | MMEJ_EI11 | 0.886 | 0.374 |
| 1 DSB, sgRNA A | Forward | MMEJ_EI12 | 0.862 | 0.390 |
| 1 DSB, sgRNA A | Forward | MMEJ_EI13 | 0.675 | 0.205 |
| 1 DSB, sgRNA A | Forward | MMEJ_EI14 | 1.192 | 0.665 |
| 1 DSB, sgRNA A | Forward | MMEJ_EI15 | 1.406 | 0.749 |
| 1 DSB, sgRNA A | Forward | MMEJ_EI16 | 1.417 | 0.790 |
| 1 DSB, sgRNA A | Forward | MMEJ_EI17 | 1.919 | 0.914 |
| 1 DSB, sgRNA A | Forward | MMEJ_EI18 | 0.866 | 0.441 |
| 1 DSB, sgRNA A | Forward | MMEJ_EI19 | 1.146 | 0.642 |
| 1 DSB, sgRNA A | Forward | MMEJ_EI20 | 0.894 | 0.369 |
| 1 DSB, sgRNA A | Forward | MMEJ_EI21 | 1.222 | 0.639 |
| 1 DSB, sgRNA A | Forward | MMEJ_EI22 | 1.327 | 0.728 |
| 1 DSB, sgRNA A | Forward | MMEJ_EI23 | 0.649 | 0.226 |
| 1 DSB, sgRNA A | Forward | MMEJ_EI24 | 1.658 | 0.870 |
| 1 DSB, sgRNA A | Forward | MMEJ_EI25 | 1.075 | 0.571 |
| 1 DSB, sgRNA A | Forward | MMEJ_EI26 | 1.250 | 0.693 |
| 1 DSB, sgRNA B | Reverse | MMEJ_EE1R | 0.836 | 0.415 |

| | | | | |
|---|---|---|---|---|
| 1 DSB, sgRNA B | Reverse | MMEJ_EE2R | 0.501 | 0.110 |
| 1 DSB, sgRNA B | Reverse | MMEJ_EE3R | 0.538 | 0.108 |
| 1 DSB, sgRNA B | Reverse | MMEJ_EE4R | 0.674 | 0.116 |
| 1 DSB, sgRNA B | Reverse | MMEJ_EE5R | 1.037 | 0.571 |
| 1 DSB, sgRNA B | Reverse | MMEJ_EE6R | 0.366 | 0.039 |
| 1 DSB, sgRNA B | Reverse | MMEJ_EE7R | 0.460 | 0.025 |
| 1 DSB, sgRNA B | Reverse | MMEJ_EE8R | 0.855 | 0.253 |
| 1 DSB, sgRNA B | Reverse | MMEJ_EE9R | 0.319 | 0.003* |
| 1 DSB, sgRNA B | Reverse | MMEJ_EE10R | 0.882 | 0.326 |
| 1 DSB, sgRNA B | Reverse | MMEJ_EE11R | 0.745 | 0.286 |
| 1 DSB, sgRNA B | Reverse | MMEJ_EE12R | 0.750 | 0.141 |
| 1 DSB, sgRNA B | Reverse | MMEJ_EE13R | 0.608 | 0.083 |
| 1 DSB, sgRNA B | Reverse | MMEJ_EI27R | 0.386 | 0.018 |
| 1 DSB, sgRNA B | Reverse | MMEJ_EI28R | 1.127 | 0.531 |
| 1 DSB, sgRNA B | Reverse | MMEJ_EI29R | 1.008 | 0.505 |
| 1 DSB, sgRNA B | Reverse | MMEJ_EI30R | 1.089 | 0.542 |
| 1 DSB, sgRNA B | Reverse | MMEJ_EI31R | 0.730 | 0.201 |
| 1 DSB, sgRNA B | Reverse | MMEJ_EI32R | 0.772 | 0.239 |
| 1 DSB, sgRNA B | Reverse | MMEJ_EI33R | 0.523 | 0.160 |
| 1 DSB, sgRNA B | Reverse | MMEJ_EI34R | 0.349 | 0.011* |
| 1 DSB, sgRNA B | Reverse | MMEJ_EI35R | 0.352 | 0.010* |
| 1 DSB, sgRNA B | Reverse | MMEJ_EI36R | 0.625 | 0.185 |
| 1 DSB, sgRNA B | Reverse | MMEJ_EI37R | 0.839 | 0.326 |
| 1 DSB, sgRNA B | Reverse | MMEJ_EI38R | 0.597 | 0.133 |
| 2 DSBs | Forward | MMEJ_EE1 | 0.368 | 0.007 |
| 2 DSBs | Forward | MMEJ_EE2 | 1.985 | 0.894 |
| 2 DSBs | Forward | MMEJ_EE3 | 0.515 | 0.119 |
| 2 DSBs | Forward | MMEJ_EE4 | 0.724 | 0.012* |
| 2 DSBs | Forward | MMEJ_EE5 | 0.763 | 0.297 |
| 2 DSBs | Forward | MMEJ_EE6 | 1.012 | 0.506 |
| 2 DSBs | Forward | MMEJ_EE7 | 0.722 | 0.228 |
| 2 DSBs | Forward | MMEJ_EE8 | 0.877 | 0.311 |
| 2 DSBs | Forward | MMEJ_EE9 | 1.001 | 0.501 |
| 2 DSBs | Forward | MMEJ_EE10 | 0.873 | 0.315 |
| 2 DSBs | Forward | MMEJ_EE11 | 0.765 | 0.316 |
| 2 DSBs | Forward | MMEJ_EE12 | 0.775 | 0.156 |
| 2 DSBs | Reverse | MMEJ_EE1R | 0.353 | 0.005* |
| 2 DSBs | Reverse | MMEJ_EE2R | 1.965 | 0.890 |
| 2 DSBs | Reverse | MMEJ_EE3R | 0.502 | 0.107 |
| 2 DSBs | Reverse | MMEJ_EE4R | 0.715 | 0.008 |
| 2 DSBs | Reverse | MMEJ_EE5R | 0.759 | 0.293 |
| 2 DSBs | Reverse | MMEJ_EE6R | 1.013 | 0.506 |
| 2 DSBs | Reverse | MMEJ_EE7R | 0.716 | 0.211 |
| 2 DSBs | Reverse | MMEJ_EE8R | 0.863 | 0.296 |
| 2 DSBs | Reverse | MMEJ_EE9R | 0.979 | 0.483 |
| 2 DSBs | Reverse | MMEJ_EE10R | 0.850 | 0.296 |
| 2 DSBs | Reverse | MMEJ_EE11R | 0.763 | 0.316 |
| 2 DSBs | Reverse | MMEJ_EE12R | 0.763 | 0.135 |

The 2 DSBs are generated by sgRNA A and sgRNA B. Permutation tests were performed to compare the ratio of Sense/BranchΔ frequency between RNase H2A wild-type and KO libraries for each of the repair mechanisms, see Methods. * *P*-value < 0.05.

#### Methods

##### Constructs to study DSB repair mediated by RNA in human cells

To study DSB repair mediated by transcript RNA in human cells, we built constructs expressing RNA transcripts varying in sequence and transcription level, three 'sense' constructs: Sense, BranchΔ, and pCMVΔ, and two 'antisense' constructs: Antisense and 5′-SplicingΔ, in which the transcribed RNA is the sense or the antisense RNA of the *DsRed* gene, respectively. An artificial intron was PCR amplified from the plasmid DNA of pSM50 (provided by Dr. Garfinkel's lab at University of Georgia) attaching the PstI restriction enzyme site at its ends, purified by spin column (QIAquick PCR Purification Kit, Qiagen), and was introduced into the *DsRed* gene of the pDsRed plasmid (Addgene, #54493) in its SbfI site, which is compatible to the PstI site, in the sense orientation to make the Sense construct. The ligated plasmid was transformed into *Escherichia coli* cells (Agilent, XL1-Blue Competent Cells). Plasmids were isolated from the *E. coli* colonies grown on kanamycin (40 μg/ml) contained LB plate by Miniprep (GeneJET Plasmid Miniprep Kit, Thermo Scientific). Each of the plasmids containing the sense or antisense orientation of the artificial intron was confirmed by Sanger sequence analysis (Eurofins Genomics). There was a residual 6 bp of repeated sequence at the junction of the exon-intron inserted during the restriction enzyme cloning process. The 6 bp located next to the 3′-splicing site of the intron was deleted from both of sense and antisense plasmids by the *in-vitro* mutagenesis (QuikChange II Site-Directed Mutagenesis kit, Agilent) to reconstitute the original sequence of the *DsRed* gene.

Construction of the antisense constructs. The plasmid construction having the antisense orientation of the artificial intron in the *DsRed* gene was digested by PciI and SspI restriction enzymes (New England BioLabs). The broken ends were filled in by the Klenow fragment (New England BioLabs) to generate blunt ends. Then, the construct was purified by gel extraction (GeneJET Gel Extraction Kit, Thermo Scientific) to isolate the fragment having the *pCMV* and the *DsRed* gene with the antisense intron, and inserted in a vector, the pEGFP-17 (Addgene, #62043) to construct the pSRAIDC. To construct the pSRAIDC, the pEGFP-17 vector was digested by HindIII and BglII restriction enzymes (New England BioLabs), treated by the Klenow fragment, and dephosphorylated with calf intestinal alkaline phosphatase (CIP, New England BioLabs) to prevent self-ligation of the vector. This dephosphorylated vector was purified by spin column and ligated with the insert containing the *pCMV* and the *DsRed* gene with the antisense intron by T4 DNA ligase. The cutting site for the I-SceI-homing endonuclease, TAGGGATAACAGGGTAAT, was inserted in the middle of the intron after a digestion by MluI restriction enzyme (New England BioLabs) and ligation with a short dsDNA oligonucleotide containing the I-SceI restriction enzyme site flanked with MluI restriction enzyme sites by the T4 DNA ligase. The *EF1α* promoter (*EF1α-2* promoter used in Zheng *et al,* Int. J. Med. Sci 2014[1]) was amplified by PCR from a plasmid (customized order#18ACPG6P_2379654_EF1a2, Thermo Fisher Scientific) flanked with the AvrII restriction enzyme site to be inserted into the antisense plasmid to drive transcription of the *DsRed* antisense transcript. Both the PCR fragment containing the *EF1α* promoter and the antisense plasmid were digested by AvrII restriction enzyme (New England BioLabs) and ligated by the T4 DNA ligase to merge them.

The Sense and Antisense constructs were then engineered to contain a protospacer adjacent motif (PAM) sequence in the intron to induce a DSB by sgRNA B for the Sense and sgRNA D for the Antisense (**Extended Data Figure 1**) at the junction between the exon and the intron. The 2 bp in the intron (TA in the 1126-1127 position of the sense plasmid, and in the 1493-1494 position of the antisense plasmid) was substituted to GG by *in vitro* mutagenesis (QuikChange II Site-Directed Mutagenesis Kit, Agilent). We used an existing XGG sequence for the PAM site for the DSB by sgRNA A for the Sense and sgRNA C for the Antisense (**Extended Data Figure 1**).

The 55-bp deletion containing the branch site for the splicing mutant of the sense system (BranchΔ) was done by in-fusion reaction (In-Fusion HD Cloning Kit, Clontech). Primers 55bpDel.AI.F and 55bpDel.AI.R (**Extended Data Table 1**) were used to amplify the plasmid to delete the branch-site containing sequence. A PCR product was purified by gel extraction (GeneJET Gel Extraction Kit, Thermo Scientific). Deleting the first 6 bp of the 5′-splicing site to make the 5′-SplicingΔ construct, was done by *in vitro* mutagenesis. All the plasmid constructs were confirmed by Sanger sequence analysis (Eurofins Genomics).
The pCMVΔ plasmid was made by deletion of the CMV promoter from the Sense construct digested by AseI and XhoI restriction enzymes (New England BioLabs). The single-stranded overhang from the digestion of the restriction enzymes was removed by Mung Bean Nuclease (New England BioLabs), then the construct was ligated by the T4 DNA ligase.

The plasmids transcribing single guide RNAs (sgRNAs) were constructed following the method presented in Zhang *et al*, Methods 2014[2]. The backbone plasmid was provided by Dr. M. Jasin's lab (Developmental Biology Program, Memorial Sloan-Kettering Cancer Center). The in-fusion reaction was used to replace the target sequence. The plasmid expressing Cas9 nuclease was purchased from Addgene (Plasmid #41815).

##### ***In-vitro* digestion assay with CRISPR-Cas9 system**

The sense and antisense plasmids isolated by Midiprep (GeneJET Plasmid Midiprep Kit, Thermo Scientific) were used for the in-vitro digestion assay to test the DSB efficiencies of each sgRNAs in the different constructs. The sense and antisense constructs were PCR-amplified using DsRed.pCis.F8 and

DsRed.pCis.R3 primers (**Supplementary Table 1**) to amplify a region near the DSB sites on the *DsRed* gene (**Extended Data Figure 4**). The PCR products were purified by using a 0.8x concentration of magnetic beads (HighPrep PCR Clean-up System, MagBio Genomics) to remove primer dimers. A DSB was induced on the PCR products by using sgRNA A, B or A and B (**Supplementary Table 2**) synthesized from Integrated DNA Technologies (Alt-R CRISPR-Cas9 sgRNA, IDT) with Cas9 nuclease (*S. pyogenes*, New England BioLabs). The total volume for each assay was 30 μl in a 1.5 ml RNase-free Microfuge tube (Applied Biosystem), which consisted of 30 nM of sgRNA, 30 nM of Cas9 nuclease and 3 nM of plasmid in 1x NEBuffer r3.1 (New England BioLabs). To determine the DSB efficiency by Cas9 with sgRNA A, B, or A and B, 30 nM of each sgRNA, 30 nM of Cas9 nuclease and 3 μl of 10x NEBuffer r3.1 with nuclease-free water for a total volume 29 μl were pre-incubated for 10 minutes at room temperature; then 1 μl of 90 nM PCR construct (from the Sense/pCMVΔ, BranchΔ, Antisense or 5′-SplicingΔ was added into the tube and incubated at 37 °C for 1 hour, then 1 μl of Proteinase K (Qiagen) was added into the tube the sample was incubated at 37 °C for 10 minutes to degrade the Cas9 nuclease to stop the cleavage. The sizes and molarity of digested fragments were determined by Bioanalyzer High Sensitivity DNA analysis (Agilent).

**Human cell lines and transfection**

Human embryonic kidney T (HEK-293T) *RNASEH2A* wild-type and KO cells were provided by Dr. Pursell's lab at Tulane University. The cells were grown in Dulbecco's modification of Eagle's medium (DMEM) containing 4.5 g/L glucose, L-glutamine and sodium pyruvate (Corning) with 10% fetal bovine serum (Sigma-Aldrich) and 1x penicillin-streptomycin (Gibco). Cells were grown at 37 ° C in a 5% $CO_2$ humified incubator. For the construction of the RNase H2A KO clones, *RNASEH2A* (Chr19, exon 2) gRNA (5'-TAACAGATGGCGTAGACCAT-3') was cloned into GeneArtTM CRISPR Nuclease Vector with OFP reporter (Invitrogen) following manufacturer's protocol. HEK-293T cells were transfected with 6 mL of Lipofectamine 2000 (Invitrogen) and 2.5 mg of gRNA vector DNA when the cells were 60-70% confluent. After 48 hours, OFP-positive cells were FACs sorted and serially diluted into 96-well plates at ~1.5 cells per well and incubated for 10-14 days. Single, well-defined clones were expanded to 24-well plates and initially screened with T7 Endonuclease I assay (NEB). KO clones were verified via western blot (α-RNASEH2A Ab Origene Cat.# TA306706) and Sanger sequencing of PCR products (TOPO Cloning Kit Invitrogen). The *RNASEH2A* locus PCR primers used to identify the mutations in the *RNASEH2A* alleles are shown in **Supplementary Table 1**. We identified a clone, HEK-293T T3-8, having three distinct frameshift mutations consistent with all three alleles modified in hypotriploid 293T cells (**Extended Data Figure 4a**). Modification of each allele was determined by TOPO cloning the PCR product (TOPO TA Cloning Kit Invitrogen) into DH5α competent cells (ThermoFisher). Sanger sequencing of the amplified PCR product showed the following observed alterations (all positions indicated with respect to the Cas9 cleavage site on the reference strand): insertion of G at position -1; deletion of five bases at position +2 to +6; complex alteration with deletion of three bases from position -1 to +2, deletion of 2 two bases from position +5 to +6, and CC>TT at position +8 and +9. HEK-293T T3-8 cells were used in this study as RNase H2A KO cells.

For transfection experiments, cells were seeded in 24-well plates at a density of ~50,000 cells per well for transfection and incubated for a day at 37 °C in a 5% $CO_2$ humified incubator. Cells were transfected using polyethylenimine (PEI, Polysciences), 5 μg were used per well for transfection. In all transfections, plasmids were used in the amount of 0.4 μg each, with a total amount of DNA not exceeding 2 μg per well. Cells were incubated for 6 days, to allow for more constructs to be cut and repaired, at 37 °C in a 5% $CO_2$ humified incubator after the transfection and used for the following experiments.

**Assays to test cell extract activity to cleave at an rGMP in DNA**

A mixture of 2.5 pmol of Cy5 5′-labeled oligonucleotide containing an rGMP (Cy5.3PS.rG, **Supplementary Table 1**) or the control DNA oligonucleotide (Cy5.3PS, **Supplementary Table 1**) and 3.75 pmol of complementary oligonucleotide (DNA.comp.3PS, **Supplementary Table 1**) in 1X Thermopol Reaction Buffer was heat denatured in boiling water for 5 min and cool down slowly to 30 °C to anneal the complementary oligonucleotides. 2.5 pmol of annealed oligonucleotide were incubated with 50 ng of protein extract from HEK-293T RNase H2A wild-type or KO cells prepared by using NP-40 lysis buffer (Alfa Aesar), or just water as the negative control, for 4 hours at room temperature. Successively, the mixture was treated with 9.95 ul of formamide (VWR, 0606-100ML) and incubated for 5 min at 95 °C to denature the double strand substrate, then the mixture was put on ice. The denatured substrate was mixed with 2.22 ul of 10X Orange Loading Dye (LI-COR, C80809-01) and loaded on a 15% 7M urea denaturing polyacrylamide gel. As a positive control for this experiment, we used 5 units of *Escherichia coli* RNase HII (NEB, M0288L) in place of the protein extract.

**Quantitative real-time PCR (qRT-PCR)**

RNA was isolated from the transfected cells after 6-days using the RNeasy Mini kit (Qiagen) and treatment by DNase (RNase-Free DNase set, Qiagen). RNA was converted into cDNA using QuantiTect Reverse Transcription kit (Qiagen). Customized TaqMan Assay containing primers and probe (Catalog# 4331348, Assay ID APU67JA, DSRED1, Applied Biosystem, **Supplementary Table 1**) and TaqMan Fast Advanced Master Mix (no UNG, Applied Biosystem) were used for analyzing RNA expression in 96-well plates (Applied Biosystem). The total volume in each well was 20 μl, which consisted of 10 μl of TaqMan Fast

Advanced Master Mix, 1 μl of TaqMan Assay (20x) containing primers and probe, 8 μl of nuclease-free water and 1 μl of cDNA. The cDNA level was determined using an ABI Prism 7000 RT-PCR machine (Applied Biosystem). Human GAPD (GAPDH) Endogenous Control (FAM/MGB probe, non-primer limited, Applied Biosystem), which is TaqMan primers and probe, was used for normalization. Values for each sample were normalized by GAPDH (ΔCt), and then each ΔCt value was normalized by ΔCt from the control sample transcribing the spliced transcript (ΔΔCt).

**Next-Generation Sequencing (NGS) library preparation**

Genomic DNA and plasmids were isolated using DNeasy Blood & Tissue kit (Qiagen) from transfected cells after 6 days of incubation at 37 °C in a 5% $CO_2$ humified incubator. The isolated DNA was amplified by a 1st PCR (20x cycles) using F primer 1 (YJo11-14, **Supplementary Table 1**) and R primer 1 (YJo15-26, **Supplementary Table 1**) having 6 nucleotides barcode (NNXXXX), where N is any nucleotide, XXXX is a specific barcode identifying each sample, and a part of adaptor sequences for Illumina sequencing. The 1st PCR products were amplified in a 2nd PCR (15x cycles) using F primer 2 (D501-508, **Supplementary Table 1**) and R primer 2 (D701, 702, 705, 707 and 712, **Supplementary Table 1**) containing an index and the adaptor sequences. The 2nd PCR products were purified by spin column (QIAquick PCR Purification Kit, Qiagen) and then run in the BluePippin (2% agarose DNA size selection cassette, Sage Science) at the Molecular Evolution Core Facility at Georgia Tech to remove the primer dimers. The extraction range was 130-500 bp for the BluePippin. Bioanalyzer High Sensitivity DNA analysis (Agilent) was used to check the sizes of the final sequencing libraries. We used Illumina HiSeq 2x150 for the NGS provided by Admera Health (Biopharma Services, South Plainfield, NJ). NGS libraries for the Antisense and 5′-SplicingΔ constructs were prepared as follows. The isolated DNA was amplified by using DsRed.pCis.F3 and DsRed.pCis.R3 primers (**Supplementary Table 1**) for the 1st PCR (20x cycles), then the amplified products were amplified by using F primer 2 (YJo1,9,10, **Supplementary Table 1**) and R primer 2 (YJo4-6,8,11, **Supplementary Table 1**) for the 2nd PCR. The extracted DNA of the Antisense and 5′-SplicingΔ constructs was used to construct sequencing libraries twice to increase the size of the libraries, using PCR primers with and without a barcode sequence, as described above. Thus, the DNA extracts for the Antisense and 5′-SplicingΔ constructs were sequenced twice.

To construct RNA sequencing libraries, RNA was isolated using the RNeasy Mini kit (Qiagen) from the plasmid-transfected cells after 6 days at 37 °C in a 5% $CO_2$ humified incubator. The isolated RNA was converted to cDNA using the QuantiTect Reverse Transcription kit (Qiagen). The cDNA was PCR amplified by the 1st PCR primers (YJo11-26) by 20x cycles, then the 1st PCR product was amplified by the

2nd PCR primers (D501-508 and D701, 702, 705, 707 and 712) by 15x cycles. All the PCR amplification was done by Q5 High-Fidelity DNA polymerase (New England BioLabs).

**Trimming of NGS data**

The reads in the sequencing libraries were tagged by a 6-nucleotides barcode NNXXXX. NN is any dinucleotide, and XXXX represents a specific 4-nucleotide barcode identifying each library. The barcode sequences were located at the beginning and the end of the reads. First, the reads were trimmed based on sequencing quality and Illumina adaptor sequence using cutadapt 2.10 with the default setting. After trimming by cutadapt, to trim the reads with the barcodes, the first 6 nucleotides of the reads (TTCAAG) following the Illumina adaptor and barcode sequences were searched in the first 12 nucleotides of the reads. If these sequences were found, we checked whether 4 nucleotides located prior to the TTCAAG sequence matched with the assigned barcode sequence. To check the barcode located at the end of sequencing reads, the length of the reads was measured first. If the reads were longer than 140 nucleotides, the barcode at the end of the reads could not be captured due to the maximum length of reads (150 nucleotides) by the Illumina HiSeq 150x2. If the length of sequencing reads was shorter than 140 bp, the last 6 nucleotides of the read sequence (ACTTCC) prior to the barcode were searched within the last 12 nucleotides of the reads. If that sequence was found, we checked whether 4 nucleotides located downstream of the ACTTCC sequence matched with the assigned barcode sequence.

Trimming for the Antisense and 5′-SplicingΔ NGS libraries that did not have the barcode sequence was only done by cutadapt 2.10 with the default setting. The FASTQ files for the two sequencing libraries derived from the same DNA source of each Antisense and 5′-SplicingΔ sample were concatenated to generate one FASTQ file for each sample.

**RNA sequencing data analysis**

The trimmed FASTQ files were aligned to the reference sequence of the corresponding construct using hisat2 2.2.1 with the default setting. The SAM files generated by hisat2 were used to categorize the sequencing reads as ‘spliced transcript’, ‘non-spliced transcript’, ‘alt-spliced transcript’, ‘non-canonical alt-spliced transcript’, and ‘unaligned’. Using the SAM file, the occurrence of splicing was checked first by sequence alignment. The reads without any splicing event were counted as non-spliced transcripts, and the reads having a splicing event with the same length as the intron were counted as spliced transcripts. Splicing on other sites starting with the dinucleotide GT and ending with the dinucleotide AG were counted as alt-spliced transcripts. The other sequencing reads having splicing that did not start with GT- and end with -AG were counted as non-canonical alt-spliced transcripts. All the other sequencing reads not aligned to the reference sequence by hisat2 were counted as unaligned.

**Calculating the frequency of RNA-templated DNA DSB repair**

We consider RNA-templated DNA DSB repair to be the mechanism that produces an identical sequence with the template RNA (except for Ts in place of Us). To calculate the frequency of RNA-templated repair, reads corresponding to the spliced transcript sequence were searched in each sequencing library to find those that are identical to the transcript RNA. To calculate the frequency, the number of reads containing this sequence was divided by the total read count.

**Categorizing sequencing reads to three groups: sequence with intron, DSB or gap repair via NHEJ, and DSB or gap repair via MMEJ**

Illumina HiSeq 2x150 was used for the NGS in which the sequencing reads have a maximum length of 150 bp. The fragments in the NGS libraries from the sense constructs are 229 bp with the intron and 118 bp without the intron if there was no mutation. The fragments in the BranchΔ library are 174 bp with the intron and 118 bp without the intron. If the sequencing reads contain the intron, the length of the reads reaches the maximum length of the Illumina reads, but if not, the length of the sequencing reads is around 118 bp with variations at the DSB site caused by the repair. Therefore, to determine whether the reads contain the intron or not, the reads were filtered by length. First, all the reads that were longer than 130 bp were classified as sequences with intron. The reads shorter than 130 bp were classified as DSB/gap repair via NHEJ or MMEJ by checking whether the reads were the products of MMEJ. The products of MMEJ were classified as DSB or gap repair via MMEJ, and all the other reads shorter than 130 bp were classified as DSB or gap repair via NHEJ.

**Identification of microhomology pairs and calculation of MMEJ frequency**

Microhomology pairs were defined as two identical DNA segments on the reference sequence that were at least 3 bp long. We identified all microhomology pairs that had one segment upstream and another downstream of the DSB and were between the primer sequences. For the sense constructs, we excluded from the analysis pairs whose segments overlapped the branch site (55 bp) since that region would not be present in the BranchΔ reference sequence. For the antisense constructs, we excluded from the analysis pairs whose segments overlapped the 5′-splice site since that region would not be present in the 5′-SplicingΔ reference sequence. We also only considered microhomology pairs whose matching segments were maximal (i.e., could not be extended to a pair of longer matching segments). The reference sequence between the primers was subdivided into 3 regions (Exon1, Exon2, and Intron) and the microhomology pairs were categorized into 4 groups (Exon1-Exon2, Exon1-Intron, Exon2-Exon1, and Exon2-Intron) depending on which regions their segments were located and where the DSB induced. A name was assigned to each microhomology pair such that corresponding microhomology pairs in the different constructs (e.g., Sense, BranchΔ, and pCMVΔ, or Antisense and 5′−SplicingΔ) had the same name. See **Extended Data Figures 9b, c** and **11b, c** for the locations and nucleotide sequences of the microhomology pairs and the regions Exon1, Exon2, and Intron on the reference sequence. In this analysis, DSB repair by MMEJ was detected in a read contained a microhomology deletion signature. A microhomology deletion signature was defined as the DNA sequence that results by taking the reference sequence, deleting the sequence between the pair of identical segments (keeping only one copy of the segment), and extracting the 10-bp flanks plus the segment in the middle. We detected all MMEJ events from the raw reads and calculated their frequency, which is the count of MMEJ events divided by the total read count. The corresponding python3 scripts to calculate the MMEJ frequency are available on GitHub.

**Bar plots and box plots for MMEJ frequencies**

Bar plots and box plots were generated to compare the MMEJ frequencies between different constructs of the sense or antisense systems. In the bar plots, the mean frequency of samples is shown, and the standard deviation is indicated by whiskers. In the box plots, the median of the points is shown as the middle line of the box. The first and third quartiles are indicated by the box frames and the whiskers represent the largest point not more than 1.5 interquartile range (IQR) beyond the box frame. All data points outside the whiskers are classified as outliers and shown as diamond points. Two-tailed Mann-Whitney *U* tests were performed to compare the frequencies of the different constructs.

**Calculating the frequency of flipped-intron insertion following 2 DSBs**

The frequency of flipped-intron insertion was determined by detecting 20 bp of the complementary sequence of the intron sequence specifically located 10 bp away from the DSB site so that is could be captured within the sequencing reads. The antisense constructs have a sequence that is different from the sequence obtained by flipping the intron in the sense constructs (**Extended Data Figure 1**); thus, reads of the antisense constructs cannot be confused for reads of the flipped intron of the sense constructs.

**Statistical analysis of data**

Two-tailed Mann-Whitney *U* test was used for statistical analysis of sequencing data. Moreover, permutation tests were performed to compare the ratio of Sense/BranchΔ frequency between RNase H2A wild-type and KO libraries for each of the repair mechanisms: NHEJ, R-TDR, and each pair of MMEJ. For each repair mechanism, we calculated the frequency ratio, $R_{WT}$ of Sense/BranchΔ as the average repair frequency of the four wild-type, Sense repeats divided by the average repair frequency of the four wild-type, BranchΔ repeats. Similarly, we calculated the frequency ratio, $R_{KO}$, for the KO experiments. Then we calculated the observed test statistic as $T = R_{KO}/R_{WT}$. With the null hypothesis that the wild-type and KO frequencies were exchangeable within the same construct, we calculated the exact permutation test null distribution of $T$, denoted $T^*$. We did so by permuting the four wild-type, Sense frequencies with the four KO, Sense frequencies, and the four wild-type, BranchΔ frequencies with the four KO, BranchΔ frequencies in all possible ways, and then calculating $T^*$ using the same procedure defined for $T$. The $P$ value was obtained as the number of $T^*$ values greater (respectively less) than $T$, for the one-sided alternative hypothesis $R_{KO} > R_{WT}$ (respectively $R_{KO} < R_{WT}$). The corresponding scripts are available on GitHub.

**Bioinformatics processing to obtain variation-distance graphs to study RNA-mediated DSB repair by NHEJ**

Each sequence read in each NGS library was aligned to the corresponding reference sequence, that is, the sequence of the construct between the two sequencing primers. We used the Bowtie2 2.4.1 software with the default setting to generate SAM files. We discarded reads that Bowtie2 failed to align and those that did align but not with the first base pair of the reference sequence. On each SAM file obtained from Bowtie2, we performed the following data processing: (i) We discarded short reads that were less than the minimum expected length (130 bp for one DSB system and 50 bp for the gap system). To analyze the sequence variations at the DSB site, we used a custom script that realigned each read to the reference sequence so that the in/dels encapsulate the DSB position. If this alignment had the same or lower number of mismatches than the original alignment, this new alignment was kept and replaced the original one. We kept only the reads which, after the previous steps, resulted in an alignment such that in/dels were both contiguous and encapsulated the DSB position. For the one-DSB system, the libraries contained at least 93% of sequences after filtering; for the gap system, libraries varied between 14% and 50%. (ii) To concentrate on variations near the break site, we extracted a *DSB-sequence window* from each aligned sequence corresponding to the nucleotides that align with positions -10 to +10 relative to the DSB site on the reference sequence. To ensure the window reflected only variations near the DSB, we additionally required it to be flanked by two anchor sequences. The left anchor sequence corresponds to nucleotides -30 to -11 relative to the DSB site on the reference sequence, and the right anchor sequence corresponds to nucleotides 11 to 30 relative to the DSB site on the reference sequence (**Extended Data Figure 1**). We discarded sequences whose left or right anchor sequences had alignments with ≥ 2 mismatches or ≥ 1 in/del. Because substitutions are common NGS sequence errors[3], and in fact, substitutions were abundant in the negative control samples with no-DSB (**Extended Data Figure 5**), we only considered in/dels in the DSB-sequence windows in the NHEJ analysis. To do so we replaced all alignment substitutions with the corresponding nucleotide of the reference sequence. Then the sequence reads with the same DSB-sequence window were combined by summing their frequencies. Since the Antisense and 5′-SplicingΔ libraries were sequenced twice, we merged the two resulting outputs of the previous steps and recomputed their frequency accordingly. The DSB-sequence windows that had a frequency $\leq 10^{-5}$ in at least one of the repeats were not used in the analysis. This process of windows extraction discarded at most an additional 3.8% of each library after step (i). For each experiment, we obtained a table of DSB-sequence windows by assigning each DSB-sequence window the average frequency of all repeats of the same experiment.

**Data availability**

The authors declare that the data supporting the findings of this study are available within the paper and its supplementary information files. The NGS datasets generated during the current study are available in NCBI's Sequence Read Archive via BioProject "PRJNA883674". All data are available from the authors upon reasonable request.

**Code availability**

Customized Python3 scripts for all NGS data analysis in this study are available on GitHub under GPLv3.0 license (https://github.com/xph9876/RNA-mediated_DSB_repair).

**Reference**


1 Zheng, C. & Baum, B. J. All human EF1alpha promoters are not equal: markedly affect gene expression in constructs from different sources. *Int J Med Sci* **11**, 404-408 (2014).

2 Zhang, Y., Vanoli, F., LaRocque, J. R., Krawczyk, P. M. & Jasin, M. Biallelic targeting of expressed genes in mouse embryonic stem cells using the Cas9 system. *Methods* **69**, 171-178 (2014).

3 Chen, L., Liu, P., Evans, T. C., Jr. & Ettwiller, L. M. DNA damage is a pervasive cause of sequencing errors, directly confounding variant identification. *Science* **355**, 752-756 (2017).
